# Physiological perfusion of human vasculature reveals a YAP/TAZ-Apelin switch linking intraluminal flow to endothelial state transitions and vessel remodeling

**DOI:** 10.64898/2026.03.21.713033

**Authors:** Tiger H.Z. Jian, Adam A. Sivitilli, Yaxin E. Guo, Callum J. Stirton, Jessica T. Gosio, Yuko Tsukahara, Johnny M. Tkach, Suying Lu, Amin Yarmand, Maria Mangos, Rod Bremner, Jeffrey L. Wrana, Liliana Attisano, Laurence Pelletier

**Affiliations:** Lunenfeld-Tanenbaum Research Institute, Mount Sinai Hospital; Toronto, Ontario, Canada; Department of Molecular Genetics, University of Toronto; Toronto, Ontario, Canada; Donnelly Centre, University of Toronto; Toronto, Ontario, Canada; Department of Biochemistry, University of Toronto; Toronto, Ontario, Canada; Department of Ophthalmology and Visual Science, University of Toronto; Toronto, Ontario, Canada; Department of Laboratory Medicine and Pathobiology, University of Toronto; Toronto, Ontario, Canada

## Abstract

Vascular flow delivers nutrients and imposes hemodynamic forces that govern vessel behavior in health and disease, yet fully human systems that recapitulate and tune physiological intraluminal flow in three-dimensional (3D) tissues are lacking. We developed VIVOS (<u>V</u>ascularized *In <u>V</u>itro* <u>O</u>rgan <u>S</u>ystems), a platform that couples perfused human vascular beds to tunable pumps, generating continuous intraluminal flow through millimetre-scale vessels and 3D tissues at physiological shear stresses and pressures. VIVOS supports integration and perfusion of diverse human organoids and tissues, including lung organoids, cerebral organoids, vascular organoids, breast spheroids, and human retinal explants, as well as enables direct measurement and control of pressure, shear stress, and perfusion-dominant compound transport over extended culture periods. By tuning intraluminal flow and applying single-cell transcriptomics, we uncover a remodeling program in which laminar shear stress acts through a YAP/TAZ-TEAD “switch” to rewire an Apelin ligand-receptor axis and bias tip-stalk endothelial states, reshaping human vascular networks and linking hemodynamic cues to cell state transitions. We further model fast-flow arteriovenous malformations (AVMs) from Hereditary Hemorrhagic Telangiectasia and show that BMP9 constrains vessel caliber and perfusion while antagonizing a VEGF-driven angiogenic program, generating flow-quantified AVM-like lesions in a fully human 3D context. Together, these findings establish VIVOS as a generalizable platform that links physiological intraluminal flow to endothelial state transitions and vessel remodeling, enabling preclinical testing and mechanistic dissection of flow-regulated vascular pathologies in perfused 3D human tissues under defined hemodynamic conditions.

## Introduction

Organ chips and organoids are *in vitro* methods designed to overcome the tissue complexity and species specificity limitations of traditional preclinical models ^1,2^. However, these methods differ fundamentally from real organs in how nutrients and other bioactive compounds move around in 3D space. Whereas real organs rely on intraluminal vascular flow to distribute nutrients and signaling molecules, support organ-specific endothelial–tissue communication, and enable key functional structures such as the lung alveolus and blood–brain barrier ^3,4^, organ chips and organoids predominantly rely on non-physiological diffusion or non-vessel flow ^1,2,5,6^. Earlier *in vitro* vascularization strategies have focused on either stem-cell-derived vasculature ^7,8^, direct incorporation of endothelial cells into organoids ^9,10^, or self-assembled vasculature on small-scale microfluidic devices using gravity-driven or pump-perfused intraluminal flow ^11–14^. While these studies are promising, with some demonstrating enhanced organoid maturation ^8,9^, the vessels lack continuous, physiologically relevant flow or remain confined to small microvascular beds, limiting perfusion of larger human tissues. To circumvent this problem, organoids have been grafted into mice to hijack their circulatory systems and use their hearts to pump flow into vessels ^15^. However, this returns organoids back to an animal context and limits experimental control. The key challenge that remains is to build a scalable artificial circulatory system that reproduces physiological vascular flow in a fully human, animal-free setting.

Our knowledge of how vasculature develops, through angiogenesis and vasculogenesis, is primarily based on animal model studies which have implicated molecular components such as vascular endothelial growth factor (VEGF), Apelin, bone morphogenetic proteins (BMPs), and Hippo-YAP/TAZ ^16–19^. Blood vessels are continuously exposed to hemodynamic cues, particularly fluid shear stress on the vessel wall from vascular flow, and changes in the magnitude, direction, or type of flow are key inputs that shape vascular development and physiology ^20,21^. These cues dictate multiple facets of vessel behavior from endothelial proliferation and migration, to vascular inflammation and atherosclerosis progression. Despite the well-established importance of vascular flow, how hemodynamic forces integrate with molecular signaling pathways remains incompletely understood, particularly in human vasculature. Vascular flow cannot be easily manipulated within animal models and applying fluid shear stress to monolayer endothelial cells has limited physiological relevance. Human *in vitro* vasculature that supports tuning of physiologically relevant vascular flow would therefore be critical for mechanistically linking hemodynamic forces to endothelial cell signaling and regulation of vascular architecture. Vascular disease modeling is similarly based on animal model studies with conditions such as Hereditary Hemorrhagic Telangiectasia (HHT), a genetic vascular disorder characterized by life-threatening fast-flow arteriovenous malformations (AVMs) that cannot be faithfully studied using monolayer cultures ^22,23^. As in animal models of vascular development, animal models of vascular diseases are constrained by throughput and limited access to tools for precise molecular perturbations. Fully human *in vitro* models that recreate disease-relevant vasculature, like those seen in HHT or other vascular conditions, could provide new insights into disease mechanisms and support the development of therapeutic strategies.

Here, we introduce VIVOS (<u>V</u>ascularized *In <u>V</u>itro* <u>O</u>rgan <u>S</u>ystems), an electromechanical platform that generates physiological intraluminal flow *in vitro* by connecting millimetre-scale human vascular beds grown in microfluidic chips to tunable impeller pumps. We use VIVOS to vascularize diverse human organoids and tissues, quantify perfusion-dominant compound transport and shear stress in an all-human setting, and mechanistically link physiological intraluminal flow to endothelial state transitions through a YAP/TAZ-Apelin switch that remodels vascular networks. Finally, we model Hereditary Hemorrhagic Telangiectasia (HHT)-like arteriovenous malformations under defined hemodynamic conditions and show that BMP9 constrains vessel calibre and perfusion while antagonizing a VEGF-driven angiogenic program. Overall, VIVOS provides a fully human preclinical platform in which physiological intraluminal flow is a measurable and precisely controlled experimental variable for dissecting flow-regulated vascular pathologies and testing therapeutics.

## Results

### Electromechanical Perfusion of Living Blood Vessels

VIVOS (<u>V</u>ascularized *In <u>V</u>itro* <u>O</u>rgan <u>S</u>ystems) consists of vascular beds grown in microfluidic chips that are continuously perfused by mechanical impeller pumps (**Fig. 1A; Extended Data Fig. 1A**). The vascular beds, which can be implanted with organoids and/or other 3D tissues, are generated from human-derived primary endothelial (i.e. BMVECs: brain microvascular endothelial cells) and stromal (i.e. NHLFs: normal human lung fibroblasts and HBVPs: human brain vascular pericytes) cell lines suspended within a fibrin-based hydrogel (**Fig. 1B; Extended Data Fig. 2**). These endothelial cells self-assemble and form robust vascular networks within 4-8 days (**Extended Data Fig. 3A**) in a manner requiring the stromal cells (**Extended Data Fig. 3B**). Perfusion of media through the vascular bed is driven by connection of vessel lumens to microfluidic channels via porous membranes (**Fig. 1C**). The impeller pumps, magnetically coupled to spinning motors, provide continuous and electronically tunable fluid flow (**Fig. 1D-F**). Implanted organoids receive media flow from the perfused vasculature and integrate into the vascular network (**Fig. 1G-I**).

**Fig. 1.**
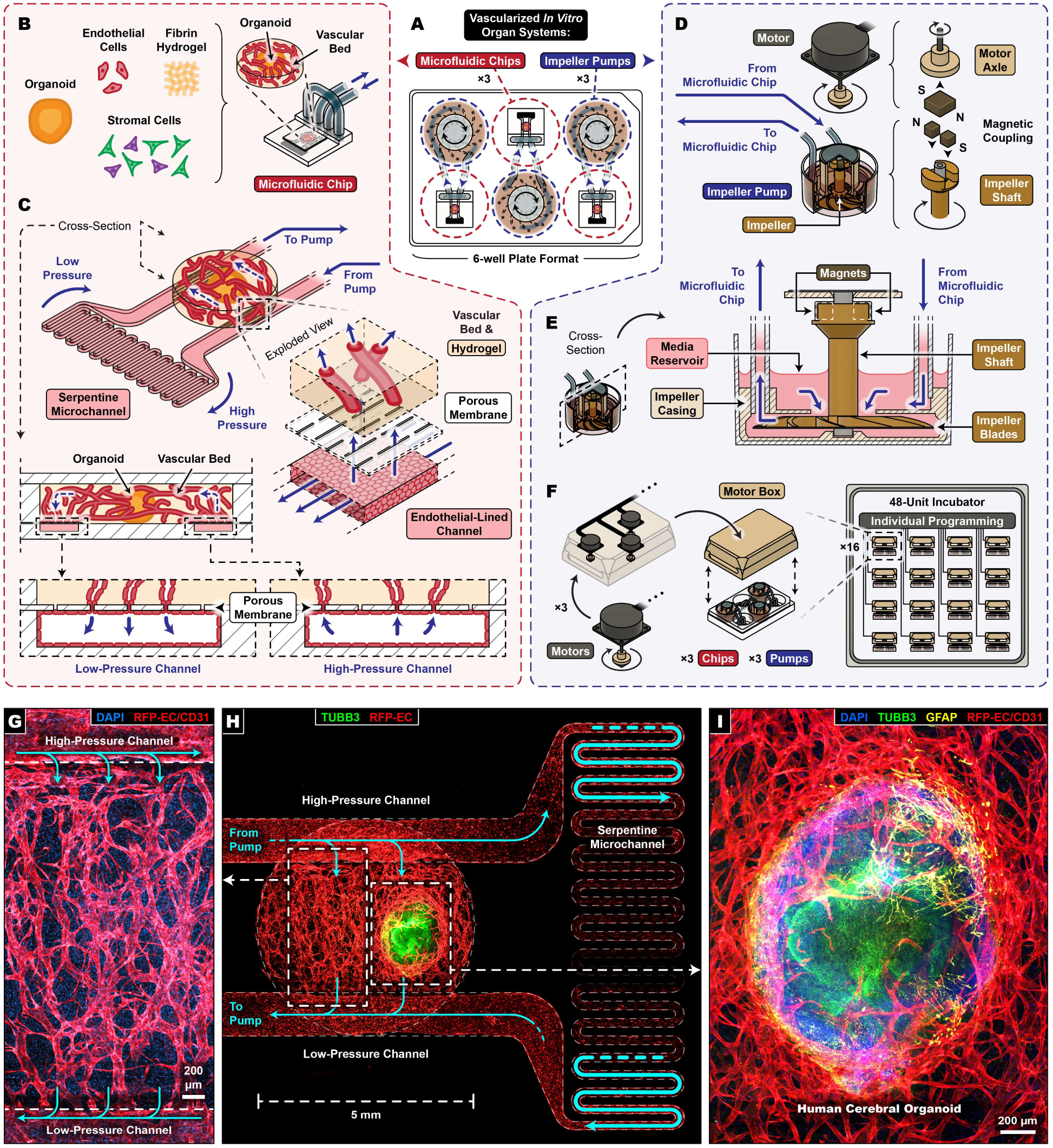
Electromechanical perfusion of *in vitro* vascular beds using VIVOS (<u>V</u>ascularized *<u>I</u>n <u>V</u>itro* <u>O</u>rgan <u>S</u>ystems). **(A)** 6-well plate containing 3 microfluidic chips connected to 3 impeller pumps. **(B)** Vascular beds, which may additionally include organoids, develop within microfluidic chips from endothelial and stromal cells embedded within a fibrin hydrogel. **(C)** Overview of the microfluidic chip (top), which features a serpentine microchannel separating two straight channels lined with endothelial cells, one at high pressure and one at low pressure, onto which the vascular bed sits. Exploded view (right) highlights interfacing of the vascular bed with the endothelial-lined channels through porous membranes. Cross-sectional view (bottom) highlights passage of fluid from the high-pressure channel to the low-pressure channel via the vascular bed and porous membrane interfaces. **(D)** The impeller pump utilizes magnetic coupling to spin impellers and create fluid flow. **(E)** Impeller pump cross-sectional view. **(F)** Overview of the motor box, which sits on top of the 6-well plates to spin the impeller pumps. **(G)** Enlarged image of the vascular bed spanning the high- and low-pressure channels. **(H)** Whole image of the microfluidic chip. **(G, H)** Cyan arrows depict fluid flow direction. **(I)** Enlarged image of a human cerebral organoid embedded within the vascular bed (TUBB3: neurons, GFAP: astrocytes). **(G, H, I)** Vessels were grown from RFP-labeled endothelial cells. **(G, I)** Imaging of RFP-labeled vessels was enhanced with a BiaPy RFP-to-CD31 deep learning model.

The microfluidic chip consists of a single path divided into two straight channels separated by a compact serpentine microchannel (**Fig. 1C; Extended Data Fig. 1A, B; Extended Data Fig. 4A**). Due to its high relative flow resistance, the serpentine microchannel imparts a pressure drop on incoming pressurized fluid from the impeller pump and leaves one straight channel at high pressure and the other straight channel at low pressure (**Extended Data Fig. 4B-E**). These high- and low-pressure channels are positioned beneath a disk-shaped “vascular bed compartment”. Interfacing the compartment with the two channels are porous membranes bearing 20-µm-wide slits (**Fig. 1C; Extended Data Fig. 1A, B**). Driven by the pressure differential, fluid from the high-pressure channel crosses the porous membrane, perfuses the vascular bed through the self-assembled vessel network, and exits to the low-pressure channel by crossing another porous membrane (**Fig. 1C; Extended Data Fig. 4D; Supplementary Video 1**). Individual vessels connect with the pores and merge with endothelial cells coating the channel walls to form a continuous surface bridging the vascular bed with the channel interiors (**Fig. 1C; Extended Data Fig. 3C-E**).

The choice of slits for the porous membrane is 2-fold. First, slits maximize surface tension across the pores to prevent cross-membrane seepage of unpolymerized hydrogel during the seeding process (**Extended Data Fig. 2**). Second, others have identified that sharp right angles within microfluidic devices lead to recirculation vortices and “disturbed” flow ^20,24,25^, which can increase cell death and lead to endothelial inflammation. To better mimic physiological vessels, where no sharp right angles are present, we chose long slits parallel to the direction of flow to minimize formation of these recirculation vortices.

By using rotating impellers to pressurize fluid (**Fig. 1D, E; Extended Data Fig. 5A, B**), the impeller pump provides many advantages over other pump types. Impellers generate a wide range of precise and reproducible flowrate and pressure values that are far greater in magnitude than those from gravity driven systems (**Extended Data Fig. 5C; Extended Data Fig. 8A**) ^13^. Flow from impellers does not have the intrinsic pulsations found in peristaltic or membrane type pumps (**Extended Data Fig. 5D**). Impellers, like the heart, approximate constant pressure across varying flow resistances, rather than the constant flowrates produced by syringe pumps (**Extended Data Fig. 5E**). Despite the impeller’s high speed of rotation, fluid shear stress values within the impeller casing do not exceed what occurs during physiological circulation (**Extended Data Fig. 5F**) ^26^. Combined with a secondary “bypass” tubing straddling the microfluidic chip, the impeller pump drives fluid recirculation and media mixing through the entire flow circuit on the order of seconds, which mimics the speed of blood recirculation (**Extended Data Fig. 5G**).

The impellers spin via magnetic coupling between magnets embedded within both the impeller shafts and the axles of spinning motors (**Fig. 1D; Extended Data Fig. 1C**). These motors permanently reside within the incubator in “motor boxes” containing three motors each (**Fig. 1F; Extended Data Fig. 1D-E**). Each motor box engages a standard 6-well plate containing three microfluidic chips connected to dedicated impeller pumps and media reservoirs, enabling scalable parallelization (**Fig. 1A, F; Extended Data Fig. 1E**). Because no moving parts are in direct contact with the plates, their routine removal is connector- and gasket-free and does not compromise sterility. This pump setup additionally enables precise, dynamic, and low latency flow control through closed-loop regulation of motor speeds using integrated rotation sensors (**Extended Data Fig. 5H, I; Supplementary Video 2**). Overall, this impeller pump system, combined with microfluidics, drives continuous perfusion through the self-assembled vascular beds.

### Physiological Intraluminal Flow on VIVOS Drives Perfusion-Dominant Compound Transport

Transport of nutrients and other bioactive compounds within tissues occurs through a mix of diffusion, avascular flow (i.e. interstitial flow), and intraluminal vascular flow ^27^. Pure diffusion is constrained by scaling limits, which restrict the size of avascular tissues in nature and in the lab. Large complex organisms such as humans therefore rely on vascular flow to sustain their size. While others have generated perfusable vasculature ^11–14^, notably with smaller devices, the physiological relevance of vascular flow in these devices is unclear. Physiological relevance in this context refers to vascular flow that is continuous, at sufficiently high levels, and the dominant mechanism by which compounds move around 3D tissues. Moreover, these smaller devices generally remain viable and still form vessels in the absence of flow ^11,14^, consistent with a scale where diffusion alone can provide sufficient nutrients to sustain the tissues.

To model compound transport on VIVOS, we generated empirical data and assessed whether compounds circulated primarily through diffusion, avascular flow or vascular flow. In this context, we defined compound transport as how compounds enter the interstitial space of the vascular bed from the microfluidic channels (**Fig. 2A-C**). Vascular flow involves compounds crossing vessel walls, while diffusion and avascular flow involve compounds leaking through gaps between endothelial cells at the porous membrane interface. We approximated total compound transport as the sum of these three compound transport mechanisms and measured each independently (**Fig. Extended Data 6A**). For standardization, we tracked the movement of 4 kDa FITC-labeled fluorescent dextran. We quantified diffusion and avascular flow by pulsing fluorescent dextran into the microfluidic channels of non-perfusable devices on Day 2 and tracking its spread into the vascular bed compartment with the impeller pump off (diffusion) or on (avascular flow) (**Extended Data Fig. 6B-H**). To estimate vascular delivery, we combined vessel flowrates with vessel permeabilities to capture both luminal transport and trans-vascular flux into the interstitial space (**Fig. 2D; Extended Data Fig. 7A**). Vessel permeabilities were measured by pulsing Day 4 vessels with fluorescent dextran, followed by time-lapse imaging to track the rate of dextran efflux (**Fig. 2D; Extended Data Fig. 7B-D; Supplementary Video 3**). Vessel flowrates were measured by combining tracking of fluorescent beads perfused through Day 4 vessels, fast time-lapse imaging, and particle image velocimetry (**Fig. 2D; Extended Data Fig. 7E; Supplementary Video 4**). This analysis also provides the vessel wall shear stresses used later in the study. Overall, comparing measurements from all analyses indicates that intraluminal vascular flow drives compound transport (**Fig. 2E**). This comparison shows that VIVOS delivers compounds to tissues predominantly via intraluminal vascular flow, distinguishing it from standard culture methods that rely largely on passive diffusion.

**Fig. 2.**
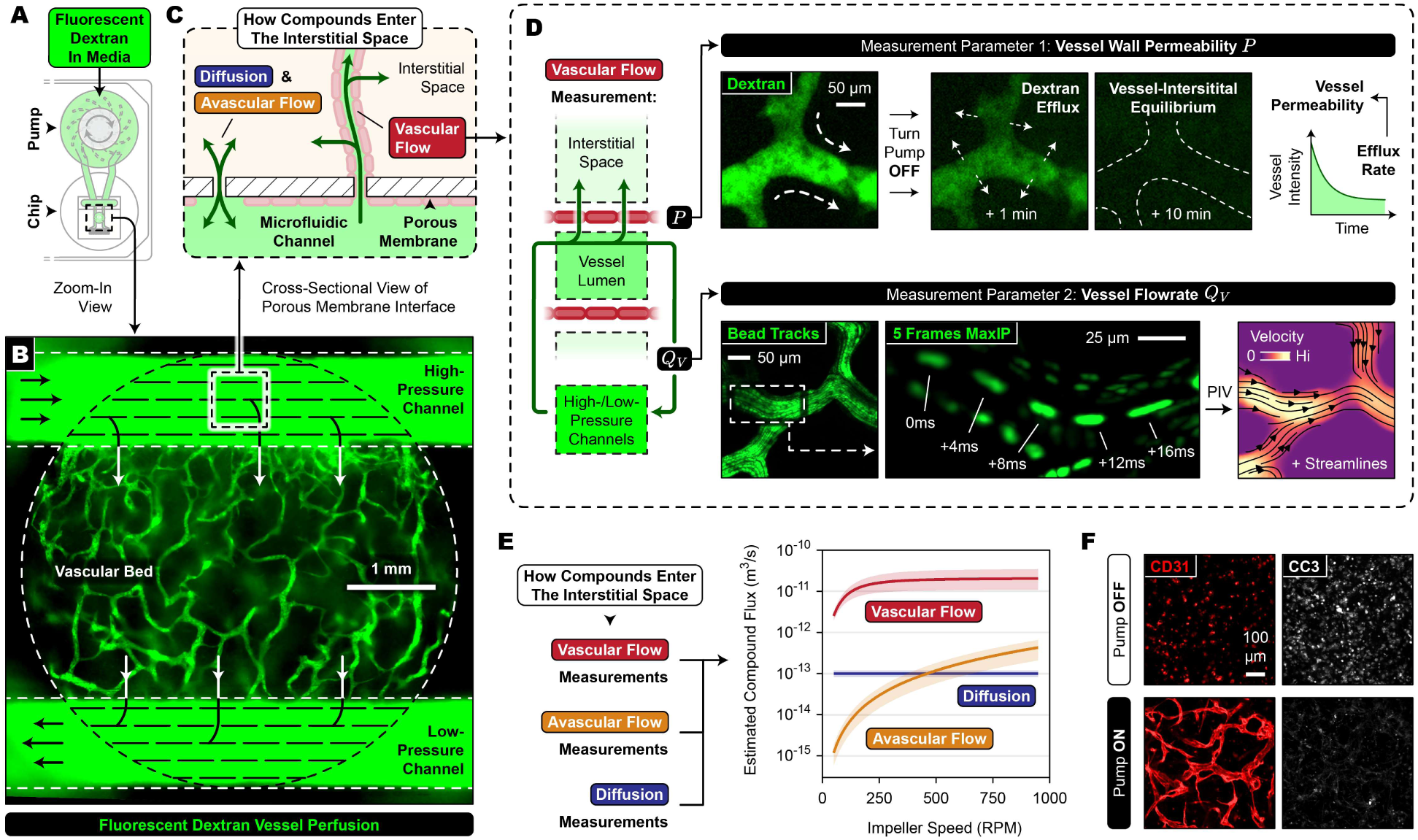
VIVOS employs vascular flow through continuously perfused vessels for compound transport. **(A)** 4 kDa FITC-labeled dextran is added to the impeller pump’s nutrient media reservoir and pumped through the microfluidic chip and vascular bed. **(B)** Image of a dextran-perfused vascular bed. **(C)** Dextran (or any compound) enters the interstitial space of the vascular bed through either diffusion, avascular flow, or vascular flow. Diffusion and avascular flow involve device imperfections where individual pores are not attached to vessels and/or breaks in the endothelial monolayer lining the channels. Vascular flow involves dextran entering lumens of endothelial vessels and crossing vessel walls into the interstitial space. **(D)** Two critical parameters are required for empirically measuring dextran transport into the interstitial space using vascular flow (left). Parameter 1 (top), vessel wall permeability, is measured by tracking the efflux rate of fluorescent dextran into the interstitial space from static vessels. Parameter 2 (bottom), vessel flowrate, is measured through fast time-lapse imaging of fluorescent beads continuously perfused through vessels combined with particle image velocimetry. **(E)** Comparison of empirical measurements for dextran entering the interstitial space. Measurements are expressed as total flux (i.e. molecules per second) divided by concentration in the incoming nutrient media. 90% confidence interval calculations are highlighted. **(F)** Vascular beds cultured for 4 days with the impeller pump either on or off.

Implanting organoids within vascular beds for modeling organoid-vessel interactions is challenging because the vascular bed’s size must match or exceed that of the organoid. As organoids can easily reach the millimeter scale, static vascular beds without continuous vascular flow will inevitably face nutrient limitations at those scales when passive diffusion becomes insufficient. This is apparent when VIVOS devices cultured without impeller-driven flow showed increased cleaved caspase-3 (CC3) levels and a complete failure to form vessels (**Fig. 2F**). Because VIVOS circulates compounds primarily by vascular flow when the impellers are spinning, we designed the vascular bed compartment to be multiple millimeters across and compatible in size with most current organoid varieties (**Fig. 3A**). To demonstrate this versatility, we implanted a wide range of tissues into VIVOS vascular beds including human-derived breast spheroids, vascular organoids, cerebral organoids, retinal explants, and lung organoids (**Fig. 3B**). Closer examination of implanted tissues showed multiple forms of engraftment and integration into the vascular networks. Cerebral organoids formed structures resembling neurovascular units with neurons, astrocytes, and endothelial cells in tight association (**Fig. 3C-E**). Cells of the spheroids extended outwards to intermingle with vessels (**Fig. 3F**). Vascular organoids integrated with the host vasculature and stromal compartment (**Fig. 3G**), and lung organoids had vessels beside the lung epithelium (**Fig. 3H**). By leveraging the wide range of organoid types, VIVOS enables modeling of vascular interfaces from a wide range of different organs with perfusable vasculature.

**Fig. 3.**
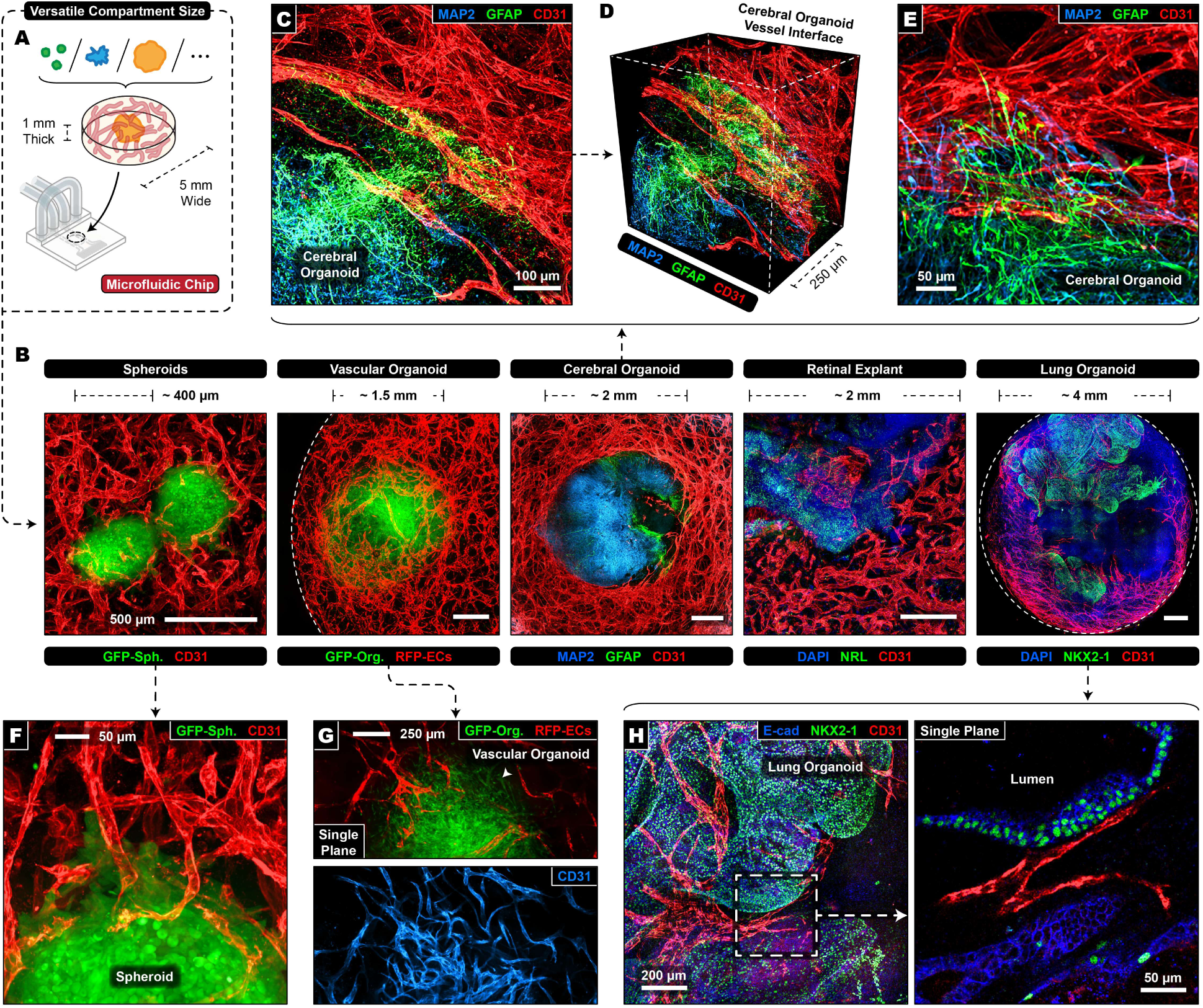
Large vascular beds on VIVOS enable versatile organoid & tissue vascularization. **(A)** Vascular beds on VIVOS are disks which measure 1 mm thick and 5 mm wide. **(B)** Organoids and 3D tissues from a variety of shapes and sizes are implanted within vascular beds on VIVOS. **(C)** Image of a human cerebral organoid implanted onto VIVOS. **(D)** 3D rendering of the cerebral organoid from (C). **(E)** Zoom-in on the interface between the vascular bed and the cerebral organoid from (C). **(F)** Image of a MCF10A spheroid implanted on VIVOS. **(G)** Image of a human vascular organoid implanted on VIVOS. **(H)** Image of a human lung organoid implanted on VIVOS. **(B-H)** CD31: vessels, MAP2: neurons, GFAP: astrocytes, NRL: rod photoreceptors, NKX2-1: lung progenitors, E-cad: lung epithelium.

### Flow-Dependent Reshaping of Transcriptional Programs and Endothelial State Transitions in the Vascular bed

To investigate how flow modulates vascular remodeling, we next grew vascular beds using “low flow” (300 RPM; ∼ 65 Pa pressure differential) or “high flow” (900 RPM; ∼ 610 Pa pressure differential) (**Fig. 4A, B; Extended Data Fig. 8A; Supplementary Video 5**). These conditions span non-physiological and physiological ranges of intraluminal shear stress and provide a controlled setting for asking how flow reprograms endothelial cell states. High flow generated vessel wall shear stresses of ∼ 2-3 dyn/cm^2^ (**Fig. 4C-F**) that was within the physiological range ^26,28^ and increased immunostaining for KLF4, a known shear stress induced factor (**Extended Data Fig. 8B, C**). Low flow vascular beds exhibited a denser, plexus-like network, whereas high flow produced a more tubular, pruned vascular architecture (**Fig. 4B**). We performed scRNA-seq of vascular beds one day after seeding (Day 1) to capture early vessel formation, and on Day 4 using either non-physiological low flow or physiological high flow (**Fig. 4G**). Vascular beds were generated using three primary cell lines (i.e. BMVECs: brain microvascular endothelial cells; NHLFs: normal human lung fibroblasts; HBVPs: human brain vascular pericytes). Single nucleotide variant (SNV) demultiplexing was used to track cell line origins (**Extended Data Fig. 9A, B**) and unsupervised clustering was applied to identify major cell types that were conserved across experimental conditions. (**Fig. 4G, H; Extended Data Fig. 9C**).

**Fig. 4.**
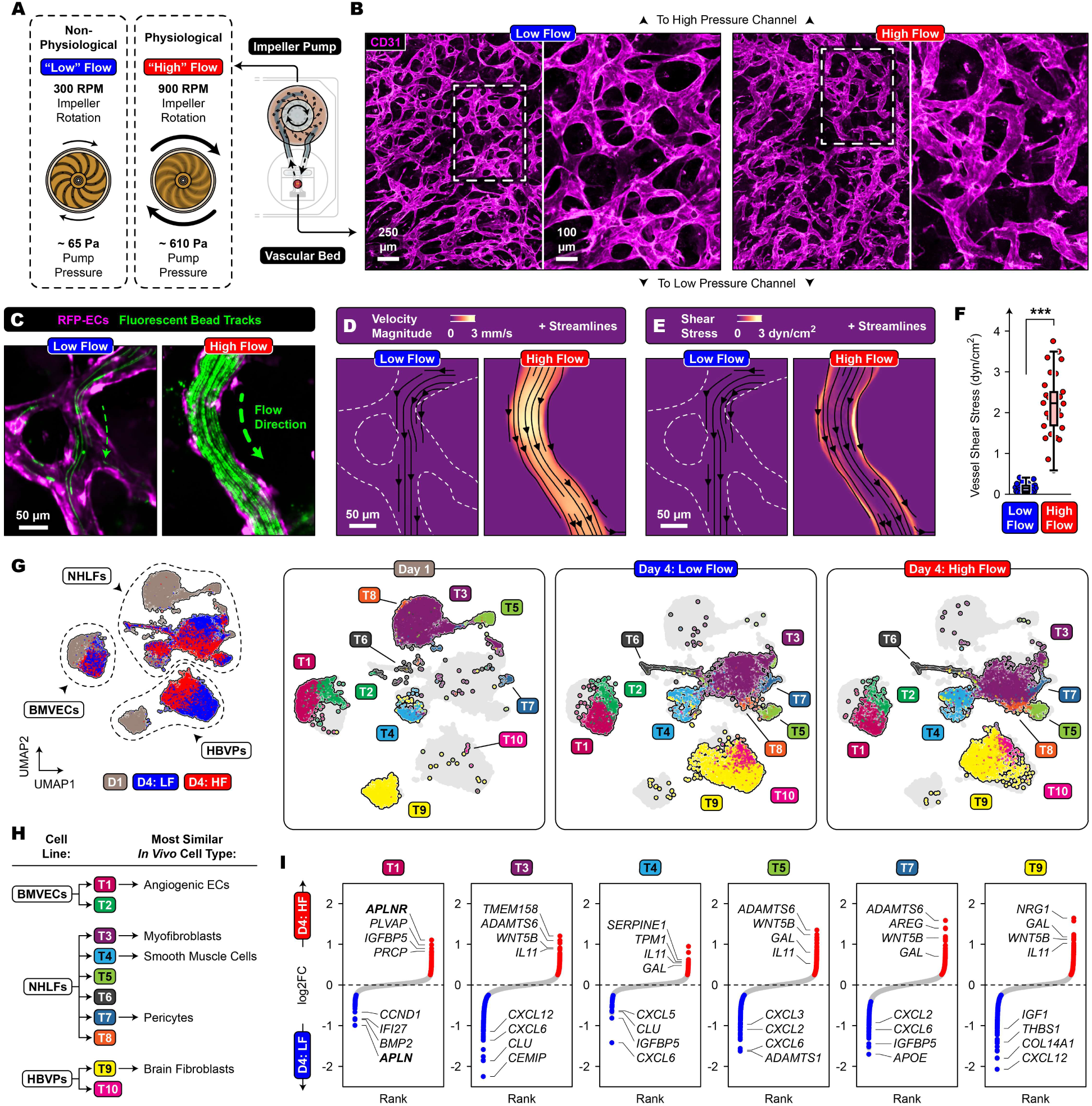
Tuning the impeller pump of VIVOS to investigate biological mechanisms of vascular flow. **(A)** Experimental setup involving setting the impeller pump’s rotation at either a non-physiological “low flow” (300 RPM) or a physiological “high flow” (900 RPM). **(B)** Images of vascular beds cultured using either low or high flow on Day 4 (CD31: vessels). **(C)** Perfusion of 1 µm diameter fluorescent beads through RFP-labeled vessels on Day 4 using either low or high flow. **(D, E)** Particle image velocimetry analysis of (C) showing velocity magnitude and shear stress. **(F)** Comparison of shear stress values from vessels grown and perfused using either low or high flow (28-30 vessels per condition across 3 devices; 2-tailed unpaired Welch’s t-test; *** p < 0.001). **(G)** Uniform Manifold Approximation and Projection (UMAP) plot showing all cells from a scRNA-seq experiment involving vascular beds from Day 1, Day 4 using low flow, and Day 4 using high flow (left). Additional plots show identified cell types (i.e. T1-T10) for each experimental condition (3 on the right). “BMVECs” (brain microvascular endothelial cells) refers to the endothelial cell line used. “NHLFs” and “HBVPs” (normal human lung fibroblasts and human brain vascular pericytes) refer to the stromal cell lines used. **(H)** Schematic of identified cell types. **(I)** Log2 fold change rank plots of differentially expressed genes between Day 4 low flow and Day 4 high flow. Highlighted genes have an adjusted p-value < 0.05 and log2 fold change magnitude > 0.25.

NHLFs formed 6 cell types (i.e. T3-T8) (**Fig. 4G, H; Extended Data Fig. 9D**). Although marker genes for T3-T8 were not substantially expressed in the NHLF cell line grown alone in monolayer when scRNA-seq pseudobulk was compared to bulk RNA-seq data, it is unclear to what degree these cell fates were acquired on VIVOS (**Extended Data Fig. 9E**). Mapping of two human lung scRNA-seq atlases revealed that T3 resembled myofibroblasts, T4 resembled smooth muscle cells, and T7 resembled pericytes (**Extended Data Fig. 9F-I**) ^29,30^. Mapping results for T5 were mixed. T6 markers were enriched for apoptotic and cell stress genes (**Supplementary Data 1**), while T8 was marked by proliferation (**Supplementary Data 1**). HBVPs formed 2 cell types (T9 & T10) (**Fig. 4G, H; Extended Data Fig. 9J**), with T10 being proliferative (**Supplementary Data 1**). Unexpectedly, mapping of a human brain scRNA-seq atlas revealed that T9 resembled brain fibroblasts (**Extended Data Fig. 9K, L**) ^31^.

BMVECs formed two cell types (T1 & T2) (**Fig. 4G, H; Extended Data Fig. 10A**). T1 was categorized as angiogenic endothelial cells which likely made up the vascular bed due to increased expression of VEGFA up-regulated genes (**Extended Data Fig. 10B, C; Supplementary Data 2**) ^32^, decreased expression of VEGFA down-regulated genes, increased expression of angiogenesis related genes (e.g. *INSR*, *ANGPT2*, *CXCR4*) (**Extended Data Fig. 10C**), and expression similarity with tip cell and stalk-like (immature) clusters from a human tumor angiogenesis atlas (**Extended Data Fig. 10D**) ^33^. In contrast, T2 was categorized as non-angiogenic endothelial cells which were likely lining the microfluidic channels because of the absence of the above features and their expression similarity with BMVECs grown in monolayer (**Extended Data Fig. 10E**). Concordantly, immunostaining for collagen type IV, encoded by top T1 marker genes *COL4A1* and *COL4A2* (**Supplementary Data 1**), was also markedly stronger in endothelial cells from the vascular bed compared to those from the microchannel (**Extended Data Fig. 10F**). RNA velocity analysis on Day 1 predicted a transition from T2 to T1, indicating that induction of T1’s angiogenic identity occurred early during the vessel formation process (**Extended Data Fig. 10G**).

We first observed large transcriptional changes for all cell types during the vessel formation process, irrespective of flow rates, between Day 1 and Day 4, with stromal cell types (i.e. T3-T10) sharing many differentially expressed genes (DEGs) (**Fig. 4G; Extended Data Fig. 11A, B; Supplementary Data 3**). Interestingly, we observed higher expression of immune related genes such as *ISG15*, *IFIT3*, and *IFI6* on Day 1 when compared to Day 4 across all cell types except T4, as well as enrichment for immune related gene sets on Day 1 (**Extended Data Fig. 11B, C**). T1 endothelial cells on Day 1 further expressed higher levels of the endothelial inflammation genes *ICAM1* and *CCL2* ^34^. Notably, analysis of vascular beds using a bulk RNA-seq timecourse revealed that these immune related genes were induced on Day 1 when compared to the starting Day 0 population (**Extended Data Fig. 11D**). Consistent with their roles in producing the extracellular matrix (ECM), stromal cells on Day 4 displayed increased expression of ECM components such as *COL1A1/2* and were enriched in ECM related gene sets (**Extended Data Fig. 11B, E**). T1 endothelial cells, consistent with formation of the vasculature, were enriched in angiogenesis and cell-substrate adhesion gene sets on Day 4 (**Extended Data Fig. 11B, E**). These ECM and angiogenesis signatures were all upregulated on Day 4 when compared to the starting Day 0 population (**Extended Data Fig. 11D**). Taken together, our observations indicate that an early immune response program on VIVOS is progressively replaced by ECM production and angiogenic signaling as vessels form. Given that immune responses, ECM production, and angiogenesis are all hallmarks of wound healing ^35^, we highlight the resemblance of vascular bed formation on VIVOS with this biological process and speculate that future experiments could involve using VIVOS to model wound healing responses and fibrosis. Interestingly on Day 4, stromal cell types, particularly T3, T4, and T9, expressed higher levels of chemokines such as *CXCL1*, *CXCL6* and *CXCL8* (**Extended Data Fig. 11B; Supplementary Data 3**). Notably, the corresponding receptors for these ligands, CXCR1 and CXCR2, are primarily found in immune cells such as neutrophils ^36^ and were not significantly detected in any cell type on VIVOS, suggesting that addition of immune cells to VIVOS could be an important next step for the system.

We next focused on exploring the effects of flow and shear stress. Although this is typically studied in the context of endothelial cells, we found that the most dramatic changes in gene expression between Day 4 low flow and high flow occurred in stromal cells (**Fig. 4G, I; Supplementary Data 4**). Notably, sub-clustering of T3 and T9, the largest stromal cell types by size, revealed that distinct sub-clusters were associated with Day 4 low flow and high flow (**Extended Data Fig. 12A, B**). Stromal cell types (i.e. T3-T10) shared many DEGs between Day 4 low flow and high flow (**Extended Data Fig. 12C**), indicating that their mechanisms for responding to flow were similar. Interestingly, we observed higher expression of chemokines such as *CXCL2*, *CXCL3*, *CXCL5* and *CXCL6* on low flow (**Fig. 4I; Supplementary Data 4**). Laminar shear stress is known to be anti-inflammatory in endothelial cells ^20^, and our data extends this observation to show that flow also reprograms inflammatory chemokine production within the stromal compartment.

We next examined DEGs between Day 4 low flow and high flow conditions in combination with cell-cell communication analysis and found that many of the DEGs in stromal cells encode ligands minimally expressed in endothelial cells but predicted to interact with endothelial cell localized receptors (**Extended Data Fig. 12D, E**). Notably, these DEGs included potential angiogenic regulators such as *WNT5B* for high flow and *CXCL12* for low flow (**Extended Data Fig. 12E**) ^37,38^. We conclude that the effect of flow on the vasculature not only directly impacts endothelial cells as is well known ^20^, but can also alter stromal cells by inducing alterations in ligand production. Although endothelial cells themselves appeared less affected by flow, we found within T1 endothelial cells that *APLN*, encoding the Apelin ligand, was the top DEG for low flow and *APLNR*, encoding the Apelin receptor, was the top DEG for high flow (**Fig. 4I**). *APLN* and *APLNR* were also strongly differentially expressed in our bulk RNA-seq comparisons of low flow and high flow (**Extended Data Fig. 13A**). We therefore focused further investigations on Apelin signaling.

### Regulation of Vessel Formation by a YAP/TAZ-Apelin Signaling Axis

Further examination of *APLN* and *APLNR* revealed that both genes were uniquely expressed in endothelial cells (**Extended Data Fig. 13B**), upregulated during vessel formation (**Extended Data Fig. 13C**), and among the most variably expressed genes (**Extended Data Fig. 13D**). Expression of the other Apelin receptor ligand, *APELA* ^39^, was not detected in any cell type. Most notably, the expressions of *APLN* and *APLNR* were negatively correlated on VIVOS as well as in the human tumor angiogenesis atlas from Goveia *et al*. (**Fig. 5A; Extended Data Fig. 13E, F**) ^33^. To uncover pathways that regulate *APLN*, we analyzed *APLN* gene co-expression patterns on VIVOS and in the human atlas. This revealed that *CTGF*, *CYR61*, *AMOTL2* and *ANKRD1*, canonical YAP/TAZ targets, were among the most strongly positively correlated transcripts (**Fig. 5B; Extended Data Fig. 13F, G**). Together with the anti-correlated expression of *APLN* and *APLNR*, these associations suggested that YAP/TAZ might partition endothelial cells into *APLN*-high/*APLNR*-low versus *APLN*-low/*APLNR*-high states. Notably, transcript levels of VGLL4, which binds TEADs and antagonizes YAP/TAZ-dependent transcription ^40^, were anti-correlated with *APLN* (**Fig. 5B**). To test whether YAP/TAZ regulates *APLN* and *APLNR*, we treated monolayer endothelial cells with IAG933, a YAP/TAZ-TEAD inhibitor ^41^, and observed potent downregulation of *APLN* and concomitant upregulation of *APLNR* (**Fig. 5C; Extended Data Fig. 13H**). Reanalysis of published YAP, TAZ, and TEAD1 ChIP-seq data from endothelial cells revealed binding sites for all three factors at regulatory regions near the *APLN* gene (**Extended Data Fig. 13I**) ^19^. Altogether, our observations are supported by previous identification of Tead1 as a regulator of the mouse *Apln* promoter ^42^, and further consistent with recent work showing that upregulation of mouse *Apln* in endothelial cells by ionizing radiation was YAP/TAZ dependent ^43^. We conclude that the negative correlation between *APLN* and *APLNR* expression is explained by opposing effects of YAP/TAZ, with evidence for a direct transcriptional effect on *APLN*. Regulation of *APLNR* by YAP/TAZ possibly occurs through intermediate regulation of KLF2/KLF4, which were previously identified to promote *APLNR* expression (**Extended Data Fig. 13H**) ^44^.

**Fig. 5.**
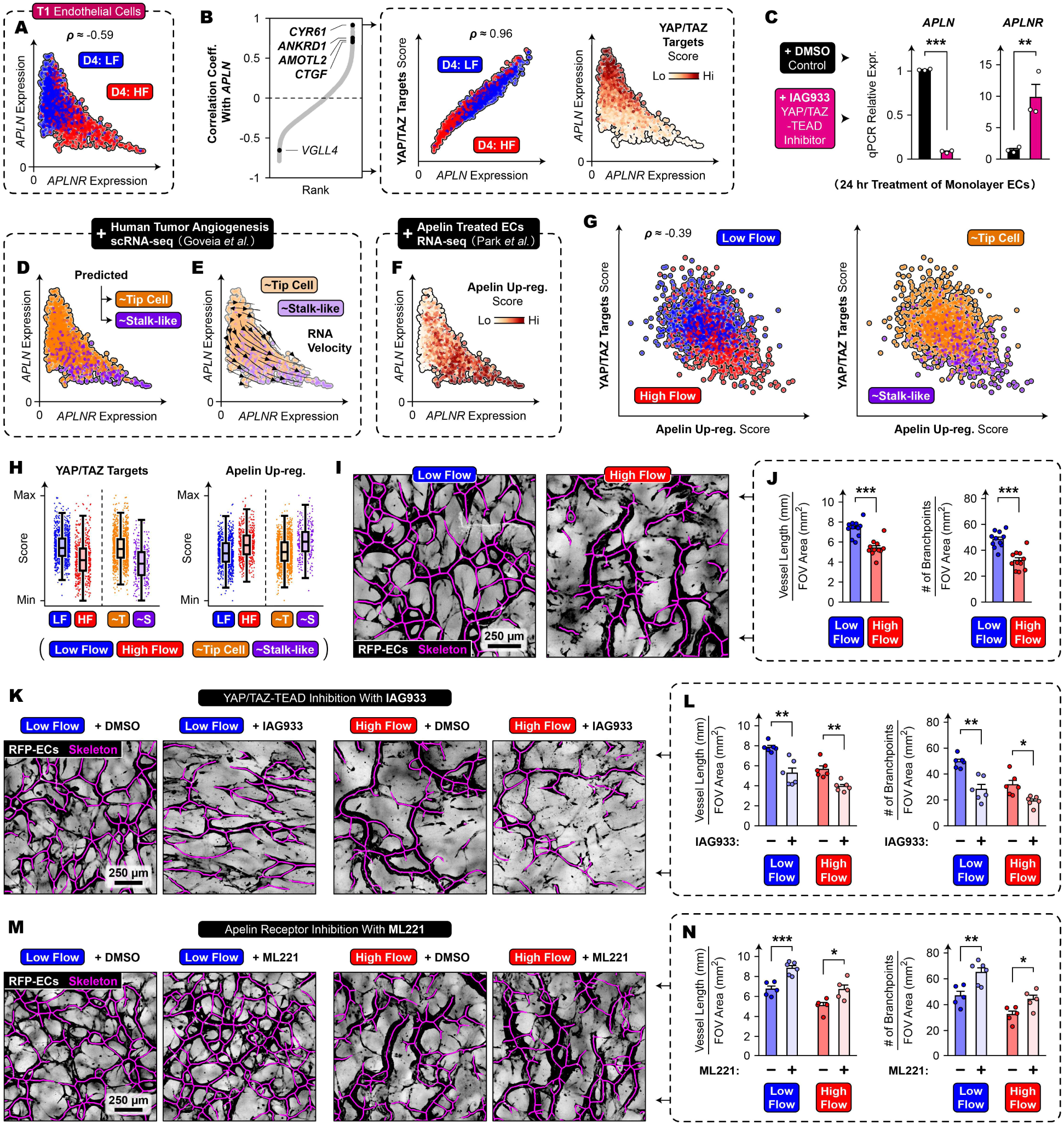
Vascular flow regulates vessel network formation through YAP/TAZ and Apelin signaling. **(A)** T1 endothelial cells on Day 4 plotted for *APLN* and *APLNR* expressions. **(B)** Rank plot of all analyzed genes and their correlation coefficient with *APLN* (left). Correlation of *APLN* expression with gene set scoring for YAP/TAZ targets (center). Overlay of scores onto the scatter plot of *APLN* and *APLNR* expressions (right). Expression values are from T1 endothelial cells on Day 4. **(C)** qPCR expression data of *APLN* and *APLNR* from monolayer endothelial cells treated with IAG933 (2 µM; 24 hrs; 3 independent experiments; 2-tailed unpaired student’s t-test). **(D, E)** Mapping tip cell and stalk-like clusters from the human tumor angiogenesis atlas of Goveia *et al*. ^33^ onto T1 endothelial cells on Day 4, with (E) additionally showing RNA velocity streamlines. **(F)** Gene set scoring of T1 endothelial cells on Day 4 for Apelin up-regulated genes (Park *et al*., ^57^). **(G)** Negative correlation between gene set scores for YAP/TAZ targets and Apelin up-regulated genes. **(H)** Box plot summaries of (G). **(A, B, G)** Spearman’s correlation coefficient. **(A, B, D-F)** Axes show log-normalized expression values of *APLN* and *APLNR* denoised using Markov Affinity-based Graph Imputation of Cells. **(I)** Skeleton visualizations of vascular beds grown using low or high flow on Day 4. **(J)** Quantification of (I) (11 devices per condition across 6 independent experiments). Data is a subset of (L) and (N). **(K)** Skeleton visualizations of low/high flow vascular beds on Day 4 treated with IAG933 (2 µM). **(L)** Quantification of (K) (6 devices per condition across 3 independent experiments). **(M)** Skeleton visualizations of low/high flow vascular beds on Day 4 treated with ML221 (10 µM). **(N)** Quantification of (M) (5-6 devices per condition across 3 independent experiments). **(J, L, N)** 2-tailed unpaired Welch’s t-test. **(C, J, L, N)** * p < 0.05; ** p < 0.01; *** p < 0.001. Error bars show SEM.

YAP/TAZ are strongly implicated in mechanotransduction ^45–47^, with their regulation by fluid shear stress depending on the type of flow. Unidirectional laminar shear stress has been shown to suppress YAP/TAZ, whereas oscillatory non-laminar shear stress is known to activate YAP/TAZ ^46,47^. The pattern of differential expression between low flow and high flow of *APLN* and *APLNR*, which we found to be regulated by YAP/TAZ, indicates that shear stress on VIVOS inhibits YAP/TAZ activity. This is consistent with reduced scRNA-seq scoring for YAP/TAZ targets on high flow (**Fig. 5A, B, G, H**), and reduced expression of predicted YAP/TAZ regulated genes on high flow when analyzing bulk RNA-seq data (**Extended Data Fig. 14A-C**). Given the dimensions and measured fluid velocities of vessels on VIVOS, we estimate a Reynolds number of around 0.01-0.1, which indicates laminar flow, and provide agreement with previous studies that laminar shear stress has a suppressive effect on YAP/TAZ. As an independent validation of our VIVOS observations, we applied shear stress to monolayer endothelial cells using an orbital shaker (**Extended Data Fig. 14D**). Because this method produces different flow regimes within the culture dish ^48^, we only cultured cells around the dish’s periphery and avoided the non-laminar regions at the dish’s center. In the following experiments, we focused on TAZ because its transcript levels were more abundant than those of YAP in endothelial cells (**Extended Data Fig. 14E**) ^19^. Mirroring high flow on VIVOS, shear stress using the orbital shaker method downregulated *APLN* and YAP/TAZ targets while upregulating *APLNR* (**Extended Data Fig. 14F**). Once YAP/TAZ are phosphorylated, they are targeted for degradation ^49,50^. Concordantly, TAZ immunostaining within both the nucleus and cytoplasm were decreased by shear stress without changing TAZ transcript levels (**Extended Data Fig. 14F-H**). YAP immunostaining was similar to that of TAZ (**Extended Data Fig. 14I**). This decrease in the levels of YAP/TAZ, specifically in the nucleus, is consistent with the reduced expression of YAP/TAZ target genes we observed (**Extended Data Fig. 14F**). Endothelial cells under shear stress additionally displayed lower immunostaining for Apelin protein (**Extended Data Fig. 14J, K**). Apelin signal was predominantly localized to the Golgi apparatus (**Extended Data Fig. 14L**), consistent with its detection during the secretory process. Cryosections of low flow and high flow vascular beds reflected these monolayer shear stress experiments, with lower overall TAZ immunostaining per vessel. (**Extended Data Fig. 14M, N**).

Overall, we find that YAP/TAZ acts as a shear stress sensitive “switch” that partitions endothelial cells into *APLN*-high/*APLNR*-low and *APLN*-low/*APLNR*-high populations. Under physiological shear stress, such as with high flow on VIVOS, YAP/TAZ is suppressed which then biases endothelial cells towards the *APLN*-low/*APLNR*-high state.

During angiogenesis, tip cells sprout to guide new vessels while stalk cells form the nascent lumen^16^. Although vascular bed formation on VIVOS more closely resembles vasculogenesis (**Extended Data Fig. 3A**), where tip and stalk cells are less clearly defined, a tip-stalk framework was informative. In mouse retinal vasculature, *Apln* is expressed in tip cells and *Aplnr* is expressed in stalk cells ^17,44^. In zebrafish, *apln* is expressed in tip cells while *aplnrb* expression is detected in both tip and stalk cells ^51^. Thus, deployment of Apelin and its receptor across tip and stalk compartments appears to be either species specific or context dependent. To examine Apelin signaling and tip-stalk from a human context, we leveraged the tip cell and stalk-like (immature) endothelial clusters described by Goveia and colleagues to predict tip and stalk-like states on VIVOS (**Fig. 5D; Extended Data Fig. 15A-D**) ^33^. Predicted tip cells showed higher expression of *ANGPT2*, *PGF*, and *ADM*, whereas predicted stalk-like cells expressed higher *TEK*, *HES1*, and *ID1*, consistent with established tip-stalk signatures (**Extended Data Fig. 15B, C**) ^17,52–54^. Consistent with the mouse studies, low flow (enriched for *APLN*-high/*APLNR*-low cells) contained more predicted tip cells, whereas high flow (enriched for *APLN*-low/*APLNR*-high cells) contained more predicted stalk-like cells (**Fig. 5D; Extended Data Fig. 15D**). RNA velocity supported a transition from predicted tip to predicted stalk-like states, consistent with resolution of sprouting as vessels mature (**Fig. 5E**). Because Goveia *et al*.’s atlas was tumor associated and they described their “immature” cluster as resembling stalk cells, we additionally performed scRNA-seq scoring using curated lists of tip and stalk cell genes (**Extended Data Fig. 15E; Extended Data Table 1**). Consistent with the Goveia atlas-based predictions, curated tip scores were higher under low flow and in *APLN*-high/*APLNR*-low cells, whereas curated stalk scores were higher under high flow and in *APLN*-low/*APLNR*-high cells (**Extended Data Fig. 15F**). Together, these analyses support a flow-conditioned *APLN*-*APLNR* state axis that aligns with tip-stalk endothelial remodeling programs.

At the tissue level, we next used vessel length density and branchpoint density as quantitative readouts of tip cell activity, consistent with the seminal Dll4-Notch1 study ^55^. These metrics can be robustly extracted by image analysis and directly correspond to sprouting. Consistent with fewer predicted tip cells, high flow reduced both vessel length and branchpoint densities, indicative of fewer tip-like phenotypes (**Fig. 5I, J**). To explore YAP/TAZ using these metrics, we treated vascular beds on VIVOS with the YAP/TAZ-TEAD inhibitor IAG933 and found reduced vessel length and branchpoint densities, indicating a loss of tip-like sprouting (**Fig. 5K, L**). We further observed that predicted tip cells showed higher scoring for YAP/TAZ target genes when compared to predicted stalk-like cells, with similar results for the human tumor angiogenesis atlas (**Fig. 5G, H; Extended Data Fig. 15G**) ^33^. Overall, these results support a role for YAP/TAZ in tip cell sprouting, aligning with previous studies ^19,56^.

We next explored the role of Apelin signaling itself by deriving an Apelin up-regulated gene signature from Park *et al*.’s bulk RNA-seq of endothelial cells under flow treated with Apelin (**Fig. 5F**) ^57^. Stalk-like cells scored higher for Apelin up-regulated genes in both VIVOS’s and Goveia *et al*.’s datasets (**Fig. 5G, H; Extended Data Fig. 15G, H**) with high flow cells also scoring higher. Apelin receptor inhibition with ML221 increased vessel length and branchpoint densities on VIVOS, indicative of more tip-like phenotypes (**Fig. 5M, N**). Together, the scRNA-seq scoring and inhibitor experiments for both YAP/TAZ and Apelin indicate that they have opposing effects: YAP/TAZ promotes a tip-like, sprouting state, whereas Apelin signaling favors a stalk-like, non-sprouting state (**Fig. 5G, H; Extended Data Fig. 15I**). Differential sensitivity to Apelin appears to be imposed by YAP/TAZ: tip-like, YAP/TAZ-high cells are *APLN*-high/*APLNR*-low and thus relatively Apelin insensitive whereas stalk-like, YAP/TAZ-low cells are *APLN*-low/*APLNR*-high and Apelin responsive. This is consistent with the positive association between *APLNR* levels and Apelin-upregulated scores (**Fig. 5F; Extended Data Fig. 15H**). Despite the previously described dynamic interchangeability of tip and stalk cells ^58^, this differential sensitivity provides a mechanism to stabilize functional differences, as Apelin ligand produced by tip-like cells preferentially act on stalk-like cells which express the Apelin receptor. The downstream pathways by which Apelin signaling promotes stalk phenotypes remain unclear. Although GPCR regulation of YAP/TAZ has been described ^59^, and the Apelin receptor is a GPCR ^60^, we did not find a direct effect of Apelin on TAZ immunostaining even when *APLNR* expression was induced beforehand by applying shear stress (**Extended Data Fig. 14F; Extended Data Fig. 15J, K**). Overall, our findings support a model in which physiological laminar intraluminal shear stress reduces YAP/TAZ-TEAD output and biases an endothelial state transition along the Apelin axis, from *APLN*-high/*APLNR*-low, tip cell states toward *APLN*-low/*APLNR*-high, stalk-like states, consistent with a flow-controlled YAP/TAZ-Apelin “switch” that couples hemodynamic forces to molecular control of vessel remodelling (**Extended Data Fig. 15L, M**).

### Modeling Hereditary Hemorrhagic Telangiectasia with Intraluminal Vascular Flow

We next explored vascular flow in a disease context. Hereditary Hemorrhagic Telangiectasia (HHT) is a condition characterized by arteriovenous malformations (AVMs) which appear clinically as clusters of enlarged vessels with increased flow ^23^. This autosomal dominant disorder is caused by mutations in TGFβ/BMP pathway components, predominantly *ENG* (encoding Endoglin) and *ALK1* (also known as *ACVRL1*). Less frequently, mutations are found in their ligand *BMP9* (also known as *GDF2*) or an intracellular mediator, *SMAD4* ^23^. Genetic *in vivo* studies have been instrumental in linking these mutations to AVM formation and in proposing mechanisms involving dysregulated angiogenesis, inflammation, or flow sensing ^23^. However, existing models involve a trade-off: genetic *in vivo* studies rely on non-human vasculature, whereas prior microfluidic HHT platforms lack physiological intraluminal flow in perfused human vasculature with directly quantifiable flow velocities ^128, 129^. To address these limitations, we modeled HHT on VIVOS by generating vascular beds using endothelial cells expressing shRNAs targeting *ENG* or *ALK1* (**Fig. 6A; Extended Data Fig. 16A**), creating a fully human, flow-quantified AVM-like context in which vessel calibres and intraluminal flow velocities can be directly measured. In both cases, vessels generated from *ENG*- and *ALK1*-knockdown endothelial cells were enlarged with substantially higher vascular flow velocities as determined by tracking of fluorescent beads (**Fig. 6A-D, Supplementary Video 6**), consistent with observations in HHT patients. In contrast, treatment with BMP9 yielded vessels which were thinner in diameter and non-perfusable (**Fig. 6A-D**). While abundant evidence has shown that loss of Endoglin, ALK1, SMAD4 or BMP9 leads to the development of AVMs, the underlying cellular mechanisms are still not fully understood^23,61^. Hyperproliferation of endothelial cells has been proposed to drive AVMs in HHT ^62,63^.

**Fig. 6.**
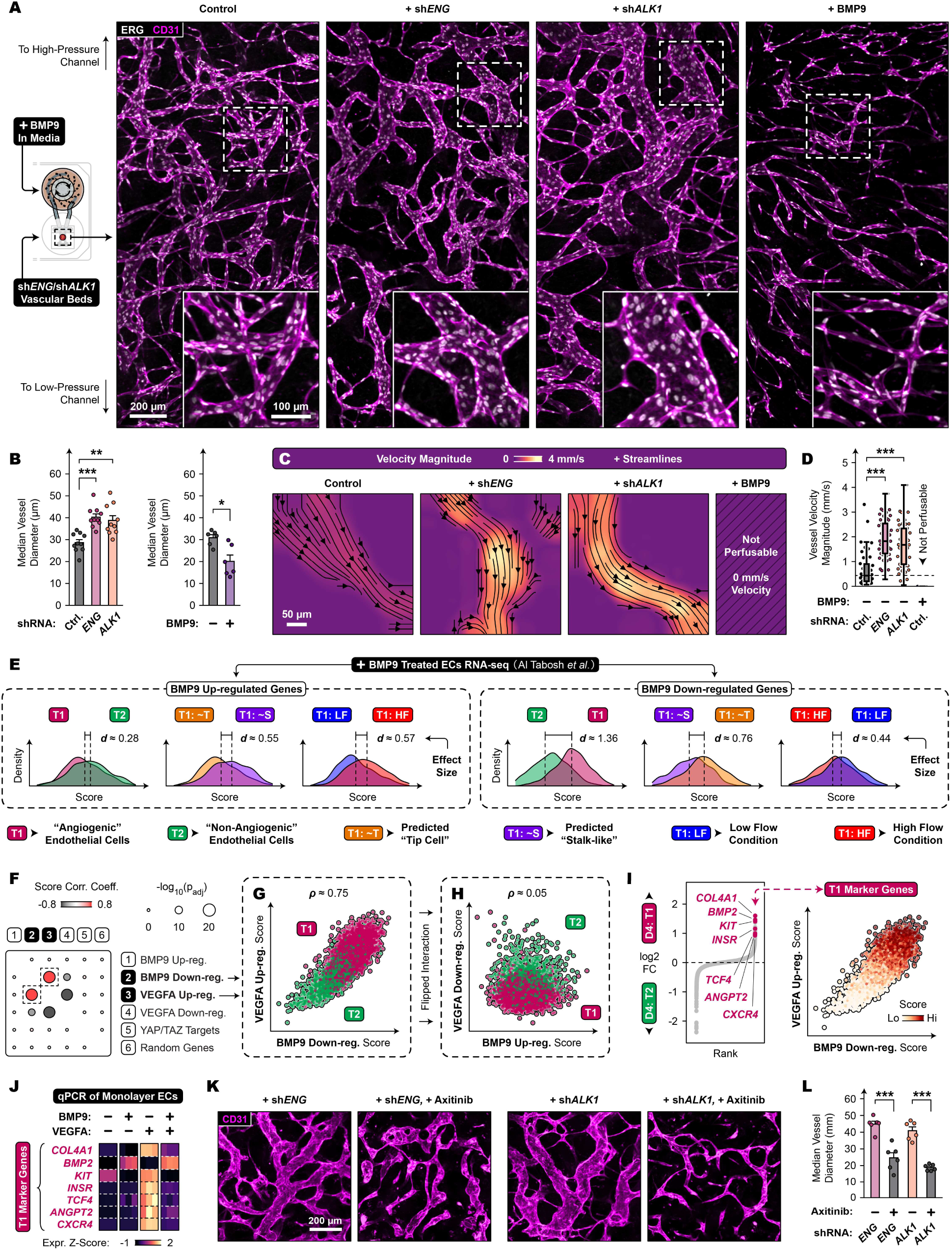
VIVOS models hereditary hemorrhagic telangiectasia with intraluminal vascular flow. **(A)** Vascular beds on Day 4 which were grown from *ENG*- or *ALK1*-knockdown endothelial cells or treated with 10 ng/mL BMP9 (ERG: endothelial nuclei, CD31: vessels). **(B)** Quantification of (A) (10 devices per condition across 4 independent experiments for shRNA vessels; 6 devices per condition across 2 independent experiments for BMP9 treatments). **(C)** Particle image velocimetry analysis showing velocity magnitude for *ENG*-knockdown, *ALK1*-knockdown, and BMP9-treated vessels on Day 4. **(D)** Quantification of (C) (42-45 vessels and 6 devices per condition across 2 independent experiments). **(E)** Gene set scoring of endothelial cells on Day 4 from the scRNA-seq experiment. Scores are for BMP9 up/down-regulated genes from Al Tabosh *et al*. ^64^. Endothelial cells are divided by either cell type (i.e. T1 or T2), predicted labels from Goveia *et al*. (i.e. tip cell or stalk-like), or experimental condition (i.e. low or high flow). Cohen’s d effect sizes are shown. **(F)** Correlation coefficients between gene set scoring of BMP9 up/down-regulated genes (Al Tabosh *et al*.), VEGFA up/down-regulated genes (Zhang *et al*., ^32^), and YAP/TAZ targets. Adjusted p-values were calculated using a permutation test followed by Bonferroni correction. **(G)** Gene set scoring of BMP9 down-regulated genes and VEGFA up-regulated genes. **(H)** Gene set scoring of BMP9 up-regulated genes and VEGFA down-regulated genes. **(I)** Log2 fold change rank plot showing a subset of markers genes for the T1 cell type (left). Gene set scoring for the subset of marker genes (right). **(F-H)** All endothelial cells from the scRNA-seq dataset were used. **(J)** Heatmap of qPCR gene expression from monolayer BMVECs treated with BMP9 (10 ng/mL) and/or VEGFA (100 ng/mL) for 48 hrs (3 independent experiments). **(K)** Images of *ENG*- or *ALK1*-knockdown vascular beds on Day 4 treated with 100 nM Axitinib (CD31: vessels). **(L)** Quantification of (K) (6 devices per condition across 2 independent experiments). **(B, D, L)** 2-tailed unpaired Welch’s t-test (* p < 0.05; ** p < 0.01; *** p < 0.001). **(B, L)** Error bars show SEM.

Analysis of the overall number of endothelial cells in *ENG*- and *ALK1*-knockdown or BMP9-treated vascular beds was comparable to controls with few proliferating (MKI67+) cells detected in any of the vessels (**Extended Data Fig. 16B, C**), suggesting that excessive endothelial cell proliferation is not a key driver of malformed vessels on VIVOS.

To gain insights into how BMP9/ALK1/Endoglin might regulate vessel morphologies, we scored endothelial cells from our scRNA-seq experiment for BMP9 up/down-regulated genes previously identified in a bulk RNA-seq study by Al Tabosh *et al*. (**Fig. 6E**) ^64^. On Day 4, BMP9 up-regulated genes scored higher in endothelial cells from the non-angiogenic cell type (T2), predicted stalk-like cells, and endothelial cells from the high flow condition. Conversely, BMP9 down-regulated genes scored higher in endothelial cells from the angiogenic cell type (T1), predicted tip cells, and endothelial cells from the low flow condition. These results were additionally confirmed by Gene Set Enrichment Analysis (**Extended Data Fig. 17A**). Altogether, these results suggest that BMP9/ALK1/Endoglin signaling is anti-angiogenic, stalk promoting, and is further enhanced by fluid shear stress. Notably, the most pronounced differences were observed between the T1 angiogenic and T2 non-angiogenic cell types for BMP9 down-regulated genes (**Fig. 6E**).

We next assessed potential interactions of BMP9 signaling with that of VEGF and YAP/TAZ, two pathways involved in angiogenesis ^16,19,65^. Analysis of correlation coefficients associated with BMP9 ^64^, VEGFA ^32^, and YAP/TAZ regulated genes revealed a striking positive correlation between BMP9 down-regulated genes and VEGFA up-regulated genes (**Fig. 6F, G**), which was not due to a major overlap of gene sets (**Extended Data Fig. 17B**). This positive correlation was also evident in the human tumor angiogenesis atlas (**Extended Data Fig. 17C**) ^33^. Of note, the flipped interaction of BMP9 up-regulated and VEGFA down-regulated genes displayed minimal correlation (**Fig. 6H**). Despite previous work suggesting a link between YAP/TAZ and BMP9/ALK1/Endoglin signaling ^66^, related gene expression correlations were negligible (**Fig. 6F; Extended Data Fig. 17D**).

The inverse correlation between BMP9 down-regulated genes and VEGFA up-regulated genes suggests that BMP9 antagonizes VEGFA-induced transcription. To test this, we treated monolayer endothelial cells with VEGFA and examined the expression of a subset of top marker genes from the T1 angiogenic cell type (*COL4A1*, *BMP2*, *KIT*, *INSR*, *TCF4*, *ANGPT2*, and *CXCR4*) (**Fig. 6I**) that were progressively up-regulated on Day 1 during early vessel formation (**Extended Data Fig. 17E**) and also mostly up-regulated in the tip cell and stalk-like human atlas clusters (**Extended Data Fig. 17F**). Analysis by qPCR revealed up-regulation of these marker genes (**Fig. 6J; Extended Data Fig. 17G, H**), consistent with a role for VEGF signaling in initiating the T1 angiogenic cell type. Co-treatment with BMP9 abolished the VEGFA-induced increase for 6 out of 7 of these genes, supporting an antagonistic role for BMP9 on VEGFA-induced transcription. The one exception, *BMP2*, was not identified as an angiogenic marker in the human atlas ^33^ and thus may be a VIVOS specific anomaly (**Extended Data Fig. 17F**). Compared to VEGFA treatment alone, treatment with VEGFA and BMP9 together had minimal effects on expression of *VEGFR1* and *VEGFR2*, suggesting that mechanisms other than transcriptional control of VEGF receptors contributed to VEGF signaling antagonization by BMP9 (**Extended Data Fig. 17I**). Of note, the expression of the decoy receptor *VEGFR1* was positively correlated with scoring of VEGFA up-regulated genes (**Extended Data Fig. 17J**). Interestingly, BMP9 treatment alone only decreased a few of the angiogenic marker genes (*KIT*, *CXCR4*) compared to BMP9 treatment in the presence of VEGFA, with *COL4A1*, *INSR*, *TCF4*, and *ANGPT2* showing little to no change (**Fig. 6J; Extended Data Fig. 17G**).

Taken together, these analyses indicate that BMP9 suppresses a VEGF-induced angiogenic gene expression program in endothelial cells, and that this suppression appears to be dysregulated in HHT (**Extended Data Fig. 17K**). These results are in line with anti-angiogenic therapies which involve targeting VEGF signaling, currently under investigation for the treatment of HHT ^23,61,67^. In support of this, we found that *ENG*- and *ALK*-knockdown vessels grown in the presence of the VEGF receptor inhibitor Axitinib produced vessels with thinner diameters (**Fig. 6K, L**). Overall, VIVOS supports the use of physiological intraluminal flow in fully human vasculature to model HHT and generate flow-quantified AVM-like lesions, and further opens avenues for exploring flow-regulated pathology and preclinical intervention strategies in other vascular diseases.

## Discussion

Until now, no fully human system has allowed intraluminal flow, endothelial state transitions, and vessel remodeling to be interrogated together under defined hemodynamic conditions in large complex 3D tissues. VIVOS addresses this need, enabling mechanistic dissection of how flow reshapes human vascular remodeling at the level of endothelial cell states. In addition to enabling biological discoveries, VIVOS also has potentially important applications in regenerative medicine, particularly for tissue replacement therapies ^68^. Current research avenues focus on transplanting either single cells or small multicellular aggregates (e.g. spheroids) ^69^. Although transplanting larger, intact tissues could better preserve their organization and function, scaling *in vitro* cultures towards organ-sized constructs requires continuous vascular flow to overcome diffusion limits. We envision that enlarged future iterations of VIVOS, which uses an electromechanical pump to provide continuous vascular flow, could enable growth of larger human tissues *in vitro*. A more immediate challenge, particularly with transplantation of pancreatic islets and liver tissue, is graft necrosis due to inadequate vascularization at the transplantation site ^70^. Through its ability to vascularize diverse tissue shapes and sizes, VIVOS could support strategies for pre-vascularizing grafts prior to transplantation.

Vascular development is tightly regulated with deleterious consequences for both too many and too few vessels. The classical explanation emphasizes feedback driven by local nutrient and hypoxia sensing to either promote or repress angiogenesis ^71^. Fluid shear stress may provide an additional layer of negative feedback regulation by coupling perfusion with sprouting behavior, as proposed previously ^72^. Specifically, lower shear in poorly perfused vascular beds would favor tip-like sprouting to grow new vessels, whereas higher shear in well-perfused beds would promote stalk-like stabilization and limit excess angiogenesis. Consistent with this model, we observe on VIVOS that increasing flow has a pruning-like effect by producing sparser networks, which mechanistically involves a shear-responsive YAP/TAZ-TEAD “switch” that regulates Apelin pathway components and shifts endothelial cell states towards stalk-like programs. Notably, spatially restricted activation of Apelin signaling, in line with spatially distinct tip/stalk states, is supported by the Apelin ligand’s minute-scale *in vivo* half-life ^73^, and the observation of self-generated gradients of Toddler/Elabela, the other Apelin receptor ligand, during zebrafish gastrulation ^74^. Because physiological flow is difficult to tune *in vivo*, VIVOS provides a useful platform for causally dissecting how hemodynamics pattern endothelial cell states and vascular architecture in a fully human 3D context. Future work using VIVOS could additionally investigate within an all-human *in vitro* setting how upstream shear sensors, such as integrin-RhoA signaling ^47^, couple flow to YAP/TAZ activity.

Although antagonization of VEGF signaling by BMP9 is known ^67^, the mechanisms and downstream effects are not fully understood. While previous work explored intracellular mediators such as PI3K/Akt and ERK ^67,75,76^, we describe inhibition of VEGF by BMP9 through repression of VEGFA-induced genes, reflecting results observed from inhibiting ALK1 signaling using an ALK1 ligand trap ^77^. Interestingly, we did not find substantial evidence for the proposed model of BMP9-induced *VEGFR1* expression as a mechanism for VEGF signaling inhibition ^53,67^. It thus remains unclear where in the VEGF signaling pathway BMP9 intervenes in order to repress VEGFA-induced genes. In HHT, AVMs have faster flowing vessels, while other diseases such as Cerebral Cavernous Malformations (CCM) produce slower flowing vessels. Future work on VIVOS recreating different types of vascular malformations could also provide mechanistic insights into the opposing effects on vascular flow from different genetic mutations.

In addition to enlarging the device, future iterations of VIVOS could interconnect multiple units containing different tissue types to model multi-organ phenomena, and incorporate circulating components (e.g. immune cells or circulating tumor cells) by perfusing suspended cells through the fluidic circuit. The modular design of VIVOS makes interconnection straightforward, and constant agitation within the impeller pump and fast media recirculation would limit sedimentation of suspended cells (**Extended Data Fig. 5G**). Moreover, preliminary computational fluid dynamics modeling shows that shear stresses within the impeller pump casing, which may damage suspended cells if too high, are not considerably greater than what is already present within the vessels (**Extended Data Fig. 5F**). Together, these advances could enable fully human *in vitro* models of processes such as immune cell trafficking and cancer metastasis.

Taken together, VIVOS provides an experimentally tractable and physiologically relevant platform for recreating vascular flow *in vitro* by coupling a tunable electromechanical impeller pump to living human vasculature. This allows the vascularization of a wide range of 3D tissues, the interrogation of how physiological intraluminal flow drives a YAP/TAZ-Apelin switch, endothelial state transitions, and vessel remodelling, as well as the modeling of AVMs from Hereditary Hemorrhagic Telangiectasia. Considering ongoing efforts to reduce animal use in preclinical testing, VIVOS establishes a fully human flow-quantified preclinical model for studying vascular development, dissecting flow-regulated disease mechanisms, and evaluating candidate therapies in physiologically perfused 3D human tissues.

## Methods

### Microfluidic Chip Fabrication

Silicon wafer molds were fabricated at University of Toronto’s Center For Research And Applications In Fluidic Technologies (CRAFT). Designs drawn in AutoCAD (Autodesk) were transferred onto chrome photomasks using a uPG501 Mask Writer (Heidelberg Instruments). Depending on film thickness, either Su-8-50 or Su-8-100 photoresist (Kakayu) was deposited onto 100 mm diameter silicon wafers (Wafer World) using spin coating (Laurell Technologies). UV exposure using a mask aligner (OAI) transferred the designs from the photomask onto the photoresist. Removal of uncured photoresist produced finished molds.

Microfluidic channels were fabricated from polydimethylsiloxane (PDMS, Sylgard 184, Dow). Base and curing agents were mixed 10:1 by weight, poured onto wafer molds, degassed under vacuum, and cured for 12 hrs at 50°C. Vascular bed compartments were similarly fabricated from PDMS, with additional features created using 3 mm and 5 mm diameter biopsy punches.

To fabricate porous membranes, a wafer mold which features posts corresponding to pores was first silanized with trichloro(1H,1H,2H,2H-perfluorooctyl)silane (Sigma-Aldrich) through overnight deposition in a vacuum chamber followed by a 10 min incubation at 150°C on a hotplate. A 1 mm thick “sacrificial” sheet of cured PDMS was plasma treated (Harrick Plasma) for 5 min, followed by the deposition of 2.5% polyvinyl alcohol (Sigma-Aldrich) through spin coating (Specialty Coating Systems) at 800 RPM for 1 min (based on Karlsson *et al*.) ^78^. The coated sacrificial sheet was left for 12 hrs at 50°C, followed by a second plasma treatment for 5 min. Mixed but unpolymerized PDMS (10:1 base and curing agent ratio) was poured onto the wafer mold followed by degassing under vacuum. The sacrificial sheet was pressed against the unpolymerized PDMS, coated side facing down, with the aid of 30 lb weights. The assembly was left at room temperature for 1 hr, and then incubated for 18 hrs at 50°C. The sacrificial sheet along with the cured porous membrane was then peeled off the wafer and excised into appropriately sized pieces with a scalpel.

To assemble the microfluidic chips, individual components were plasma treated (Harrick Plasma) for 2 min, pressed together, and incubated for 12 hrs at 70°C. To remove the sacrificial slab from the porous membranes, bonded pieces were incubated in ultrapure water for 12 hrs at 70°C to dissolve the polyvinyl alcohol interface layer.

### Impeller Pump Fabrication

Impeller pump designs were drawn using Fusion360 (Autodesk), exported to PrusaSlicer (Prusa Research), and 3D printed using polylactic acid filament (MatterHackers) on a Prusa MK3S 3D printer (Prusa Research). Printed parts were sterilized by immersion in 70% ethanol for 12 hrs. Individual components were assembled using epoxy glue where required. Glue joints were designed to not be in contact with nutrient media. Impellers were further attached to 3.175 mm by 3.175 mm magnets (K&J Magnetics) and polytetrafluoroethylene (PTFE) tubing (Cole-Parmer), which acted as axles. These PTFE tubing axles rode against 12 mm diameter #1.5 glass coverslips (Electron Microscopy Sciences), which acted as flat bearings.

Final device assembly involved joining microfluidic chips to impeller pumps using 1.5875 mm inner diameter polyvinyl chloride tubing (Sigma-Aldrich). 3 chips and 3 pumps were arranged inside 6 well plates (Corning).

Each motor box, which was also 3D printed from polylactic acid filament (MatterHackers), contained 3 brushless DC motors with integrated proportional integral controllers (Faulhaber). Each motor was connected to 6.35 mm by 3.175 mm by 3.175 mm magnets (K&J Magnetics). 16 motor boxes per incubator were connected using I^2^C to a Raspberry PI computer using a PCA9685 16-channel pulse-width modulation (PWM) multiplexer (Adafruit). A custom user interface, displayed on a touch screen, was written in Python and Tkinter.

### Impeller Pump Experimental Flow Characterization

Flowrates were measured with SLI-0430 and SLI-2000 sensors (Sensirion). Pressure differentials were measured by connecting two vertical tubing segments, each open on one end, to either side of the impeller pumps. Values were obtained using *ΔP = pgΔh* by measuring the difference in water column heights between the two vertical tubing segments. To apply different flow resistances to the impeller pump, tubing was pinched off with progressively narrower gaps using a 3D-printed “comb”.

### Generation of Complex Waveforms Using the Impeller Pump

The impeller pump, through the Raspberry PI computer, I^2^C interface, and PWM input signal, can be programmed to have any waveform.

The following formula was utilized to generate the heartbeat-like flowrate waveform (related to **Extended Data Fig. 5I**, middle panel):

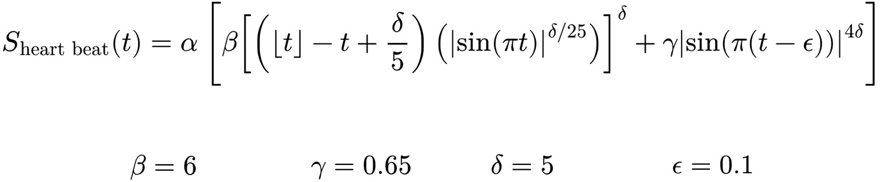

The following formula was utilized to generate the heart-shaped flowrate waveform (related to **Extended Data Fig. 5I**, bottom panel):

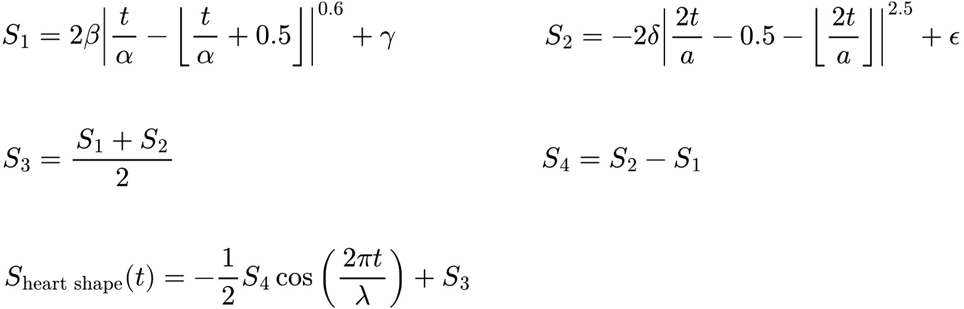

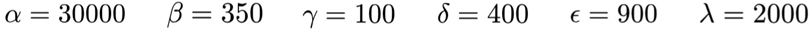

Because flowrate is proportional to the square of impeller speed (ω), I^2^C and PWM input signals were calculated by taking the square root of the waveforms (*S*).

### Computational Fluid Dynamics

Simulations were performed in COMSOL Multiphysics 6.2. For microfluidic chip simulations, we utilized laminar Navier Stokes. For impeller pump simulations, we utilized Navier Stokes with the k-ω turbulence model and frozen rotor analysis. All simulations utilized fluid properties of water. For analyzing the distribution of fluid shear stresses within the impeller pump casing (related to **Extended Data Fig. 5F**), values were weighted based on individual mesh element volumes.

### Cell Culture

Brain microvascular endothelial cells (BMVECs; Cell Systems; #ACBRI376) and RFP-labeled brain microvascular endothelial cells (RFP-BMVECs; Alphabioregen; #RFP2) were cultured in EGM-2 (Lonza). Unless otherwise stated, the non-labeled BMVECs were used. Normal human lung fibroblasts (NHLFs; Lonza; #CC-2512) were cultured in FGM-2 (Lonza). Human brain vascular pericytes (HBVPs; ScienCell; #1200) were cultured in Pericyte Medium (ScienCell). Cell lines were cryopreserved at passage 2 or 3 and used for 5 additional passages post-thaw.

### 3D Tissue Culture

Spheroids were generated from aggregation in 96-well V-bottom plates (Greiner Bio-One) using MCF10A cells (a gift from Dr. Daniel Schramek) which were further endogenously tagged with moxGFP at the AAVS1 locus using the qTAG system ^79^. 8-day-old spheroids were implanted on VIVOS. Human fetal eyes at fetal week 14 were collected through the services of the RCWIH Biobank under REB#1138. Retinal explant cultures were prepared as described previously ^80^. Vascular organoids were generated from GFP-tagged H9 hESCs as described previously with modifications ^7^. H9 hESCs (WiCell; #WA09) were tagged by inserting the GFP gene into the AAVS1 locus using transcription activator-like effector nucleases (TALENs). H9 aggregates were created in 96-well V-bottom plates (Greiner Bio-One) with 50 µM Y-27632 (Selleck Chemicals), followed by mesoderm induction and vascular differentiation according to the previous protocol. 2-week-old vascular organoids were implanted on VIVOS. Cerebral organoids were generated from H9 hESCs (WiCell; #WA09) according to Sivitilli *et al*. ^81^. 1-month-old cerebral organoids were implanted into VIVOS. Lung organoids were generated from WTC-11 iPSCs (Coriell Institute for Medical Research; #GM25256) as described previously with modifications ^82^. To induce endoderm, embryoid bodies were treated for 24 hrs with 3 µM CHIR (Sigma-Aldrich) and 100 ng/mL Activin A (ThermoFisher) in RPMI (Gibco), followed by 48 hrs with 100 ng/mL Activin A and 0.2% FBS, and 24 hrs with 100 ng/mL Activin A and 2% FBS. 1.5-month-old organoids were implanted on VIVOS.

### Vascular Bed Seeding

Prior to seeding, vascular bed compartments of microfluidic chips from assembled VIVOS devices were coated for 2 hrs with 2 mg/mL polydopamine (Sigma-Aldrich) freshly dissolved in 10 mM Tris-HCl pH 8.5, washed 3 times with ultrapure water, and left to air dry. This step reduced the risk of hydrogel detaching from the microfluidic chip during culture ^83,84^. BMVECs, NHLFs, and HBVPs were trypsinized, pelleted, and resuspended in a 2X concentrated hydrogel mixture dissolved in PBS without calcium and magnesium. An equal volume of PBS, optionally containing 3D tissues, was combined and mixed with the resuspended cells. The final mixture contained 750k BMVECs/mL, 1000k NHLFs/mL, and 250k HBVPs/mL, 10 mg/mL bovine fibrinogen (Sigma-Aldrich), 0.15 TIU/mL bovine aprotinin (BioShop), and 0.2 mg/mL rat tail collagen I (Corning). After addition of bovine thrombin (Sigma-Aldrich) to a final concentration of 3 U/mL, the mixture was immediately mixed by pipetting and then transferred to the vascular bed compartments of microfluidic chips. A 10 mm by 10 mm by 1 mm sheet of PDMS was placed on top of each vascular bed compartment to create seals with the microfluidic chips. These sheets were then held in place by 3D-printed inserts. Seeded devices were placed in the incubator at 37°C for 15 min to facilitate hydrogel polymerization. 1 mg/mL mouse laminin (Sigma-Aldrich) was then pipetted into the microfluidic channels and allowed to coat for an additional 15 min at 37°C. After washing the microfluidic channels with EGM-2 (Lonza) to remove the laminin solution, BMVECs were pipetted into the channels at 20 million cells/mL and allowed to attach for 1 hr in the incubator with the devices turned upside down. 8 mL of EGM-2, supplemented with penicillin-streptomycin (Gibco), was then introduced to the media reservoirs from each impeller pump. Impeller rotation was started by placing the motor boxes on top of the devices inside an incubator. All devices started with an impeller speed of 300 RPM for the first 18 hrs to promote adhesion of BMVECs to the microfluidic channels. After 18 hrs, device kept at 300 RPM were referred to as “low flow” while devices increased to 900 RPM were referred to as “high flow”. Media was changed every other day by aspirating and replacing 4 mL of EGM-2.

### Compound Transport Comparison Principle

We devised the following metric to quantify transport of compounds from the circulating nutrient media into the interstitial space of the vascular bed compartment. Compound transport (*T*) was defined as compound flux (*J*) into the interstitial space, expressed as number of molecules (*N*) per unit time, normalized to the compound’s concentration in the nutrient media (*C_M_*).

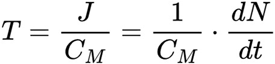

### Diffusion and Avascular Flow Measurements

4 kDa FITC-labeled dextran (Sigma-Aldrich) was added to the nutrient media reservoirs of VIVOS devices on Day 2 of culture at a final concentration of 1 mg/mL. For measuring diffusion, the impeller pump was turned on for 10 s at 900 RPM to pulse dextran into the microfluidic channels, followed by leaving the impeller pump off and time-lapse imaging the vascular bed compartment every 10 s. For measuring avascular flow, the same method was followed except the impeller pump was left on at 900 RPM. Vascular beds were imaged from below the microfluidic channels.

Entry of dextran into the vascular bed compartment can be described as two processes (**Extended Data Fig. 6C**). First, dextran starts in the microfluidic channel (which is at a concentration of *C_M_*), crosses the channel endothelial lining and porous membrane at a rate ***J***, and enters the hydrogel immediately above (which is at a concentration of *C_M_(t)*). Second, dextran diffuses towards the center of the vascular bed compartment to form a concentration gradient *C_B_(x,t)*. Given *N = CV*, the compound transport metric can be expressed as:

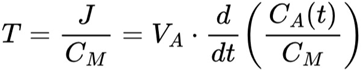

Here, *V_A_* refers to the volume of the hydrogel immediately above the porous membrane. Because dextran fluorescent intensity primarily comes from the imaging setup’s depth of field, the height of *V_A_* was approximated to be ∼0.1 mm.

*C_B_(x,t)* was measured from the dextran time-lapse imaging. We assumed that dextran fluorescent intensity was directly proportional to dextran concentration. To relate *C_A_(t)* to *C_B_(x,t)*, we used the analytical solution of Fick’s second law for diffusion past a semi-infinite boundary:

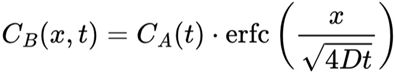

We treated *C_A_(t)* to be varying with time and applied nonlinear least squares curve fitting using SciPy in Python to obtain regression values for *C_A_(t)* and the diffusion coefficient (*D*). Here, the variable *x* refers to the perpendicular distance between a point within the vascular bed compartment and the edge of the microfluidic channel (described in **Extended Data Fig. 6B, C**).

Because regression diffusion coefficient values did not substantially differ between performing the experiment with the pump off or the pump on (**Extended Data Fig. 6F**), we did not extend the model to include convection. Moreover, this shows that of the two processes, the limiting factor in the system is dextran cross the channel endothelial coating and porous membrane.

The compound transport metric for diffusion was defined to be that of the pump off condition. The compound transport metric for avascular flow was defined to be the pump on condition minus the pump off condition.

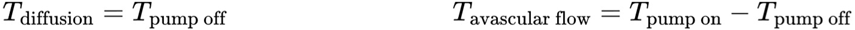

For an ad hoc comparison of avascular flow across different impeller speeds (ω), we assumed that avascular flow was directly proportional to the pressure differential (*Δ_P_*) across the vascular bed compartment (related to **Fig. 2E**).

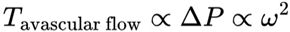

Analysis code for diffusion and avascular flow measurements can be found in the source data.

### Vascular Flow Physics Model

Given a compound’s concentration in the microfluidic channel (*C_M_*), vessel lumen (*C_V_*), and interstitial space (*C_I_*), compound movement into (*J_I_*) and out of (*J_O_*) the vessel lumen can be described using the total vessel flowrate (*Q_V_*) (**Extended Data Fig. 7A**):

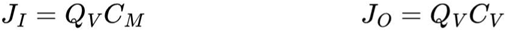

Movement across vessels is further described using vessel wall permeability (*P*) and vessel wall surface area (*S_V_*):

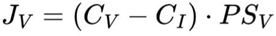

Compound transport can be described since *J_O_ = J_I_ - J_V_*. Assume that *C_I_ ≂ O*.

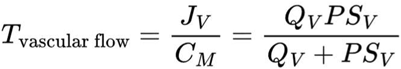

To solve this formula, we utilized the vessel permeability assay to calculate *P*, and tracking of fluorescent beads to calculate *Q_V_*.

### Vessel Permeability Assay

4 kDa or 7kDa FITC-labeled dextran (Sigma-Aldrich) was added to the nutrient media reservoirs of VIVOS devices on Day 4 of culture to a final concentration of 1 mg/mL. The impeller pump was turned on for 10 s at 900 RPM to pulse dextran into the vessels, followed by leaving the impeller pump off and time-lapse imaging the vascular bed compartment every 10 s for 20 min to track dextran efflux into the interstitial space. The first frame of the time-lapse imaging was used to segment the perfused vessels from the interstitial space.

For each time-lapse, ImageJ/FIJI was used to measure average fluorescent intensities over time for both vessel regions (*I_V_(t)*) and interstitial regions (*I_I_(t)*). Exponential regression was performed on these values using nonlinear least squares curve fitting in R:

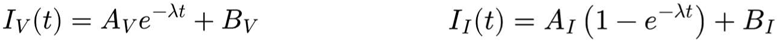

Here, *A_V_, B_V_, A_I_, B_I_*, and λ refer to coefficients from the exponential regression. Perfused vessel volume fraction (*α*) was calculated using the following:

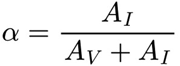

Vessel wall permeability (*P*) was calculated using the following:

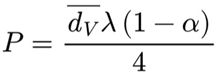

Here, *d̅_v_* refers to the average diameter of perfused vessels. Derivation of these formulas is presented in the **Supplementary Methods**. Analysis code can also be found in the source data.

### Tracking of Fluorescent Beads

1 µm diameter Fluoresbrite YG Microspheres (Polysciences) were added to the media reservoirs of VIVOS devices on Day 4 of culture to a final concentration of ∼10^8^ beads/mL. The impeller pump was turned on at 900 RPM followed by fast time-lapse imaging at 500 frames per second (FPS) for 20 seconds. Following denoising using NIS-Elements’ Denoise.AI, the resulting time-lapse sequences were analyzed using Python, scikit-image (v0.24.0), and OpenPIV (v0.25.3).

OpenPIV produces a velocity vector field (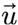) across the entire vessel, from which average velocities are calculated. To obtain wall shear stress, the following formula was used:

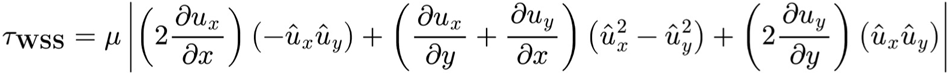

Here, *μ* is the fluid’s dynamic viscosity, *u_x_ & u_y_* are the velocity vector components, and 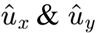 are their unit vector components. Derivation of this formula is presented in the **Supplementary Methods**. Analysis code can also be found in the source data.

### Bulk RNA-seq

Vascular beds, grown using RFP-labeled BMVECs, were collected on Day 1 (i.e. 24 hrs), Day 2, Day 4, or Day 8 using either low or high flow. All samples began with an impeller speed of 300 RPM for the first 18 hrs. Afterwards, devices using low flow were kept at 300 RPM while devices using high flow were increased to 900 RPM. On the day of collection, vascular beds were excised from microfluidic chips with a scalpel, washed 3 times with warm PBS, minced, and lysed with RLT buffer (Qiagen). The vascular bed compartments of the microfluidic chips were further rinsed with RLT buffer to dislodge remaining vascular bed fragments. Monolayer RFP-labeled BMVECs, NHLFs, and HBVPs were trypsinized, pelleted, and lysed with RLT buffer. RNA extraction was performed using RNeasy (Qiagen) and the resulting samples were processed at the Network Biology Collaborative Centre (NBCC). After TruSeq mRNA library prep (Illumina), Fragment Analyzer (Agilent) assessment showed all RNA quality numbers to be greater than 9. Following sequencing on a NextSeq 500 (Illumina), single-end reads were mapped to GRCh38 using STAR (v2.7.9a) with ∼15-40 million uniquely mapped reads per sample (**Supplementary Data 5**) ^85^. Gene counts were quantified using featureCounts (v2.0.3) ^86^. Aggregated counts data was normalized using DESeq2 (v1.40.2) with further analyses for gene differential expression ^87^.

### Vascular Bed Dissociation for scRNA-seq

Vascular beds, grown using RFP-labeled BMVECs, were collected on Day 1 (i.e. 24 hrs), Day 4 using low flow, or Day 4 using high flow. The impeller speed for both Day 1 and Day 4 low flow was 300 RPM for the entire duration. For Day 4 high flow, devices began at 300 RPM for the first 18 hrs and then switched to 900 RPM. On the day of collection, vascular beds were excised from microfluidic devices with a scalpel, washed 3 times with warm PBS, minced, and immersed in Accutase (Gibco) dissolved with 10 mg/mL Nattokinase (Japan Bio Science Laboratory). The vascular bed compartments of the microfluidic devices were further rinsed with Accutase to dislodge remaining vascular bed fragments. Samples from 4 separate devices were combined per condition and were incubated for 20 min on an orbital shaker at 37°C with mixing by pipetting up and down after 10 minutes. Twice, cell suspensions were passed through 35 µm cell strainers (Falcon) followed by centrifuging and resuspension in fresh EGM-2 media. Samples were processed at the Network Biology Collaborative Centre (NBCC) using the Chromium platform (single cell 3’ v3 chemistry; 10X Genomics), followed by sequencing of cDNA libraries on a NovaSeq 6000 (Illumina).

### scRNA-seq Preprocessing

Sequencing reads were processed using Cell Ranger (v7.1.0; 10X Genomics) and aligned to GRCh38, with ∼14k-17.5k cells/sample, and ∼35-45k mean reads/cell (**Supplementary Data 5**). Prior to creating a merged object in Seurat (v4.4.0) with all samples ^88^, velocyto (v0.17.16; Python version) was used to finding spliced and unspliced gene counts ^89^, SoupX (v1.6.2) was used to correct for ambient RNA contamination ^90^, and Souporcell (v2.5) was used to assign single nucleotide variant (SNV) profiles and remove doublets ^91^. All later analyses, with the exception of RNA velocity, were performed on the SoupX corrected counts. To identify which SNV profile belonged to which cell line (i.e. BMVECs, NHLFs, and HBVPs), BMVECs were assigned to the profile expressing endothelial markers and NHLFs were assigned to the larger of the remaining profiles because NHLFs and HBVPs were seeded on VIVOS at a 4:1 ratio. Low quality cells were excluded based on a mitochondrial gene percentage greater than 10% or a unique molecular identifier (UMI) count of less than 4000. After an initial unsupervised clustering of BMVECs, two clusters which contained 5.8% and 2.9% of total BMVECs were further removed. We excluded the first cluster because its top marker genes were primarily uncharacterized transcripts such as *AC007319.1*, *AC012668.3*, and *AP003086.1* (**Supplementary Data 6**), and thus was likely a technical artifact. We excluded the second cluster because its top marker genes were highly expressed in NHLFs and HBVPs, such as *COL1A1*, *COL3A1*, and *DCN* (**Supplementary Data 6**). Depending on the sample, Souporcell annotated ∼10-20% of total cells to be doublets, indicating that doublets were a significant technical issue. Given that the second cluster only represented ∼0.3-0.4% of total cells, we considered it more plausible that this cluster was composed of Souporcell false negative doublets, rather than composed of an endothelial-stromal hybrid cell type. Seurat R objects for the two excluded clusters can be found in the source data.

### scRNA-seq Cell Type Identification

After Souporcell unmixing, BMVECs, NHLFs, and HBVPs were clustered independently by following the method presented by Butler *et al*. ^92^. Briefly, anchor-based dataset integration in Seurat using “FindIntegrationAnchors()” and “IntegrateData()” was first applied to overlap cells from each experimental condition. Next, clusters shared between conditions were identified in Seurat using the integrated counts through PCA dimensionality reduction, shared nearest neighbor graph construction, and Louvain clustering ^93^. Lastly, cluster labels were transferred back to the non-integrated dataset. These clusters are hereafter referred to as cell types. All downstream analyses such as differential expression used the original, non-integrated counts.

### Human scRNA-seq Atlases

Integration of atlases was performed using anchor-based methods in Seurat. Prediction of cell types used “FindTransferAnchors()” and “MapQuery()” while generation of integrated counts used “FindIntegrationAnchors()” and “IntegrateData()”. scRNA-seq of adult human lung stromal cells was obtained from Adams *et al*. ^29^. “Myofibroblast”, “Pericyte”, “SMC” (i.e. smooth muscle cells), and “Fibroblast” clusters were used. scRNA-seq of fetal human lung stromal cells was obtained from He *et al*. ^30^. Cells from “Distal” and “Proximal” dissections were used while “Whole-lung” cells were excluded. Cells from “Normal” and “Trypsin+” chemistries were used while “Trypsin+CD326” cells were excluded. Mesothelial clusters were further excluded. scRNA-seq of human brain perivascular cells was obtained from Wälchli *et al*. ^31^. “Smooth muscle cells”, “Pericytes”, and “Fibroblasts” clusters from the unsorted adult control brain samples were used. scRNA-seq of human tumor angiogenesis was obtained from Goveia *et al*. ^33^.

### scRNA-seq Pseudobulk

scRNA-seq pseudobulk counts were calculated using “AggregateExpression()” in Seurat. We performed both scRNA-seq and bulk RNA-seq on vascular beds on Day 1 (i.e. 24 hrs) using an impeller speed of 300 RPM. To leverage this overlap between the scRNA-seq and bulk RNA-seq datasets, we applied the following formula to all pseudobulk counts in an effort to correct for technical differences between scRNA-seq pseudobulk counts and bulk RNA-seq counts:

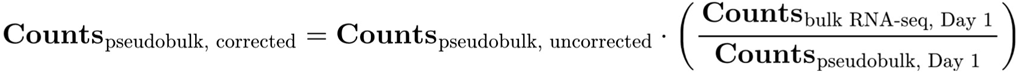

After this correction, all counts were additionally normalized using DESeq2 with a list of housekeeping genes serving as control genes for estimating size factors. The list of 20 most stable transcripts from the HRT Atlas v1.0 database was used as the housekeeping genes ^94^.

### scRNA-seq Downstream Analyses

Where noted, scRNA-seq counts were denoised using Markov Affinity-based Graph Imputation of Cells (MAGIC; v2.0.3.999; R version) with default parameters ^95^.

scRNA-seq gene set scoring was performed using “AddModuleScore()” in Seurat. For small gene sets (i.e. ≤ 10), scores were calculated on MAGIC-denoised expression values. All feature plots show scores calculated on non-denoised expression values. Specific information for each figure panel involving gene set scoring can be found in the source data. The gene set of YAP/TAZ targets involved *CTGF*, *CYR61*, *ANKRD1*, *AMOTL2*, *CCND1*, and *ANGPT2*. *ANGPT2* was included because it was previously characterized as a YAP/TAZ regulated gene in endothelial cells ^65,96,97^. Gene sets of VEGFA up/down-regulated genes were derived from bulk RNA-seq data of VEGFA-treated endothelial cells from Zhang *et al*. (GSE41166; **Supplementary Data 2**) ^32^. Raw counts (GSM1009635 to GSM1009638) were normalized using DESeq2 and filtered to include only genes with an average normalized expression greater than 100. Using the normalized counts, VEGFA up-regulated genes were defined has having a log2 fold change > 0.75 when comparing the 12 hr VEGFA-treated condition to the control condition. VEGFA down-regulated genes were similarly defined but with a log2 fold change < −0.75. The gene set of Apelin up-regulated genes was derived from bulk RNA-seq data of endothelial cells under flow treated with Apelin from Park *et al*. (GSE230549; **Supplementary Data 2**) ^57^. Differential expression was performed on raw counts (GSM7226664 to GSM7226669) using DESeq2. Only genes with total counts across all samples ≥ 10 were considered. Apelin up-regulated genes were defined as the top 100 genes, ranked by log2 fold change, which had an adjusted p-value of < 10^-6^. Gene sets of BMP9 up/down-regulated genes were obtained from bulk RNA-seq data of BMP9-treated human microvascular endothelial cells (HMVECs) from Al Tabosh *et al*. (**Supplementary Data 2**) ^64^. BMP9 up-regulated genes were defined as all up-regulated genes reported by Al Tabosh *et al*. (log2 fold change > 1 and adjusted p-value < 0.05). BMP9 down-regulated genes were defined as all down-regulated genes reported by Al Tabosh *et al*. (log2 fold change < −1 and adjusted p-value < 0.05). Curated tip cell and stalk cell marker genes are listed in **Extended Data Table 1**.

Gene Set Enrichment Analysis (GSEA) was performed using fgsea (v.1.26.0) with log2 fold changes as the ranking metric and a weight parameter of 2 ^98^. Only custom gene sets were utilized. Gene over-representation analysis (ORA) was performed using “gprofiler2” in R on default settings with Reactome and Gene Ontology Biological Process gene sets ^99–101^. Only gene sets smaller than 700 were considered.

Spliced and unspliced gene counts from velocyto were analyzed using scVelo with the dynamical model (v0.2.5) to obtain cell trajectories ^102^. We projected streamlines onto scatter plots involving expression of single genes (i.e. *APLN* and *APLNR*) as well as gene set scores for enhanced and more intuitive visualizations.

Differential expression and marker gene identification were performed using “FindMarkers()” and “FindAllMarkers()” in Seurat with the Wilcoxon rank sum test on counts normalized using SCTransform v2 ^103^. Marker genes for each cell type can be found in **Supplementary Data 1**. Differentially expressed genes between Day 1 and Day 4 for each cell type can be found in **Supplementary Data 3**. Differentially expressed genes between Day 4 low flow and Day 4 high flow for each cell type can be found in **Supplementary Data 4**.

To assess a gene’s co-expression with *APLN*, Spearman’s correlation coefficient was calculated in R between that gene’s expression values and *APLN* expression values. MAGIC-denoised expression values were utilized. Co-expression was assessed for all detected genes in the scRNA-seq dataset (**Supplementary Data 7**).

Cell-cell communication analysis was performed using CellChat (v2.0.0) with the “triMean” method when calculating communication probabilities ^104^. “Secreted Signaling” and “Cell-Cell Contact” receptor-ligand categories were used.

Coordinates for Uniform Manifold Approximation and Projection (UMAP) plots ^105^ and gene expression scatter plots were generated using Seurat and exported to Scanpy (v1.9.5) for creating figures^106^. Rank plots, box plots, and other plots were created using ggplot2 in R (v3.5.1) ^107^. Heatmaps were created using ComplexHeatmap (v2.16.0) ^108^.

### ChIP-seq Re-analysis

Endothelial ChIP-seq data of YAP (GSM4979550), TAZ (GSM4979551), and TEAD1 (GSM5989667) was obtained from Ong *et al*. (GSE163458) ^19^. Endothelial ChIP-seq data for H3K27ac and H3K4me3 was obtained from ENCODE (ENCFF145DLH, ENCFF722HTM, ENCFF379VLB, ENCFF161GMO) ^109,110^. Plots were created using IGV (v2.19.5) ^111^.

### Monolayer Endothelial Cell Inhibitor & Growth Factor Treatments

BMVECs were seeded in 6-well plates at 600k cells/well and allowed to attach for 24 hrs. IAG933 (Piramal Pharma Solutions, kindly provided by Methvin Isaac, Ontario Institute for Cancer Research; 2 µM) was added for 4, 24, or 48 hrs. Human recombinant BMP9 (R&D Systems; 10 ng/mL) and/or human recombinant VEGF-165 (Gibco; 100 ng/mL) were added for 48 hrs. The appropriate amount of solvent (e.g. DMSO) was added to non-treated wells.

### Monolayer Endothelial Shear Stress Assay

12 mm diameter #1.5 glass coverslips (Electron Microscopy Sciences) were arranged around the edges of 60 mm dishes. Hollow cylinders 12 mm in height, 29 mm in diameter, and 0.8 mm in thickness were placed at the center of the 60 mm dishes, keeping the coverslips in place. These cylinders were 3D printed out of polylactic acid (MatterHackers). 600k BMVECs were seeded onto the coverslips and allowed to attach for 24 hrs. Shear stress was applied by placing the assembly onto an orbital shaker (ThermoFisher; 9.5 mm orbit radius) at 90 RPM for 4 days.

For Apelin-17 treatment, human Apelin-17 (MedChemExpress; 10 µM) was additionally added for 30 min after the 4-day shear stress period. Shear stress (or absence thereof) was maintained during the 30 min. The appropriate amount of solvent (i.e. PBS) was added to non-treated dishes.

### Vascular Bed Inhibitor & Growth Factor Treatments

All inhibitors and growth factors were added to vascular beds immediately after device seeding and maintained through media changes. IAG933 (Piramal Pharma Solutions, kindly provided by Methvin Isaac, Ontario Institute for Cancer Research) was used at 2 µM, ML221 (MedChemExpress) was used at 10 µM, human recombinant BMP (R&D Systems) was used at 10 ng/mL, and Axitinib (MedChemExpress) was used at 100 nM. The appropriate amount of solvent (e.g. DMSO) was added to non-treated devices.

### shRNA Knockdowns

shRNA target sequences for *ALK1* and *ENG* were cloned into pLKO.1-puro lentiviral vectors (Addgene #8453), followed by lentivirus production as described previously ^112^ and infection of BMVECs with 8 µg/mL polybrene (Sigma-Aldrich). The control vector with a luciferase targeting sequence was a gift from Dr. Daniel Schramek. shRNA target sequences can be found in **Extended Data Table 2** and were obtained from The RNAi Consortium ^113^. Media was changed 24 hrs after infection and cells were selected with 2 µg/mL puromycin for 2 days. Selected cells were maintained in 1 µg/mL puromycin for a maximum of 3 additional passages prior to vascular bed formation on VIVOS.

### Quantitative Real-Time PCR

RNA was extracted from trypsinized and pelleted cells using RNeasy (Qiagen) followed by processing using High-Capacity cDNA Reverse Transcription (ThermoFisher). qPCR reactions were performed using Power SYBR Green (ThermoFisher) on a CFX Opus 384 (Bio-Rad). Primers were designed using PrimerQuest (Integrated DNA Technologies) to span exon-exon junctions, and ordered through ThermoFisher Scientific. Gene expression was calculated using the ΔΔCt method and utilized the average of *GAPDH* and *RPLP0* Ct values as the housekeeping reference. Primer sequences can be found in **Extended Data Table 3**. Relative expression and raw Ct values for all experiments are reported in the source data.

### Immunofluorescence

A list of antibody information can be found in **Extended Data Table 4**.

For whole mount immunostainings, vascular beds were excised from the microfluidic chip using a scalpel and fixed in 4% PFA (Electron Microscopy Sciences) for 12 hrs at 4°C. All steps were performed on a plate rocker. After 3 washes with PBS for 1 hr, vascular beds were blocked in 2% BSA Fraction V (Roche) + 0.5 % Triton X-100 (BioShop) for 1 hr at room temperature. Primary antibodies were diluted in blocking buffer and incubated for 12 hrs at 4°C. After 3 washes with blocking buffer for 1 hr at 4°C, secondary antibodies were diluted in blocking buffer and incubated for 12 hrs at 4°C. After 3 washes with blocking buffer for 1 hr at 4°C, vascular beds were cleared using the following modified SeeDB protocol ^114^. Vascular beds were incubated with 40% fructose (BioShop) for 1 hr at room temperature, followed by a 12 hr incubation in SeeDB saturated fructose solution.

For monolayer immunostainings, endothelial cells grown on 12 mm diameter #1.5 glass coverslips (Electron Microscopy Sciences) were fixed with 4% PFA (Electron Microscopy Sciences) for 15 min, washed once with PBS, and then blocked with 2% BSA Fraction V (Roche) + 0.3 % Triton X-100 (BioShop) for 30 min. Primary antibodies were diluted in blocking buffer and incubated for 12 hrs at 4°C. After 3 washes with PBS + 0.1% Tween-20 (BioShop), secondary antibodies were diluted in blocking buffer and incubated for 3 hrs at room temperature. After 3 washes with PBS + 0.1% Tween-20 (BioShop), coverslips were mounted using ProLong Gold Antifade (Invitrogen).

For immunostainings of vascular bed cryosections, vascular beds were excised from the microfluidic chip using a scalpel and fixed in 4% PFA (Electron Microscopy Sciences) for 12 hrs at 4°C. All steps were performed on a plate rocker. After 3 washes with PBS for 1 hr, vascular beds were immersed in 15% sucrose (Sigma-Aldrich) for 6 hrs at room temperature, followed by 30% sucrose for 12 hrs at 4°C. After embedding in Frozen Section Compound (VWR) and flash freezing on dry ice, vascular beds were sectioned at 14 µm using a Leica CM3050 S. The remaining steps of blocking and applying antibodies are identical to those for the monolayer immunostainings except that blocking was performed for 1 hr.

### Imaging

All imaging was performed on Nikon ECLIPSE Ti2-E microscopes with various scan head and detector setups. Fluorescent dextran, fluorescent beads, fixed monolayer endothelial cells, and fixed whole mount vascular beds were imaged using a CrestOptics X-Light V3 spinning disk confocal. Non-fixed vascular beds were imaged using a CrestOptics X-Light V2 spinning disk confocal. Fixed vascular beds with implanted 3D tissues were imaged either with a CrestOptics X-Light V3 spinning disk confocal, or a Nikon A1R-MP laser scanning confocal. Structured illumination imaging of Apelin was performed using CrestOptics’s DeepSIM.

Where noted in the source data, images were deconvolved using the Richarson-Lucy algorithm and/or denoised using Denoise.AI in NIS-Elements. For stitching of fluorescent dextran images, flat-field correction was performed using BaSiC (12 Jan 2017 release) ^115^. Zoom-in, and 3D view images of cerebral organoids were processed using CARE isotropic reconstruction ^116^. The 3D vessel reconstruction was created using Imaris (v10.2.0).

### Neural Network Image Enhancement of RFP-labeled Vessels

Vascular beds grown from RFP-labeled endothelial cells were fixed and stained with CD31. Image-to-image convolutional neural networks were trained using BiaPy (v3.5.12; GUI v1.1.8) to predict CD31 images from RFP images ^117^. These neural networks were applied to vascular beds grown from RFP-labeled endothelial cells when CD31 staining wasn’t possible due to host-species incompatibilities with other antibodies. RFP signal, while strong, was punctated throughout vessels. The same imaging setup was utilized for both training and prediction of the neural networks. No quantification was performed on BiaPy-processed images. Corresponding non-processed image files are provided in the source data.

### Vessel Metrics Quantification

Vessel metrics quantification utilized either RFP-labeled vessels or vessels immunostained for CD31. Maximum intensity projections of multiple adjacent imaging planes, spaced 5 µm apart, were utilized. Vessels were first segmented using Phansalkar’s adaptive local thresholding method ^118^ in ImageJ. We found this simple algorithmic method accurately segmented vessels, with manual touch ups only required when imaging quality was suboptimal. Binaries were next analyzed using scikit-image (v0.24.0) in Python ^119^. After removing small objects and performing morphological closing, medial axis skeletons and distance transforms were calculated on the cleaned binaries. The skeletons were used to calculate vessel length densities and branchpoint densities, with a branchpoint defined as a pixel on the skeleton with 3 or more 8-connectivity positive neighbors. The skeletons, in conjunction with the distance transforms, were used to calculate mean diameters for each vessel segment. Code for this analysis pipeline is available in the source data.

### Immunofluorescence Quantifications

To quantify the number of KLF4+ endothelial nuclei in whole mount vascular beds, Phansalkar’s adaptive local thresholding method ^118^ in ImageJ was used to segment RFP-labeled vessels and StarDist (v0.3.0; ImageJ plugin) ^120^ was used to segment nuclei from DAPI images. Nuclei which were enclosed by the RFP labeling were assigned to be KLF4+ if its KLF4 mean intensity z-score was > 1. Maximum intensity projections of 3 adjacent imaging planes, spaced 5 µm apart, were used.

To quantify nuclear and cytoplasmic TAZ intensities in monolayer endothelial cells, CellProfiler (v4.2.8) ^121^ was first used to segment nuclei from DAPI images using adaptive Otsu’s method. “Expand objects until touching” under “ExpandOrShrinkObjects” was then applied to segmented nuclei in order to define cell boundaries. All images were of confluent endothelial cells. Nuclear and cytoplasmic mean intensities of TAZ were normalized to each cell’s DAPI mean intensity. Maximum intensity projections were used. Apelin whole-cell max intensities in monolayer endothelial cells were similarly measured using CellProfiler with the modification that max intensities were calculated throughout the entire cell.

To quantify TAZ whole-vessel mean intensities in vascular bed cryosections, CellProfiler was first used to segment RFP-labeled vessels using adaptive minimum cross-entropy. Next, TAZ mean intensities within entire vessels were measured followed by subtraction of background intensities.

To quantify the number of endothelial nuclei in wholemount vascular beds, StarDist was used to segment ERG images for each imaging plane, followed by counting the number of ERG objects. Each ERG+ nucleus was additionally assigned to be MKI67+ if its MKI67 mean intensity z-score was > 7. For both ERG and MKI67 quantifications, reported values are averages from 10 imaging planes, spaced 5 µm apart.

### Statistics

Description of statistical tests as well as numbers of biological and technical replicates are listed in the corresponding figure legends and source data. R was used to perform t-tests. For qPCR experiments, t-tests were performed on log2-transformed relative expression values. For experiments involving large numbers of data points such as scRNA-seq gene set scoring and immunofluorescence quantification of monolayer endothelial cells, we calculated the effect size metric Cohen’s d using the “effectsize” package in R ^122^.

## Acknowledgments

We thank D. Trcka, K. Chan, R. Padilla, L. Caldwell, and A. Obersterescu at the Network Biology Collaborative Center (NBCC) for scRNA-seq and bulk RNA-seq processing, and L. Caldwell for help with GEO submission. We thank M. Gerrie, R. Devi, M.Z. Li, and M. Yacoub for help with assembling VIVOS devices, L. Wang, J.J. Hernandez and J. Nurtanto for help establishing organoid cultures, and M. Bourmoum for help with lentivirus. We thank D. Schramek & Y.Q. Lü for providing the pLKO.1-puro control plasmid and MCF10A cells, and M. Issac for providing IAG933. Lastly, we thank T.-H. Kim, Y. Aghazadeh, I. Scott, and members of the Pelletier, Wrana, and Attisano labs for their feedback and discussions.

## Funding

This work was supported by the Lunenfeld-Tanenbaum Research Institute’s Network Biology Collaborative Centre, which is funded by the Canada Foundation for Innovation, the Ontario Government, and Genome Canada and Ontario Genomics (OGI-139) and the Nikon Center of Excellence at the Lunenfeld-Tanenbaum Research Institute. This work has been made possible with the financial support of Health Canada, through the Canada Brain Research Fund, an innovative partnership between the Government of Canada (through Health Canada) and Brain Canada, and of the Krembil Foundation. This project also received financial support from the Canadian Institutes of Health Research (CIHR) and the Natural Sciences and Engineering Research Council of Canada (NSERC) through a Collaborative Health Research Project (CHRP, 549533-20), as well as CIHR grants awarded to L.A. (FRN-148455), L.P. (FRN-167279), and J.W. (FRN-143252). L.P. is a Tier 1 Canada Research Chair in Centrosome Biogenesis and Function.

## Author contributions

T.H.Z.J., A.A.S., L.P., J.L.W, and L.A. co-conceived the study. T.H.Z.J. and A.A.S. designed and built the system with L.P., J.L.W., and L.A. providing input. T.H.Z.J. ran experiments, performed physics modeling, developed the vessel measurement assays, analyzed the sequencing data, and made the figures. Y.E.G. grew spheroids, A.A.S., C.J.S, Y.T, M.M, and J.T.G grew organoids, and S.L. cultured the retinal explants. Y.E.G., A.A.S., and C.J.S. additionally ran VIVOS experiments. A.Y. analyzed bead perfusion videos. J.M.T helped with VIVOS experiments involving shRNA knockdowns. L.P., J.L.W, L.A., and R.B. acquired funding and supervised the project. T.H.Z.J., J.L.W, L.A., and L.P. wrote the manuscript with feedback from all authors.

## Competing interests

T.H.Z.J., A.A.S., L.P., J.L.W, and L.A have submitted US provisional and PCT patent applications related to VIVOS.

## Data, code, and materials availability

Bulk RNA-seq and scRNA-seq files are available at the Gene Expression Omnibus (GEO) (accession GSE318597). Description of scRNA-seq files, source data for all figures and movies, as well as new code generated are available at Zenodo (https://doi.org/10.5281/zenodo.18529053). Please contact L.P., J.L.W, and L.A for material requests.

---

**Extended Data Fig. 1.**
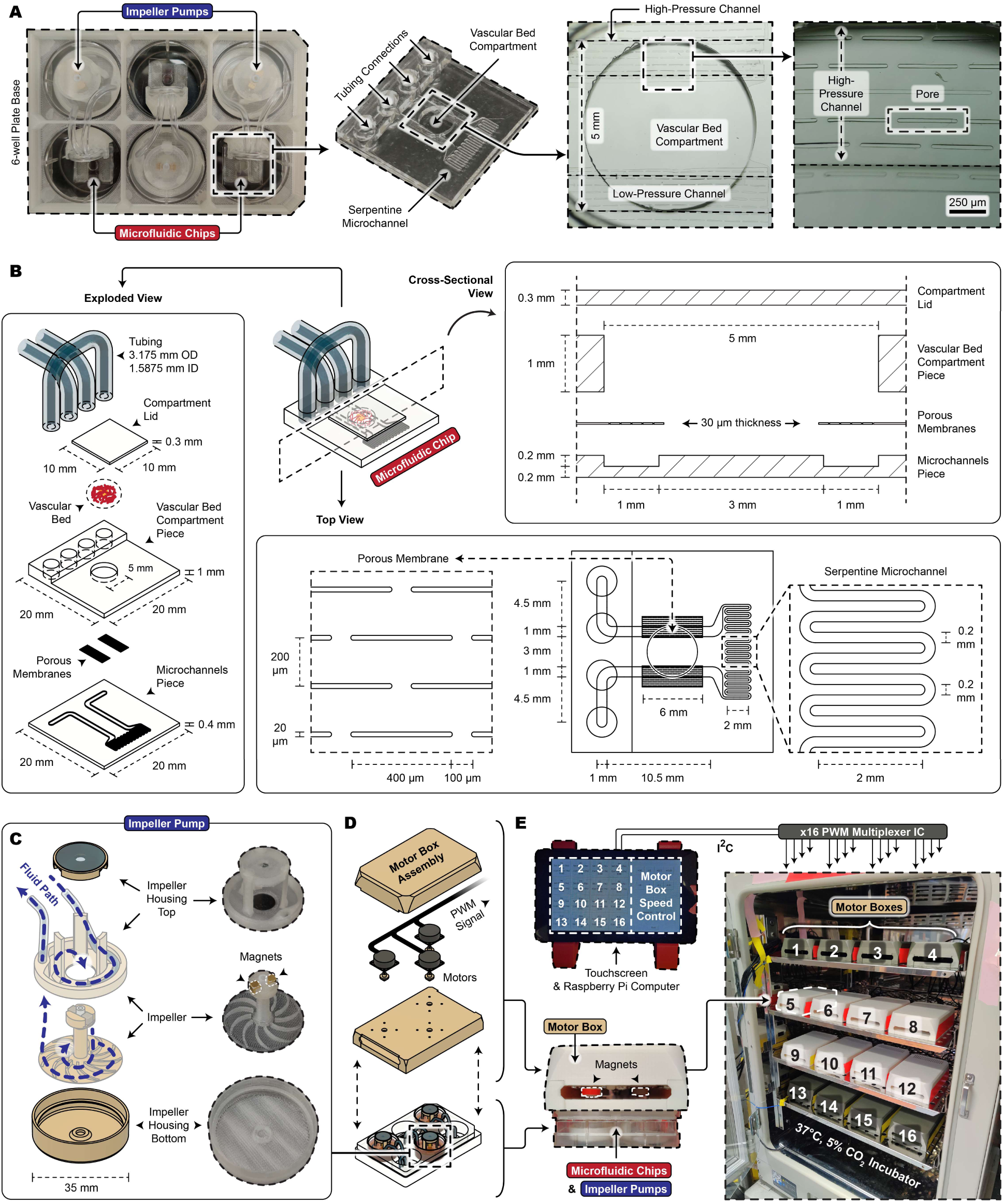
Technical schematics for the microfluidic chip and impeller pump of VIVOS. **(A)** Pictures of the microfluidic chips and impeller pumps situated within the 6-well plate. Zoomed-in pictures highlight the serpentine microchannel, high-pressure channel, low-pressure channel, vascular bed compartment, and pores of the porous membrane. **(B)** Exploded, cross-sectional, and top views of the microfluidic chip depicting the dimensions of individual components. **(C)** Exploded view of the 3D-printed impeller pump. **(D)** Exploded view of the motor box, which houses 3 motors each. **(E)** Pictures of a 48-unit VIVOS incubator containing 16 individually programmable motor boxes, with each motor box actuating a 6-well plate containing 3 microfluidic chips and 3 impeller pumps.

**Extended Data Fig. 2.**
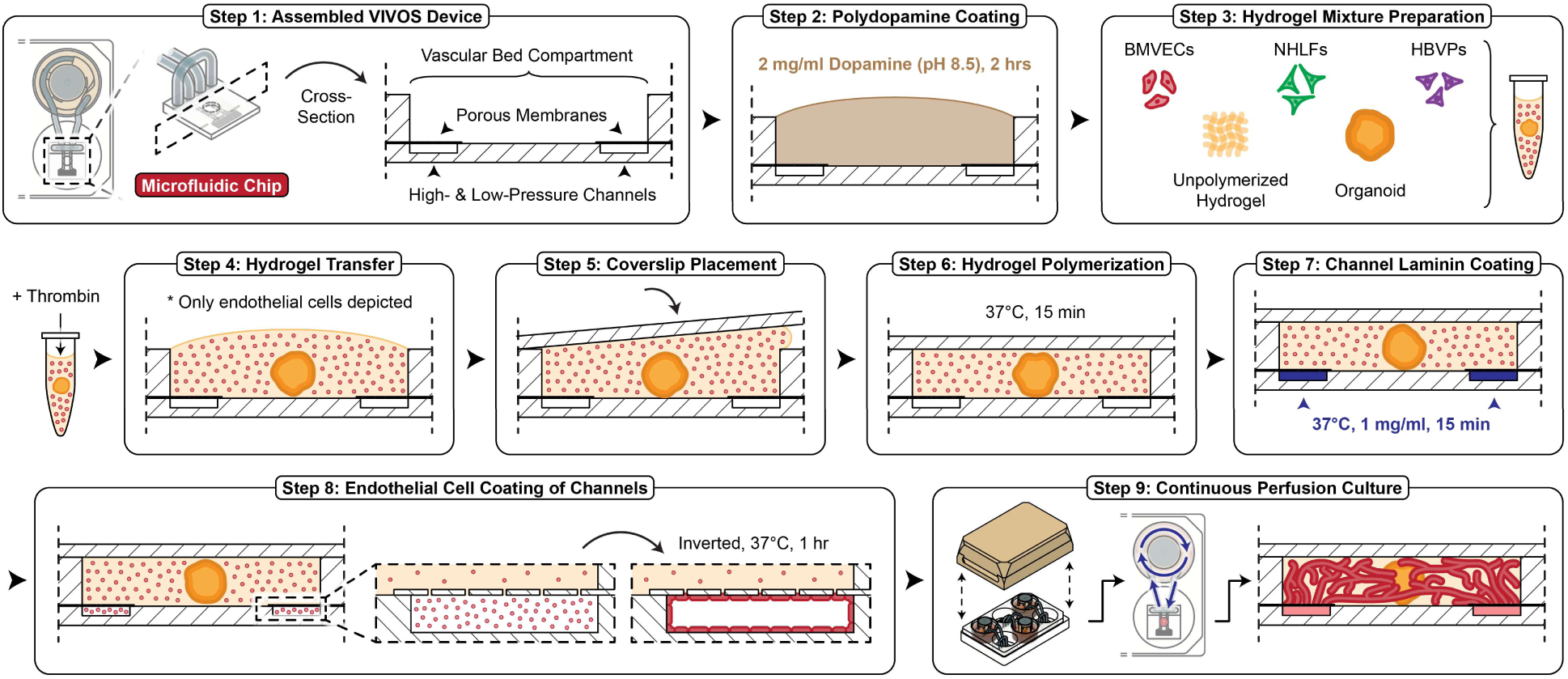
VIVOS seeding process. Illustrations showing the steps for initiating vascular beds on VIVOS devices. BMVECs: brain microvascular endothelial cells. NHLFs: normal human lung fibroblasts. HBVPs: human brain vascular pericytes.

**Extended Data Fig. 3.**
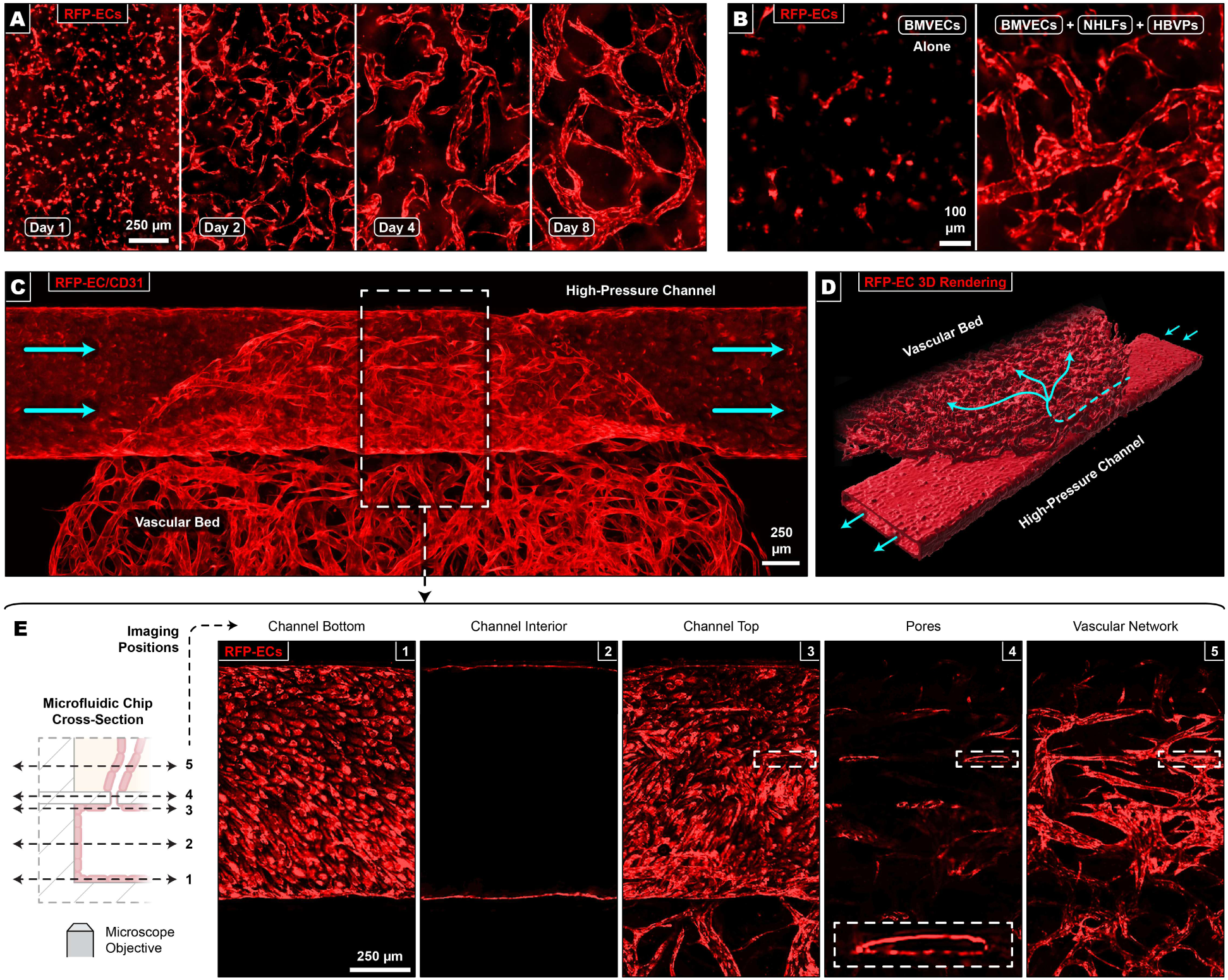
Formation of perfusable vascular beds on VIVOS. **(A)** Timecourse of vascular beds grown from RFP-labeled endothelial cells. **(B)** VIVOS devices on Day 4 seeded with either endothelial cells alone (BMVECs), or together with stromal cells (BMVECs + NHLFs + HBVPs). **(C)** Maximum intensity projection image of the endothelial-lined high-pressure channel interfacing with the vascular bed. Imaging of RFP-labeled vessels was enhanced with a BiaPy RFP-to-CD31 deep learning model. Cyan arrows depict fluid flow direction. **(D)** Imaris 3D rendering of (C). Cyan arrows depict fluid flow direction. **(E)** Individual imaging positions from (C) which show the endothelial cell lining of the high-pressure channel connecting with the vascular bed through the pores of the porous membrane.

**Extended Data Fig. 4.**
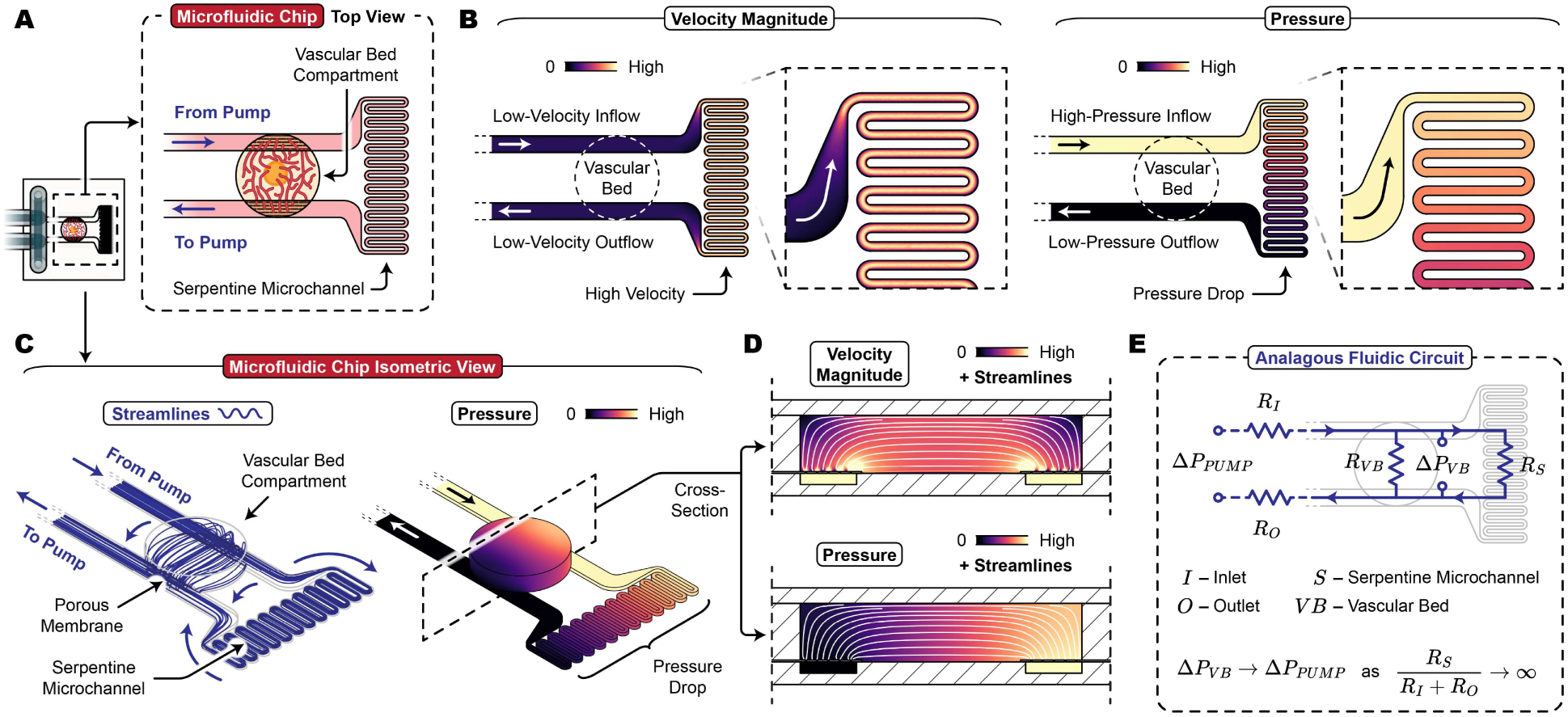
Physics of the microfluidic chip from VIVOS. **(A)** Top view of the microfluidic chip depicting the serpentine microchannel and vascular bed compartment. **(B)** Top view of computational fluid dynamics (CFD) simulations depicting velocity magnitude (left) and pressure (right) within the microfluidic channels. **(C)** Isometric view of CFD simulations depicting flow streamlines (left) and pressure (right) within the entire microfluidic chip. **(D)** Cross-sectional view of CFD simulations depicting velocity magnitude (top) and pressure (bottom) within the vascular bed compartment. **(E)** Fluidic circuit analogy of the microfluidic chip. The pressure gradient across the vascular bed compartment (*Δ*P*_VB_*) is maximized and approaches the output pressure of the impeller pump (*Δ*P*_PUMP_*) when the flow resistance of the serpentine microchannel (**R*_S_*) is significantly greater than the combined flow resistances of the rest of the fluidic circuit (**R*_I_ + *R*_O_*).

**Extended Data Fig. 5.**
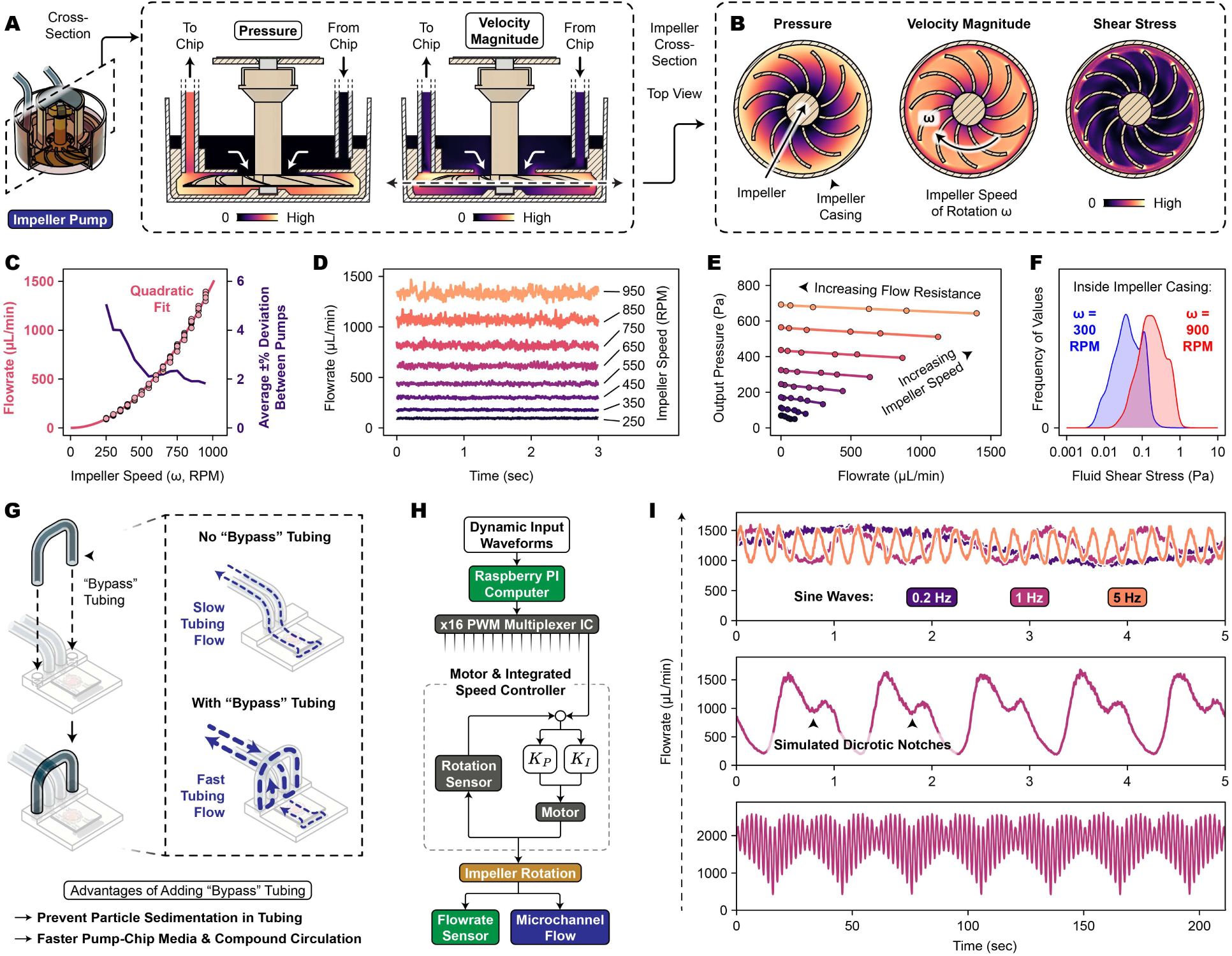
Physics of the impeller pump from VIVOS. **(A)** Cross-sectional view of computational fluid dynamics (CFD) simulations depicting pressure (left) and velocity magnitude (right) within the impeller pump. **(B)** Cross-sectional view of CFD simulations depicting pressure (left), velocity magnitude (center), and shear stress (right) within the impeller pump’s casing. **(C)** Relationship between impeller speed and measured flowrate. The average deviation between measurements from different impeller pump units is also shown. Measurements are from 6 impeller pump units. **(D)** Flowrate measurements over time for various impeller speeds. **(E)** Relationship between the impeller pump’s output pressure and measured flowrate for various impeller speeds and various flow resistances. Flow resistance refers to how the fluid path between the impeller pump’s inlet and outlet (i.e. through the microfluidic chip) resists flow. While keeping the impeller pump’s speed constant, flow resistances were manually varied by connecting progressively narrower tubing between the impeller pump’s inlet and outlet. **(F)** Frequency distribution of CFD-derived fluid shear stress values within the impeller pump’s casing. **(G)** Overview of a secondary “bypass” tubing which increases tubing flow between the microfluidic chip and impeller pump. **(H)** Schematic for the motor control mechanism which produces consistent and dynamically tunable flow. **(I)** Flowrate measurements over time for various dynamic waveforms.

**Extended Data Fig. 6.**
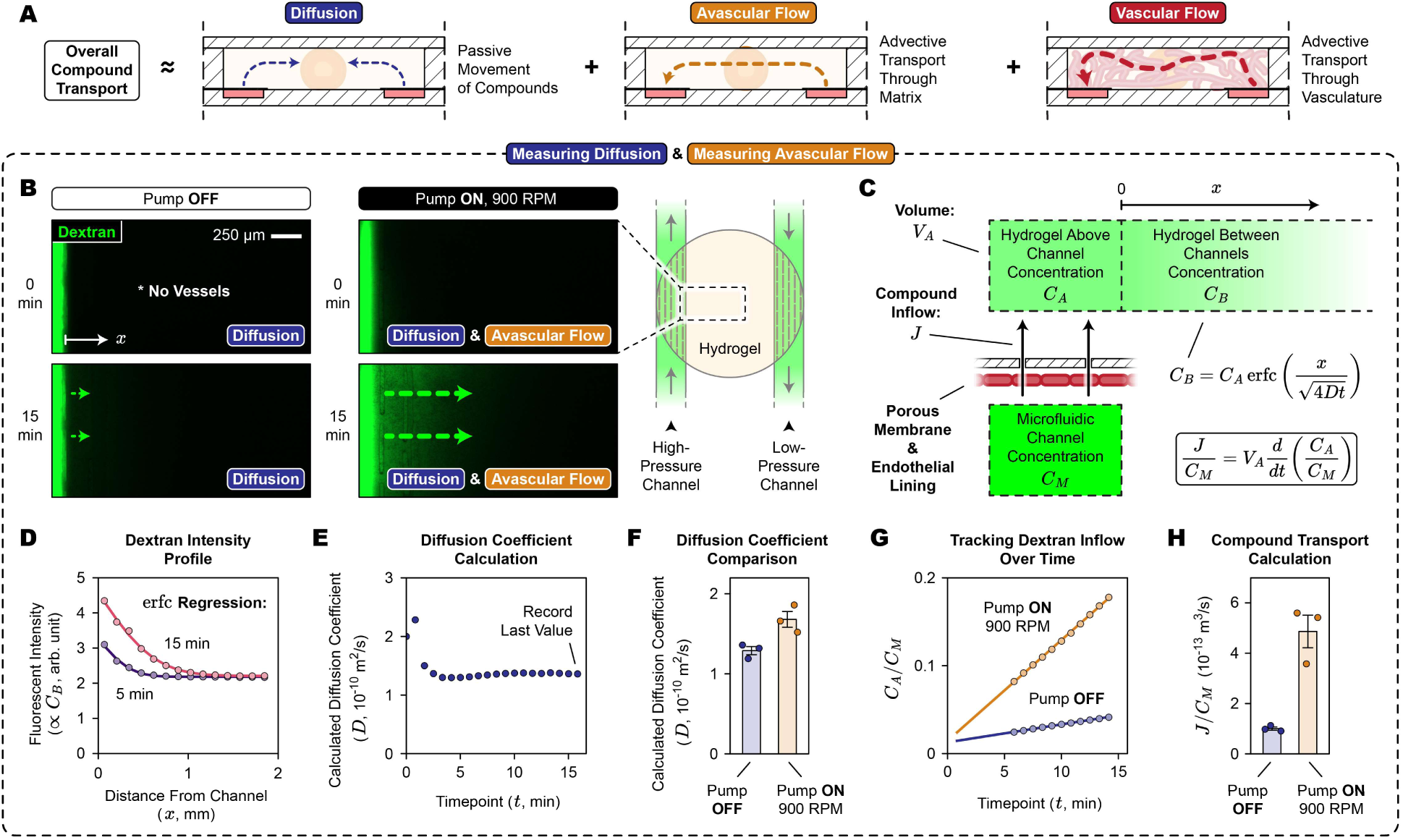
Measuring diffusion and avascular flow on VIVOS. **(A)** Overall compound transport is approximated as the sum between diffusion, avascular flow, and vascular flow. **(B)** Experimental setup for measuring diffusion and avascular flow. 4 kDa FITC-labeled dextran is pulsed into the microfluidic chip, followed by tracking of dextran movement after either turning the pump off or leaving the pump on at 900 RPM. Turning the pump off measures diffusion while leaving the pump on measures both diffusion and avascular flow. Microfluidic chips contain the standard mixture of cells and hydrogel on Day 2 of culture when perfused vessels are absent. **(C)** Mathematical model of (B). The variable *D* refers to the dextran’s diffusion coefficient. **(D)** Dextran fluorescent intensity within the vascular bed compartment measured at different distances from the channel boundary. Example shows pump-on values. **(E)** Diffusion coefficients obtained through applying nonlinear least squares regression to each timepoint. **(F)** Pump-off and pump-on diffusion coefficient regression values (3 devices per condition). **(G)** Ratio between dextran concentration in the vascular bed compartment immediately above the channel (**C*_A_*) and dextran concentration within the channel itself (**C*_M_*), over time. **(H)** Numerical values for dextran inflow into the vascular bed compartment (3 devices per condition). Inflow from diffusion equals pump-off values. Inflow from avascular flow equals pump-on values minus pump-off values. **(F, H)** Error bars show SEM.

**Extended Data Fig. 7.**
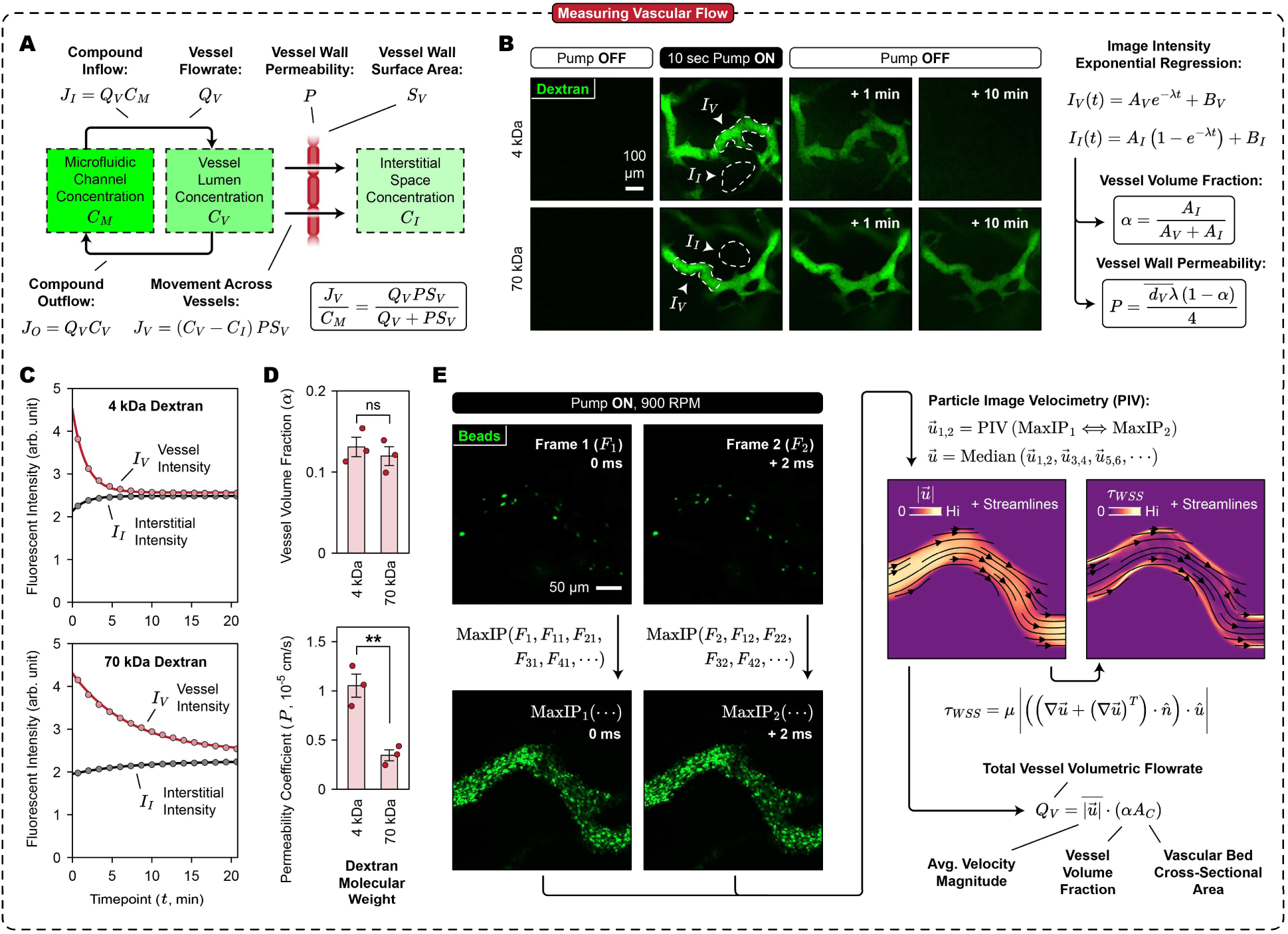
Measuring vascular flow on VIVOS. **(A)** Mathematical model for describing dextran entering the interstitial space of the vascular bed compartment through vascular flow. **(B)** Experimental setup for measuring vessel wall permeability. 4 kDa or 70 kDa FITC-labeled dextran is pulsed into perfused vessels, followed by tracking of dextran fluorescent intensities at vessel (**I*_V_*) and interstitial (**I*_I_*) regions over time. Vessel wall permeability (P), as well as vessel volume fraction (α), is derived from exponential regression coefficients (*A_V_, B_V_, A_I_, B_I_, λ*). **(C)** Dextran fluorescent intensities at vessel and interstitial regions over time for 4 kDa or 70 kDa dextran. **(D)** Calculated vessel volume fractions and vessel wall permeabilities for 4 kDa or 70 kDa dextran. Vessel volume fraction refers to the fraction of the vascular bed compartment that is occupied by perfused vessels. 2-tailed unpaired student’s t-test (ns: not significant; ** p < 0.01). Error bars show SEM. **(E)** Experimental setup for measuring vessel flow velocities, vessel wall shear stresses, and vessel volumetric flowrates. 1 µm diameter fluorescent beads are continuously perfused through vessels. Fast time-lapse imaging coupled with particle image velocimetry (PIV) produces velocity and shear stress fields across the entire field of view. Volumetric flowrates are further calculated by incorporating vessel volume fractions (α) and the cross-sectional area of the vascular bed compartment.

**Extended Data Fig. 8.**
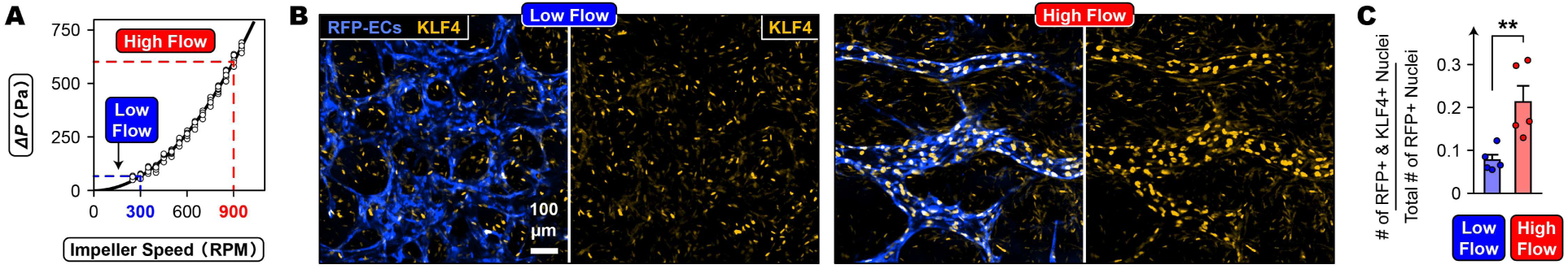
Establishment of low flow and high flow experimental conditions on VIVOS. **(A)** Relationship between impeller speed and pump output pressure (*Δ*P**). Low flow and high flow correspond to the impeller pump set to 300 and 900 RPM, respectively. **(B)** KLF4 immunostaining of Day 4 vascular beds grown using either low or high flow. Vessels were from RFP-labeled endothelial cells. **(C)** Quantification of (B). Numerical values are of the number of nuclei double positive for RFP and KLF4, divided by the total number of RFP positive nuclei (5 devices per condition across 3 independent experiments). 2-tailed unpaired student’s t-test (** p < 0.01). Error bars show SEM.

**Extended Data Fig. 9.**
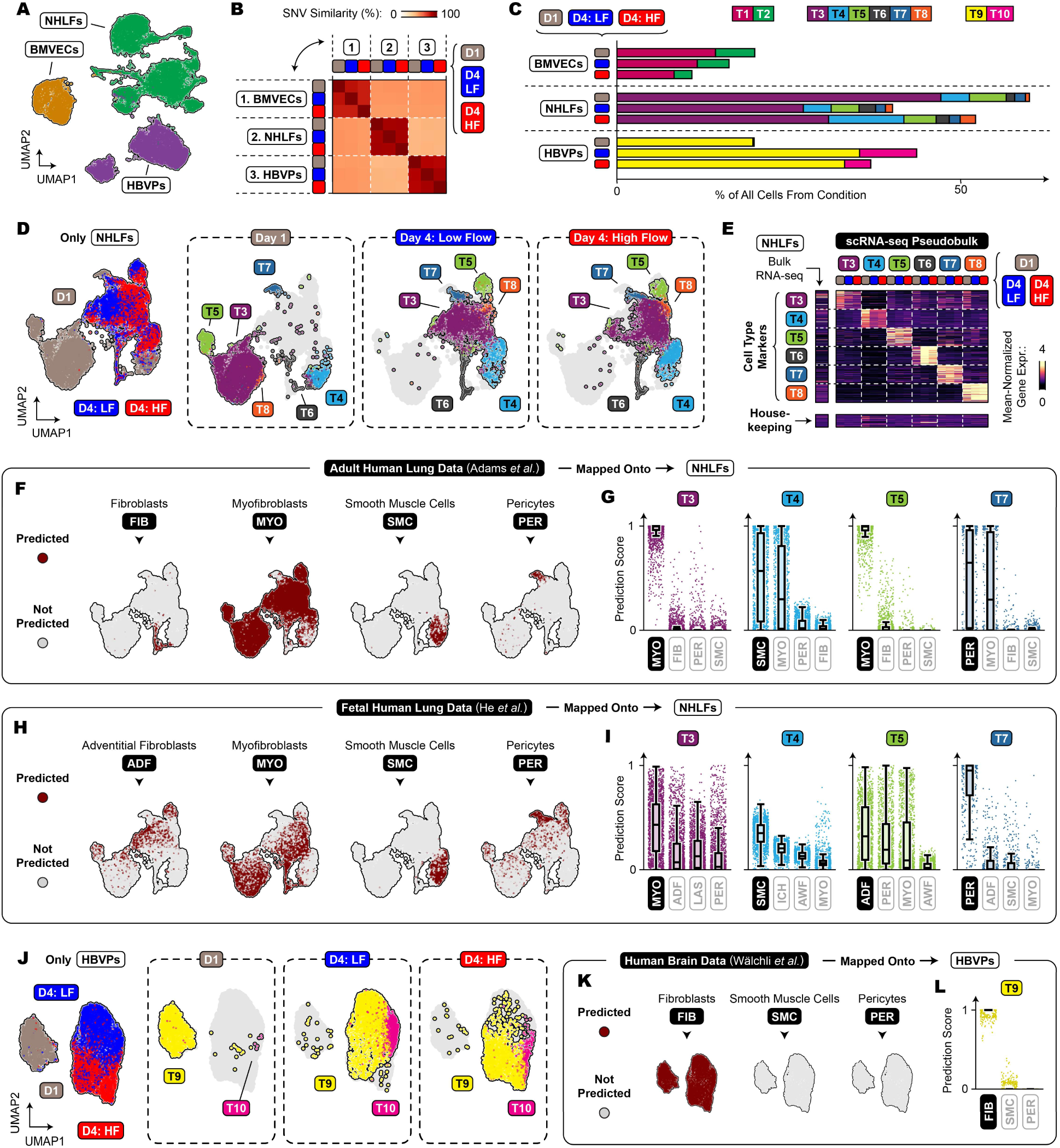
scRNA-seq characterization of stromal cell types on VIVOS. **(A)** Uniform Manifold Approximation and Projection (UMAP) plot of all cells. Origin cell lines (i.e. BMVECs, NHLFs, HBVPs) were identified using single nucleotide variant (SNV) based unmixing. **(B)** SNV similarity heatmap. **(C)** Relative proportions of identified cell types (i.e. T1-T10). **(D)** UMAP plots of cells originating from NHLFs, showing experimental conditions (left) and cell types (3 on the right). **(E)** scRNA-seq pseudobulk expression heatmap of marker genes from each NHLF cell type, shown alongside bulk RNA-seq data of the NHLF cell line grown in monolayer. **(F)** UMAP plots of predicted annotations from mapping adult human lung scRNA-seq data (Adams *et al*., ^29^) onto NHLFs. **(G)** Prediction scores from (F) for individual NHLF cell types. **(H)** UMAP plots of predicted annotations from mapping fetal human lung scRNA-seq data (He *et al*., ^30^) onto NHLFs. **(I)** Prediction scores from (H) for individual NHLF cell types. LAS: late airway SMC. ICH: intermediate chondrocyte. AWF: airway fibroblast. **(J)** UMAP plot of cells originating from HBVPs, showing experimental conditions (left) and cell types (3 on the right). **(K)** UMAP plots of predicted annotations from mapping adult human brain scRNA-seq data (Wälchli *et al*., ^31^) onto HBVPs. **(L)** Prediction scores from (K) for the T9 cell type.

**Extended Data Fig. 10.**
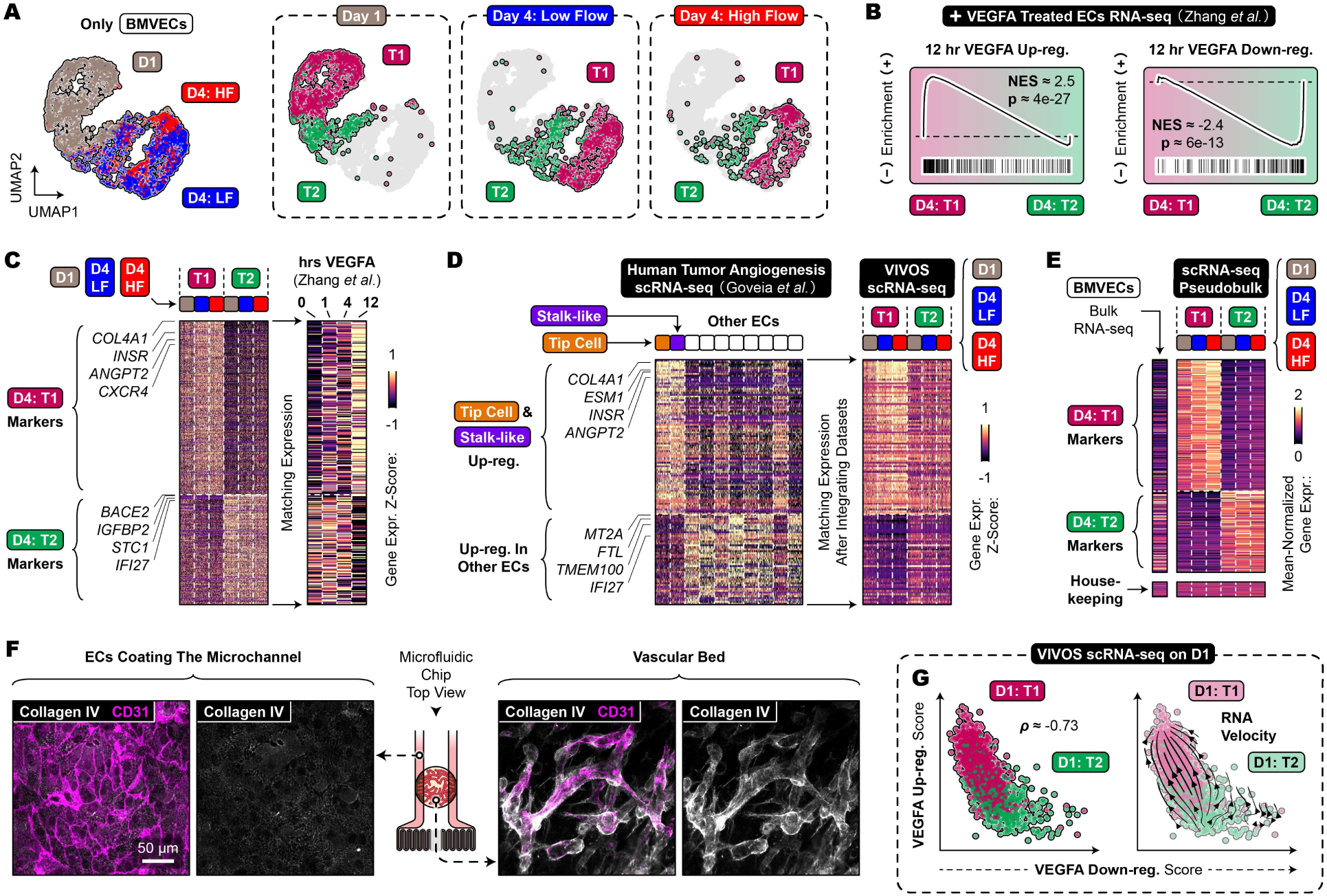
scRNA-seq characterization of endothelial cell types on VIVOS. **(A)** Uniform Manifold Approximation and Projection plot of cells originating from BMVECs, showing experimental conditions (left) and cell types (3 on the right). **(B)** Gene Set Enrichment Analysis of differentially expressed genes (DEGs) between T1 and T2 cell types on Day 4 for VEGFA up/down-regulated genes (Zhang *et al*., ^32^). **(C)** Expression heatmap of marker genes from T1 and T2 cell types on Day 4. Matching gene expression from bulk RNA-seq data of VEGFA-treated endothelial cells is shown (Zhang *et al*.). **(D)** Dataset integration of T1 and T2 cell types with the human tumor angiogenesis atlas from Goveia *et al*. ^33^. Heatmap depicts expression of DEGs from tip cell and stalk-like human atlas angiogenic clusters (left). Matching gene expression from T1 and T2 cell types is shown (right). **(E)** scRNA-seq pseudobulk expression heatmap of marker genes from T1 and T2 cell types on Day 4, shown alongside bulk RNA-seq data of the BMVEC cell line grown in monolayer. **(F)** Microfluidic chips on Day 2 stained for type IV collagen. Images are from either the endothelial cells lining the channels (left) or the vascular bed (right). **(G)** Gene set scoring of T1 and T2 cell types on Day 1 for VEGFA up/down-regulated genes (Zhang *et al*.). Spearman’s correlation coefficient (left). RNA velocity streamlines (right).

**Extended Data Fig. 11.**
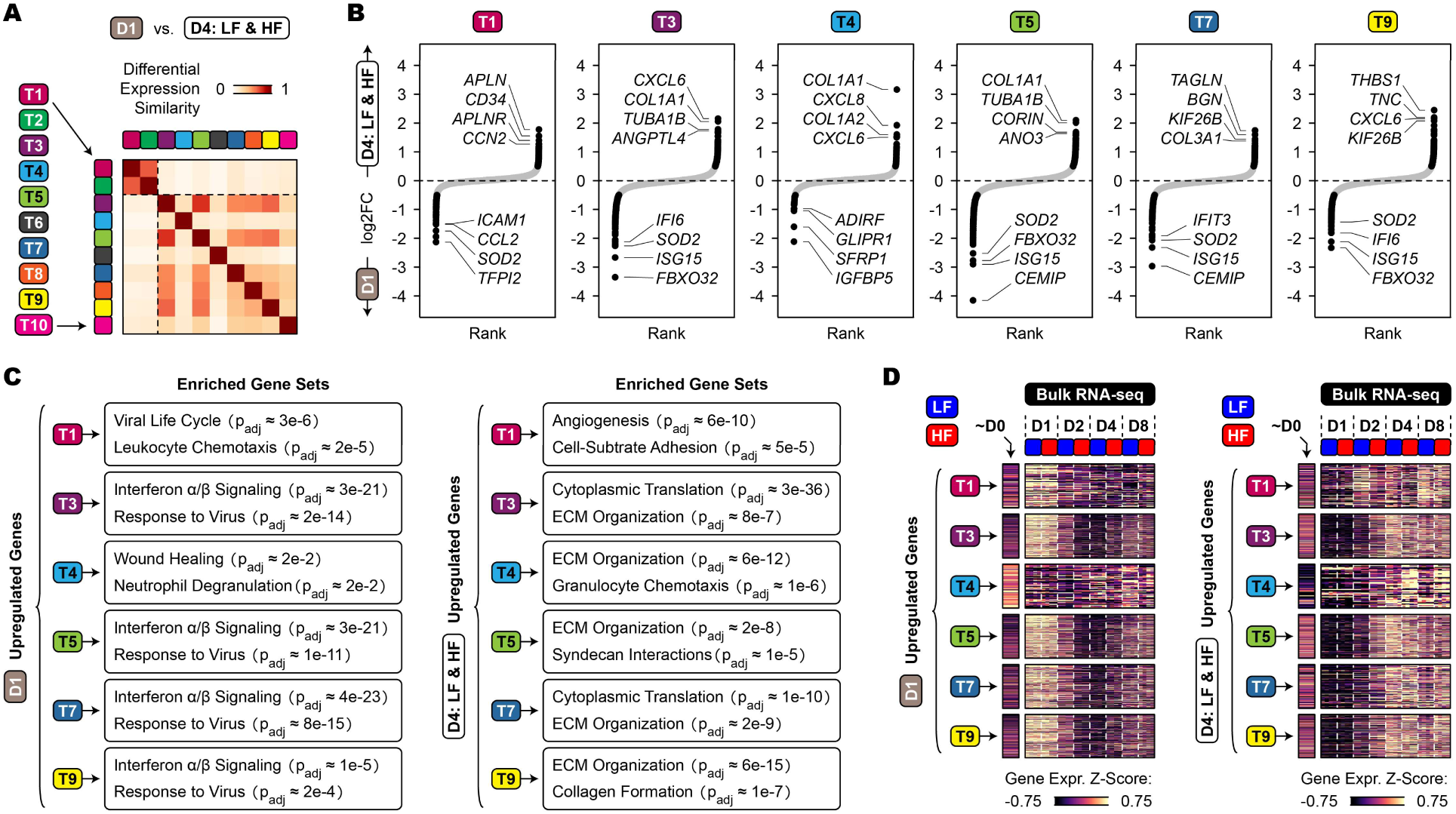
Transcriptomic differences during vessel formation. **(A)** Heatmap of cosine similarities between lists of differentially expressed genes (DEGs) for each cell type. DEGs were between Day 1 and Day 4. **(B)** Log2 fold change rank plots of DEGs between Day 1 and Day 4. **(C)** Enriched gene sets for Day 1 up-regulated genes (left) and Day 4 up-regulated genes (right). Over-representation analysis was performed using gProfiler with GO Biological Process and Reactome gene sets. **(D)** Expression heatmaps depicting bulk RNA-seq data of vascular beds grown using low or high flow at multiple timepoints. Genes on the heatmap are Day 1 up-regulated genes identified from scRNA-seq (left), and Day 4 up-regulated genes identified from scRNA-seq (right). “∼Day 0” (∼D0) refers to the average bulk RNA-seq expression from BMVECs, NHLFs, and HBVPs grown in monolayer, weighted by the ratio in which the 3 cell lines were mixed when seeded onto VIVOS. **(B-D)** Highlighted genes in (B) and genes used for analysis in (C) and (D) have an adjusted p-value < 0.01 and log2 fold change magnitude > 0.5.

**Extended Data Fig. 12.**
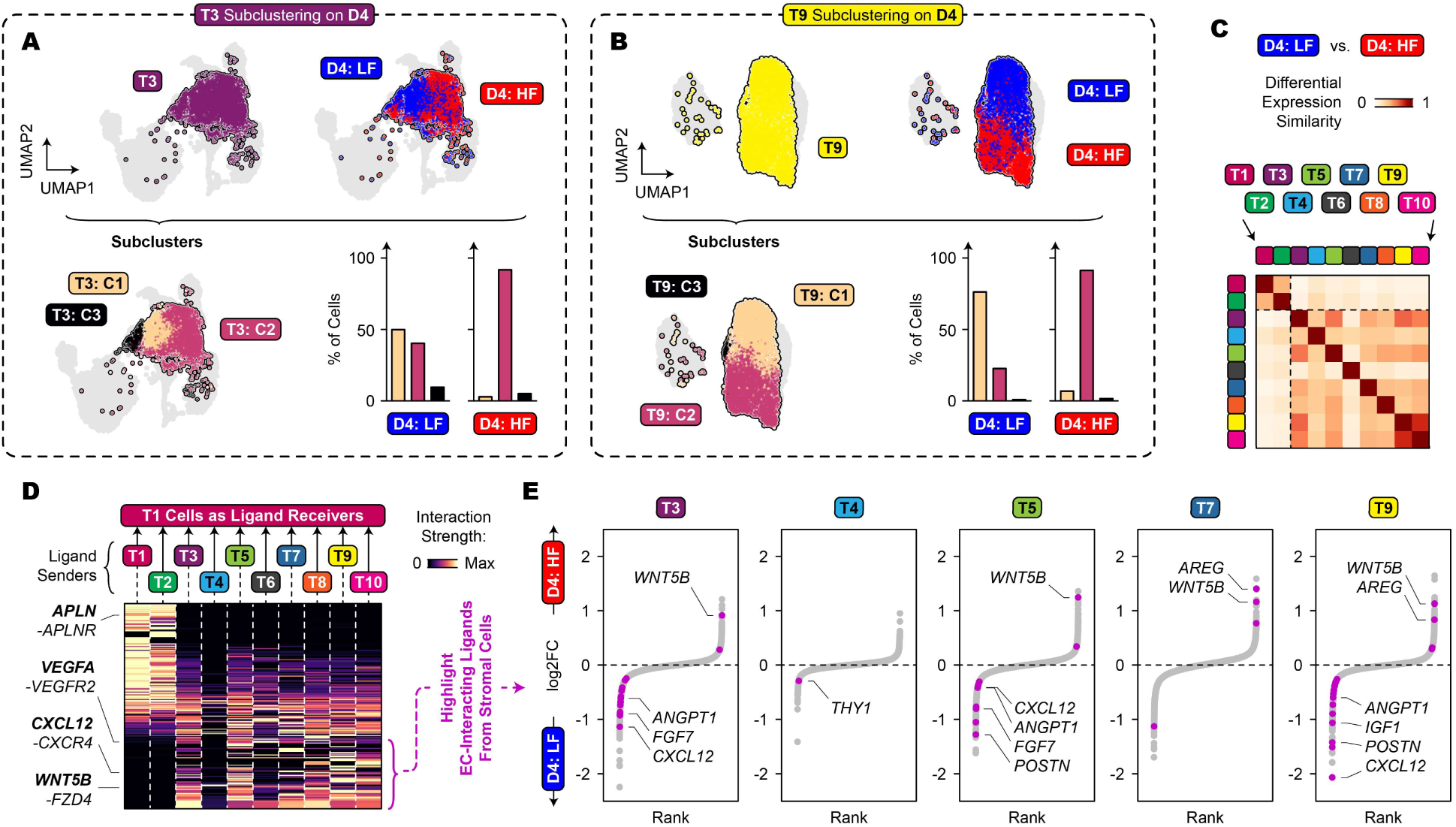
Transcriptomic differences in stromal cells between low flow and high flow. **(A)** Unsupervised sub-clustering of the T3 cell type on Day 4. **(B)** Unsupervised sub-clustering of the T9 cell type on Day 4. **(C)** Heatmap of cosine similarities between lists of differentially expressed genes (DEGs) for each cell type. DEGs were between Day 4 low flow and Day 4 high flow. **(D)** Cell-cell communication analysis using CellChat. The heatmap shows predicted interactions between each cell type (i.e. T1-T10) and the T1 endothelial cell type. Rows on the heatmap are ordered based on whether the interactions were stronger coming from endothelial cells (i.e. T1, T2) or coming from stromal cells (i.e. T3-T10). Interaction strengths are averages from the 3 conditions (i.e. Day 1, Day 4 low flow, and Day 4 high flow). **(E)** Log2 fold change rank plots of DEGs between Day 4 low flow and Day 4 high flow. Highlighted genes encode ligands primarily expressed in stromal cells and identified in panel (E) to interact with T1 endothelial cells.

**Extended Data Fig. 13.**
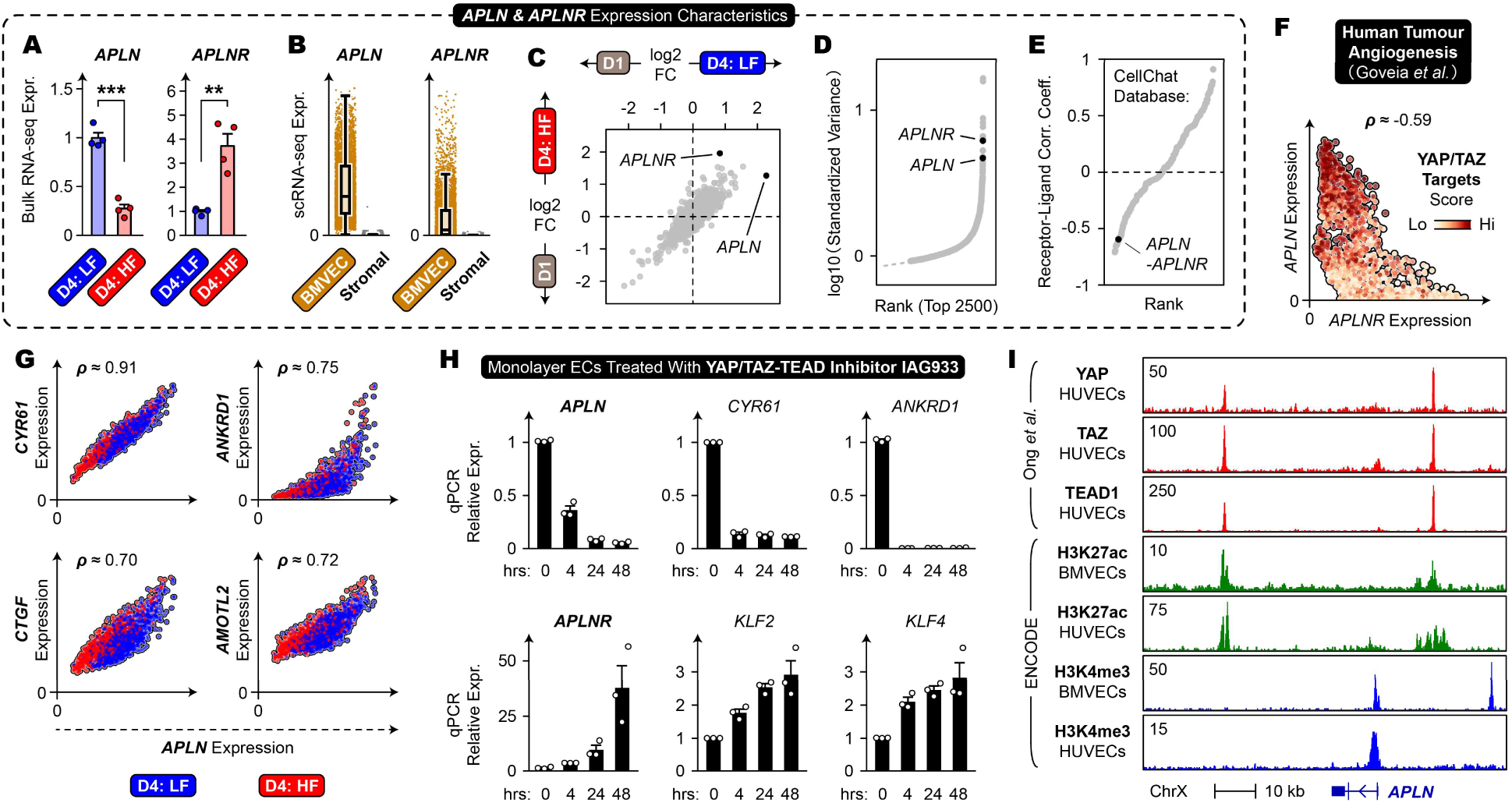
YAP/TAZ regulates the expressions of *APLN* and *APLNR*. **(A)** DESeq2-normalized counts of *APLN* and *APLNR* from bulk RNA-seq of vascular beds on Day 4 (4 devices per condition across 2 independent experiments). 2-tailed unpaired student’s t-test (** p < 0.01; *** p < 0.001). **(B)** scRNA-seq expression of *APLN* and *APLNR* from either endothelial cells (i.e. BMVECs) or stromal cells (i.e. NHLFs and HBVPs). Values are from all conditions combined. **(C)** Comparing differentially expressed genes between Day 1 and Day 4 for either Day 4 low flow or Day 4 high flow. **(D)** Rank plot of the most variably expressed genes. **(E)** Rank plot of receptor-ligand pairs from CellChat’s database showing the pair’s Spearman’s correlation coefficient. **(F)** Tip cell and stalk-like clusters from the human tumor angiogenesis atlas of Goveia *et al*. plotted for *APLN* and *APLNR* expressions ^33^. Gene set scoring for YAP/TAZ targets is shown. **(G)** Correlation of *APLN* with core YAP/TAZ target genes. Spearman’s correlation coefficients are shown. **(H)** qPCR gene expression of monolayer BMVECs treated with the YAP/TAZ-TEAD inhibitor IAG933 (2 µM; 3 independent experiments). **(I)** Re-analysis of endothelial ChIP-seq data for YAP, TAZ, and TEAD1 binding regions surrounding the *APLN* gene (Ong *et al*., ^19^; ENCODE, ^110^). **(B, E-G)** Expression values were denoised using Markov Affinity-based Graph Imputation of Cells. **(B, F, G)** Axes are log-normalized expression values. **(C-E, G)** Values are from T1 endothelial cells on Day 4. **(A, H)** Error bars show SEM.

**Extended Data Fig. 14.**
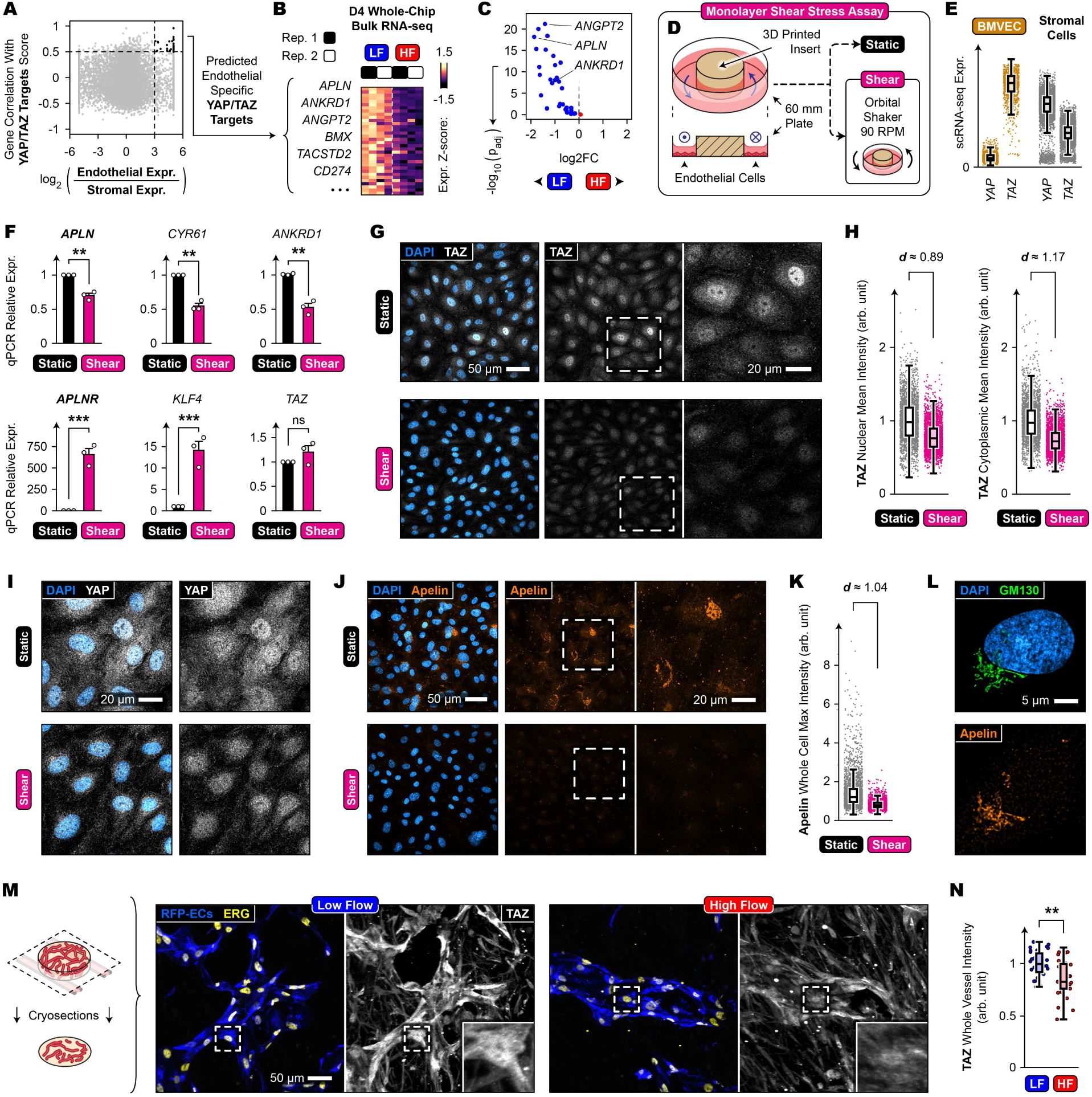
Fluid shear stress inhibits YAP/TAZ in endothelial cells. **(A)** Predicting endothelial-specific YAP/TAZ-regulated genes. Predicted genes correlate with gene set scoring of YAP/TAZ targets (Spearman’s correlation coefficient > 0.5) and are primarily expressed in endothelial cells (log2 fold change > 3). **(B)** Expression heatmap depicting bulk RNA-seq data of low or high flow vascular beds on Day 4. Predicted endothelial-specific YAP/TAZ-regulated genes are shown. **(C)** Differential expression volcano plot of data from (B). **(D)** Schematic for applying shear stress to monolayer endothelial cells by placing culture plates on an orbital shaker. **(E)** scRNA-seq expression of *YAP* and *TAZ* from either endothelial cells (i.e. BMVECs) or stromal cells (i.e. NHLFs and HBVPs). Values are from all conditions combined. **(F)** qPCR gene expression of monolayer BMVECs grown with and without shear stress (3 independent experiments; 2-tailed unpaired student’s t-test). Error bars show SEM. **(G, I, J)** BMVECs grown with and without shear stress for 4 days. Immunostainings for TAZ **(G)**, YAP **(I)**, and Apelin **(J)**. **(H)** Quantification of (G) (∼15k cells per condition across 3 independent experiments). **(K)** Quantification of (J) (∼8k cells per condition across 3 independent experiments). **(L)** Super-resolution imaging of a static condition cell from (J), stained for the Golgi marker GM130 and Apelin. **(M)** TAZ immunostaining of vascular bed cryosections from Day 4 using either low or high flow (ERG: endothelial nuclei). **(N)** Quantification of (M) (27-30 vessels and 8-9 devices per condition across 3 independent experiments; 2-tailed unpaired Welch’s t-test). **(H, K)** Cohen’s d effect size metric. **(F, N)** ns: not significant; * p < 0.05; ** p < 0.01; *** p < 0.001.

**Extended Data Fig. 15.**
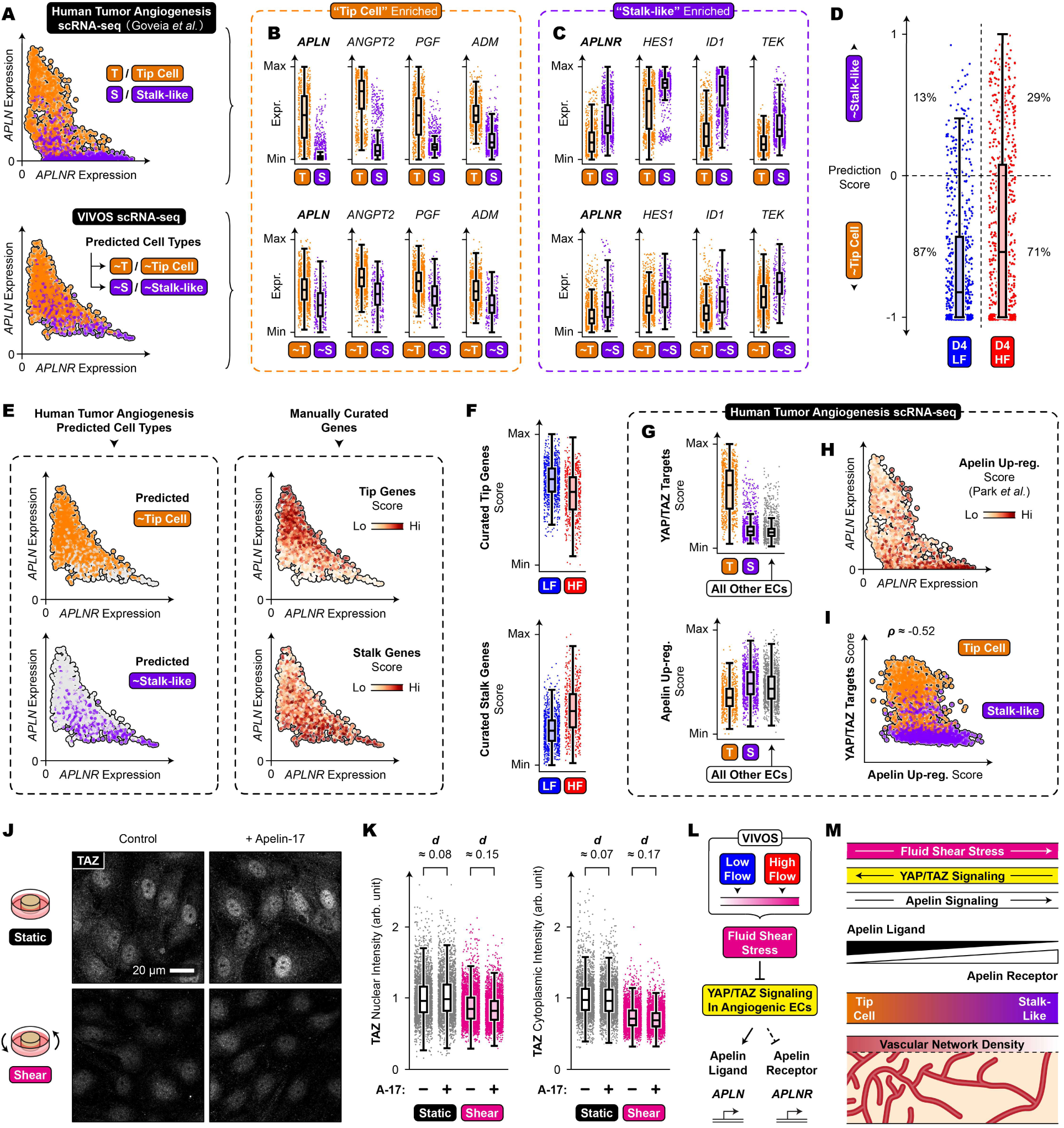
scRNA-seq characterization of tip and stalk cells on VIVOS. **(A)** Mapping tip cell and stalk-like clusters from the human tumor angiogenesis atlas of Goveia *et al*. onto T1 endothelial cells on Day 4 ^33^. **(B, C)** Up-regulated genes from tip cell and stalk-like human atlas clusters. Box plots are shown for both the human atlas (top row) and T1 endothelial cells on Day 4 (bottom row). **(D)** Prediction scores from (A). **(E)** Comparison between predicted cell types from (A) and gene set scoring using manually curated lists of tip cell and stalk cell genes. **(F)** Scores from (E). **(G)** Box plots depicting gene set scoring of the human atlas for YAP/TAZ targets and Apelin up-regulated genes (Park *et al*., ^57^). **(H-I)** Scatter plots depicting gene set scoring of the human atlas for YAP/TAZ targets and Apelin up-regulated genes (Park *et al*.), showing only tip cell and stalk-like human atlas clusters. **(J)** TAZ immunostaining of monolayer BMVECs grown with and without shear stress for 4 days, followed by addition of 10 µM Apelin-17 for 30 min. **(K)** Quantification of (J) (∼8k cells per condition across 3 independent experiments; Cohen’s d effect size metric). **(L)** Proposed model involving fluid shear stress inhibiting YAP/TAZ in angiogenic endothelial cells to modulate expressions of *APLN* and *APLNR*. **(M)** Proposed model involving the interplay between fluid shear stress, YAP/TAZ signaling, and Apelin signaling to regulate vascular network densities. **(A, B, C, E, H)** Axes show log-normalized expression values denoised using Markov Affinity-based Graph Imputation of Cells.

**Extended Data Fig. 16.**
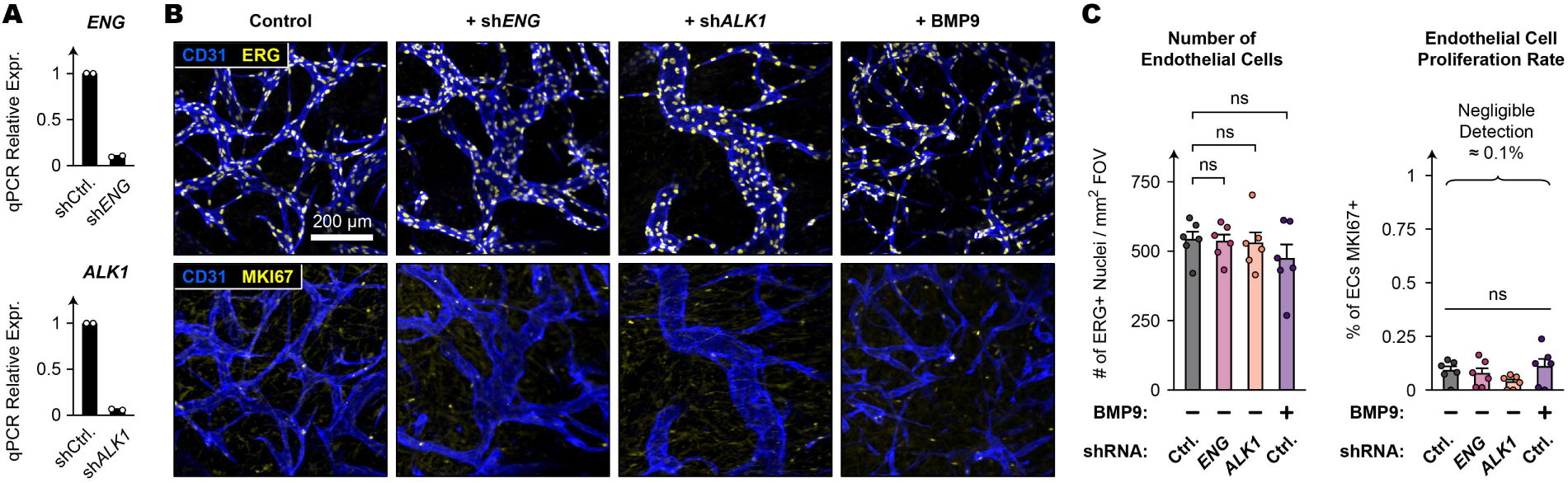
Assessing endothelial cell numbers and endothelial proliferation in vascular beds. **(A)** Evaluation of shRNA knockdown efficiency (2 independent experiments). **(B)** ERG and MKI67 immunostainings of vascular beds on Day 4 which were grown from shRNA-treated endothelial cells or additionally treated with BMP9 (10 ng/mL). **(C)** Quantifications of (B) (6 devices per condition across 2 independent experiments). 2-tailed unpaired student’s t-test (ns: not significant). **(A, C)** Error bars show SEM.

**Extended Data Fig. 17.**
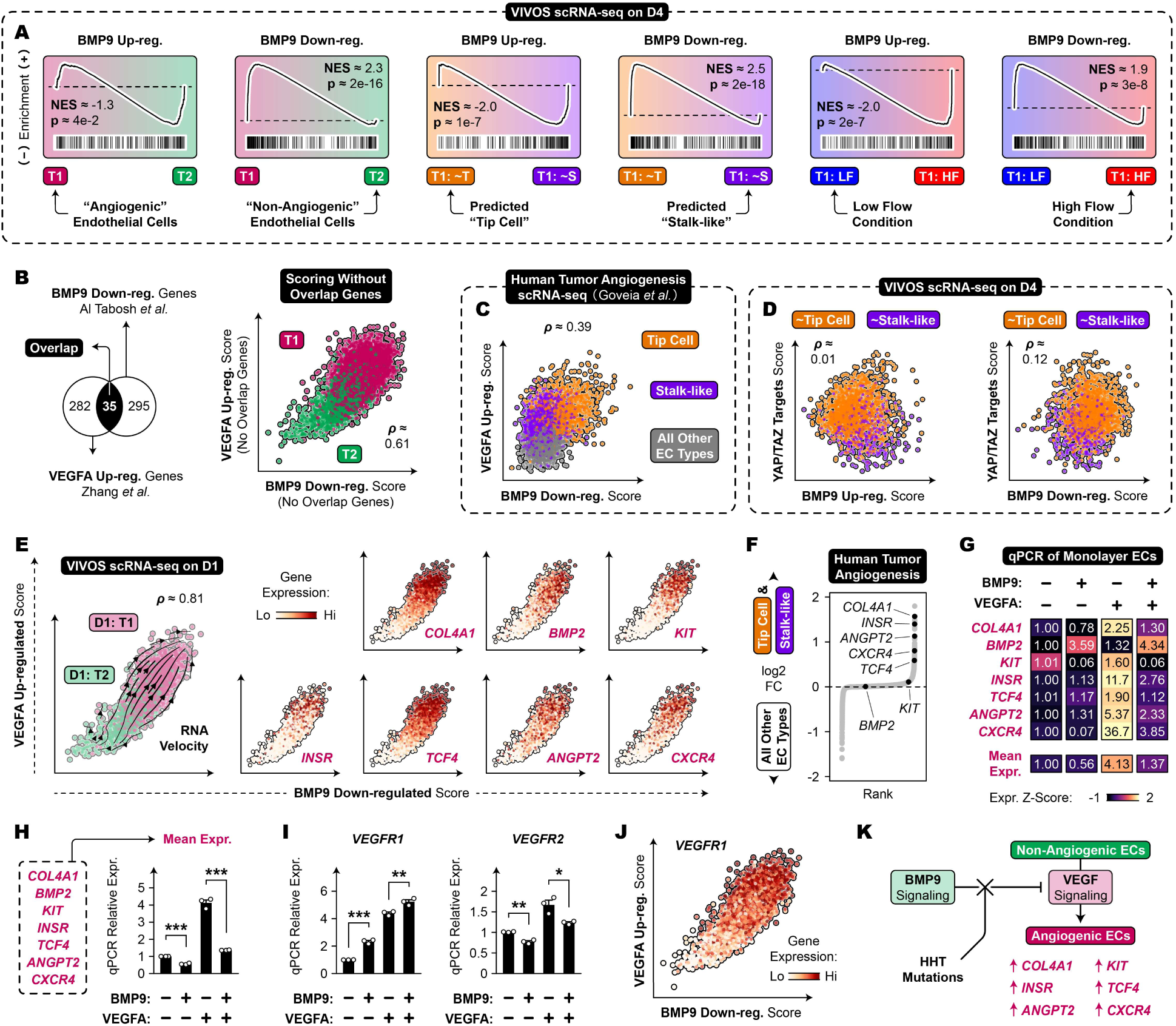
BMP9 antagonizes VEGFA-induced transcription. **(A)** Gene Set Enrichment Analysis of VIVOS scRNA-seq data for BMP9 up/down-regulated genes (Al Tabosh *et al.*, ^64^). **(B)** Left: Venn diagram between BMP9 down-regulated genes (Al Tabosh *et al.*) and VEGFA up-regulated genes (Zhang *et al.*, ^32^). Right: Gene set scoring of VIVOS scRNA-seq data for BMP9 down-regulated genes and VEGFA up-regulated genes, with overlapping genes removed. Endothelial cells from all conditions are shown. **(C)** Gene set scoring of the human tumor angiogenesis atlas from Goveia *et al.* ^33^ for BMP9 down-regulated genes and VEGFA up-regulated genes. **(D)** Gene set scoring of VIVOS scRNA-seq data for BMP9 up/down-regulated genes and YAP/TAZ targets. T1 endothelial cells on Day 4 are shown. **(E)** Gene set scoring of VIVOS scRNA-seq data for BMP9 down-regulated genes and VEGFA up-regulated genes. Endothelial cells from Day 1 are shown. RNA velocity streamlines (left). Individual gene expressions (right). **(F)** Rank plot of differentially expressed genes for tip cell and stalk-like human atlas clusters. **(G)** Heatmap of qPCR gene expression from monolayer BMVECs treated with BMP9 (10 ng/mL) and/or VEGFA (100 ng/mL) for 48 hrs. Numbers represent average relative expressions from 3 independent experiments. “Mean expression” refers to the geometric mean of the 7 individual genes. **(H)** Mean expression values from (G). **(I)** *VEGFR1* and *VEGFR2* expressions from the same experiment as (G). **(J)** *VEGFR1* expression in endothelial cells from all conditions. **(K)** Proposed model for antagonization of VEGFA-induced transcription by BMP9. **(B-E)** Spearman’s correlation coefficient. **(H-I)** 2-tailed unpaired student’s t-test (* p < 0.05; ** p < 0.01; *** p < 0.001). Error bars show SEM.

**Extended Data Table 1.** Curated tip cell and stalk cell marker genes.

| **Marker of** | **Gene** | **Reference** | **Reference #** |
| --- | --- | --- | --- |
| Tip Cells | *DLL4* | Suchting *et al.*; Hellstrom *et al.* | 55,123 |
| Tip Cells | *KDR* | Gerhardt *et al.* | 124 |
| Tip Cells | *CXCR4* | Strasser *et al.*; Crouch *et al.* | 54,125 |
| Tip Cells | *ANGPT2* | del Toro *et al.*; Crouch *et al.* | 17,54 |
| Tip Cells | *APLN* | del Toro *et al.*; Crouch *et al.* | 17,54 |
| Tip Cells | *ESM1* | del Toro *et al.* | 17 |
| Tip Cells | *ADM* | Crouch *et al.* | 54 |
| Tip Cells | *PGF* | Crouch *et al.* | 54 |
| Tip Cells | *NRP1* | Fantin *et al.* | 126 |
| Tip Cells | *PDGFB* | Gerhardt *et al.*; Suchting *et al.* | 123,124 |
| Stalk Cells | *JAG1* | Benedito *et al.* | 127 |
| Stalk Cells | *FLT1* | Jakobsson *et al.* | 58 |
| Stalk Cells | *TEK* | del Toro *et al.* | 17 |
| Stalk Cells | *HEY1* | Larrivée *et al.*; Moya *et al.* | 52,53 |
| Stalk Cells | *HES1* | Larrivée *et al.*; Moya *et al.* | 52,53 |
| Stalk Cells | *SMAD6* | Larrivée *et al.*; Moya *et al.* | 52,53 |
| Stalk Cells | *ID1* | Larrivée *et al.*; Moya *et al.* | 52,53 |
| Stalk Cells | *MKI67* | Gerhardt *et al.* | 124 |
| Stalk Cells | *PCNA* | Gerhardt *et al.* | 124 |

**Extended Data Table 2.**
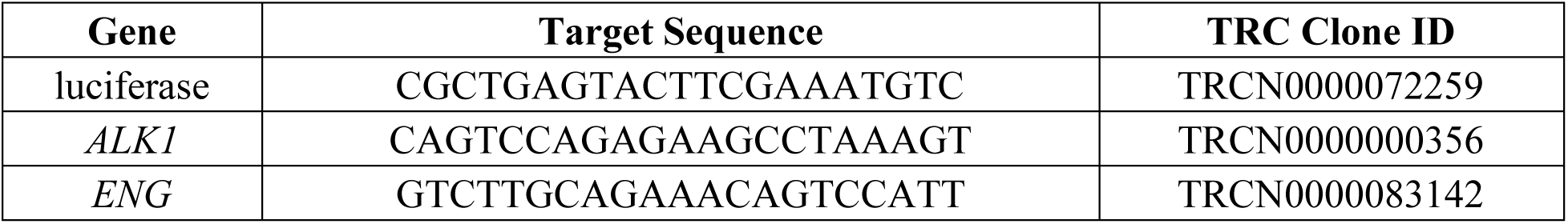
shRNA sequences.

**Extended Data Table 3.**
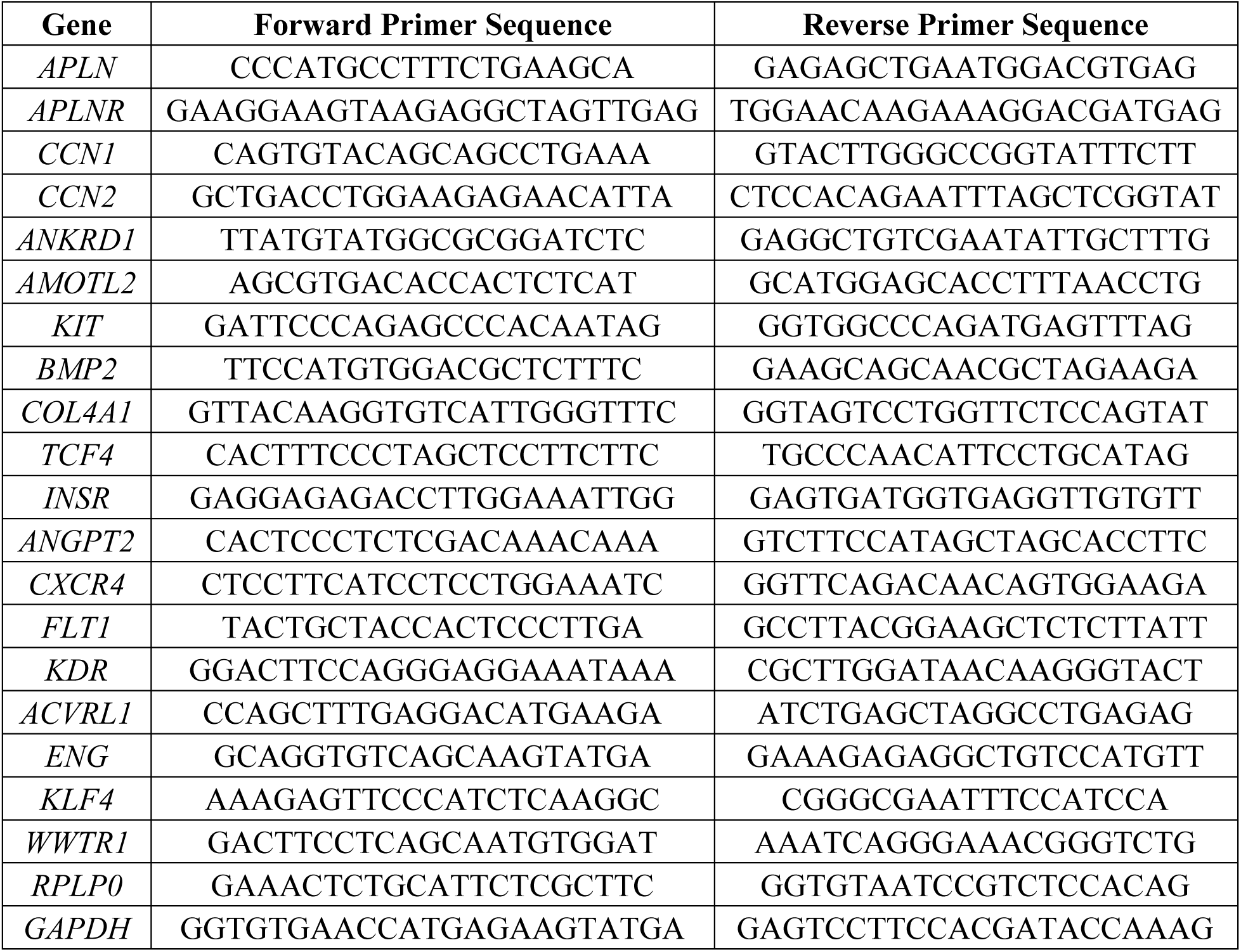
qPCR primer sequences. Aliases: *CCN1*/*CYR61*, *CCN2*/*CTGF*, *FLT1*/*VEGFR1*, *KDR*/*VEGFR2*, *ACVRL1*/*ALK1*, *WWTR1*/*TAZ*.

**Extended Data Table 4.**
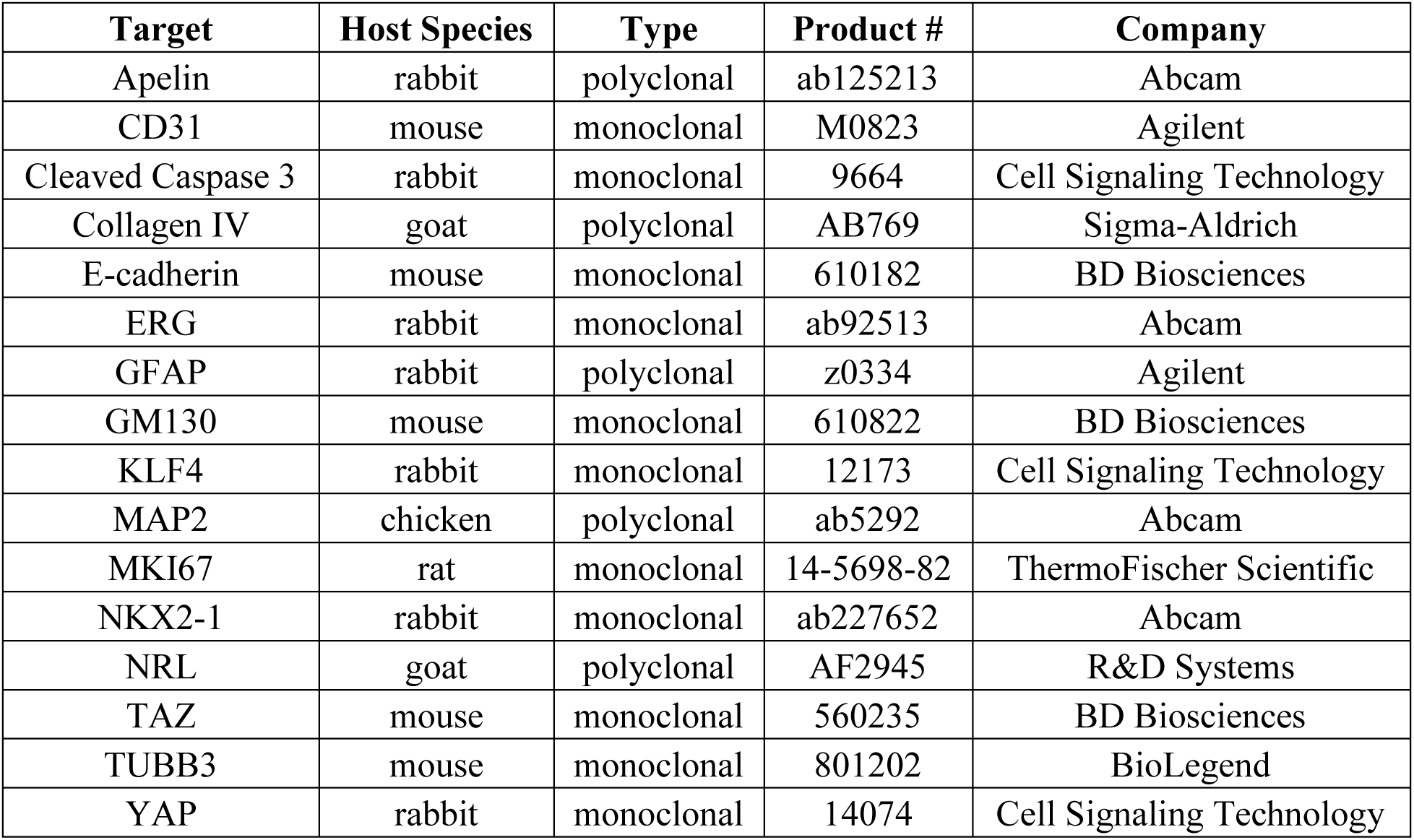
Antibody information.

| **Target** | **Host Species** | **Type** | **Product #** | **Company** |
| --- | --- | --- | --- | --- |
| Apelin | rabbit | polyclonal | ab125213 | Abcam |
| CD31 | mouse | monoclonal | M0823 | Agilent |
| Cleaved Caspase 3 | rabbit | monoclonal | 9664 | Cell Signaling Technology |
| Collagen IV | goat | polyclonal | AB769 | Sigma-Aldrich |
| E-cadherin | mouse | monoclonal | 610182 | BD Biosciences |
| ERG | rabbit | monoclonal | ab92513 | Abcam |
| GFAP | rabbit | polyclonal | z0334 | Agilent |
| GM130 | mouse | monoclonal | 610822 | BD Biosciences |
| KLF4 | rabbit | monoclonal | 12173 | Cell Signaling Technology |
| MAP2 | chicken | polyclonal | ab5292 | Abcam |
| MKI67 | rat | monoclonal | 14-5698-82 | ThermoFischer Scientific |
| NKX2-1 | rabbit | monoclonal | ab227652 | Abcam |
| NRL | goat | polyclonal | AF2945 | R&D Systems |
| TAZ | mouse | monoclonal | 560235 | BD Biosciences |
| TUBB3 | mouse | monoclonal | 801202 | BioLegend |
| YAP | rabbit | monoclonal | 14074 | Cell Signaling Technology |

## Notes

https://doi.org/10.5281/zenodo.18529054

https://www.ncbi.nlm.nih.gov/geo/query/acc.cgi?acc=GSE318597

### Competing Interest Statement

T.H.Z.J., A.A.S., L.P., J.L.W, and L.A have submitted US provisional and PCT patent applications related to VIVOS, the platform described in this manuscript.

