## Supplementary material for "Physiological perfusion of human vasculature reveals a YAP/TAZ-Apelin switch linking intraluminal flow to endothelial state transitions and vessel remodeling": VIVOS_Supplementary_Information_Guide_biorxiv.pdf

### Supplementary Methods

Derivation of formulas for the vessel permeability assay and tracking of fluorescent beads.

**Supplementary Video 1. Fluorescent bead perfusion of vascular beds.**

Addition of 1 μm diameter fluorescent beads to the impeller pump's nutrient media reservoir and tracking of bead movement at multiple locations within the microfluidic chip.

**Supplementary Video 2. Dynamic waveforms from the impeller pump.**

Various flow sensor readings over time for the impeller pump.

**Supplementary Video 3. Fluorescent dextran vessel permeability assay.**

Working principle of the fluorescent dextran assay which measures vessel wall permeabilities and vessel volume fractions.

**Supplementary Video 4. Particle image velocimetry analysis of fluorescent bead perfusion.**

Working principle of the fluorescent bead perfusion assay which applies particle image velocimetry to measure vessel flowrates, vessel flow velocities, and vessel wall shear stresses.

**Supplementary Video 5. Fluorescent bead perfusion of low flow and high flow experimental conditions.**

Example fluorescent bead perfusion time-lapse sequences from Day 4 using either low flow (300 RPM impeller speed) or high flow (900 RPM impeller speed).

**Supplementary Video 6. Fluorescent bead perfusion of *ENG*-knockdown and *ALK1*-knockdown vascular beds.**

Example fluorescent bead perfusion time-lapse sequences from *ENG*-knockdown and *ALK1*-knockdown vascular beds on Day 4.

**Supplementary Data 1. scRNA-seq cell type marker genes.**

Output data from "FindAllMarkers()" in Seurat following SCTransform v2 normalization. "FindAllMarkers()" was applied separately to cells originating from each cell line (i.e. BMVECs, NHLFs, and HBVPs), as well as separately to cells from Day 1, Day 4, or all conditions together (i.e. Day 1 and Day 4).

**Supplementary Data 2. Gene sets used for scRNA-seq scoring.**

VEGFA up/down-regulated gene sets are derived from Zhang *et al*. [32]. The Apelin up-regulated gene set is derived from Park *et al*. [57]. BMP9 up/down-regulated gene sets are derived from Al Tabosh *et al*. [64]. Detailed steps for deriving these gene sets can be found in the methods section.

**Supplementary Data 3. Differentially expressed genes between Day 1 and Day 4.**

Output data from "FindMarkers()" in Seurat following SCTransform v2 normalization. For each cell type (i.e. T1-T10), differentially expressed genes were calculated separately for Day 1 vs. Day

4 low flow, and Day 1 vs. Day 4 high flow. The “MERGED_LF_HF_avg_log2FC” column refers to the arithmetic mean of the log2 fold changes from the two separate calculations. The “MERGED_LF_HF_p_val_adj” column refers to the geometric mean between the adjusted p-values from the two separate calculations. A positive log2 fold change refers to greater expression on Day 4 while a negative log2 fold change refers to greater expression on Day 1.

**Supplementary Data 4. Differentially expressed genes between Day 4 low flow and Day 4 high flow.**

Output data from “FindMarkers()” in Seurat following SCTransform v2 normalization. For each cell type (i.e. T1-T10), differentially expressed genes were calculated between Day 4 low flow and Day 4 high flow. A positive log2 fold change refers to greater expression for Day 4 high flow while a negative log2 fold change refers to greater expression for Day 4 low flow.

**Supplementary Data 5. Bulk RNA-seq and scRNA-seq mapping metrics.**

Bulk RNA-seq metrics are from STAR. scRNA-seq metrics are from CellRanger.

**Supplementary Data 6. Marker genes of excluded endothelial clusters during scRNA-seq quality control.**

Output data from “FindMarkers()” in Seurat following SCTransform v2 normalization. Excluded cluster 1 was assigned to be low quality cells. Excluded cluster 2 was assigned to be endothelial-stromal doublets. Additional information can be found in the methods section.

**Supplementary Data 7. *APLN* co-expression analysis.**

A list of all genes detected by scRNA-seq and their Spearman’s correlation coefficient with *APLN*. Correlation coefficients were calculated using T1 endothelial cells on Day 4. MAGIC-denoised expression values were used (Markov Affinity-based Graph Imputation of Cells).
