## Supplementary material for "Physiological perfusion of human vasculature reveals a YAP/TAZ-Apelin switch linking intraluminal flow to endothelial state transitions and vessel remodeling": VIVOS_Supplementary_Methods_biorxiv.pdf

<u>Derivation of Formulas for the Vessel Permeability Assay</u>

To calculate vessel wall permeability, we applied lumped system analysis and simplified the general formula for compound flux ( $J$ ) across a membrane (i.e. the vessel wall) as a function of a global permeability coefficient ( $P$ ), a global concentration difference ( $\Delta C$ ), and total vessel surface area ( $S_V$ ).

$$J = \int_S (\Delta C \cdot P)\, dS = \Delta C \cdot PS_V$$

The measured fluorescent intensity ( $I_m$ ) was assumed to correlate linearly with compound concentration ( $C$ ) using a conversion factor ( $k_C$ ) plus a background intensity value ( $I_{bg}$ ):

$$I_m(t) = k_C \cdot C(t) + I_{bg}$$

For modeling vessel compound concentration ( $C_V(t)$ ) with exponential dynamics:

$$C_V(t) = k_{0,V} \cdot e^{-k_V t} + k_{r,V} \qquad \frac{d}{dt} C_V(t) = -k_V \cdot k_{0,V} \cdot e^{-k_V t}$$

Expressed in terms of regression coefficients from applying nonlinear least squares curve fitting to measured fluorescent intensities in R:

$$I_{m,V}(t) = A_V \cdot e^{-\lambda_V t} + B_V$$

$$A_V = k_C \cdot k_{0,V} \qquad B_V = k_C \cdot k_{r,V} + I_{bg} \qquad \lambda_V = k_V$$

For modeling interstitial compound concentration ( $C_I(t)$ ) with exponential dynamics:

$$C_I(t) = k_{\infty,I} \cdot \left(1 - e^{-k_I t}\right) + k_{r,I} \qquad \frac{d}{dt} C_I(t) = k_I \cdot k_{\infty,I} \cdot e^{-k_I t}$$

Expressed in terms of regression coefficients from applying nonlinear least squares curve fitting to measured fluorescent intensities in R:

$$I_{m,I}(t) = A_I \cdot \left(1 - e^{-\lambda_I t}\right) + B_I$$

$$A_I = k_C \cdot k_{\infty,I} \qquad B_I = k_C \cdot k_{r,I} + I_{bg} \qquad \lambda_I = k_I$$

Start with the mass balance of compounds crossing the vessel wall:

$$\frac{d}{dt} N_I(t) = -\frac{d}{dt} N_V(t)$$

Substituting $N = CV$:

$$V_I \cdot \frac{d}{dt} C_I(t) = -V_V \cdot \frac{d}{dt} C_V(t)$$

$$V_I \cdot \left(k_I \cdot k_{\infty,I} \cdot e^{-k_I t}\right) = -V_V \cdot \left(-k_V \cdot k_{0,V} \cdot e^{-k_V t}\right)$$

$k_V = k_I$ because compound movement occurs on the same timescale for the vessel and interstitial regions:

$$V_I \cdot k_{\infty,I} = V_V \cdot k_{0,V}$$

Expressing $V_I$ & $V_V$ in terms of a vessel volume fraction ( $\alpha$) and total vascular bed compartment volume ( $V_{\text{total}}$ ):

$$V_I = (1-\alpha) \cdot V_{\text{total}} \qquad V_V = \alpha \cdot V_{\text{total}}$$

Substituting in regression coefficients and vessel volume fraction ( $\alpha$):

$$(1-\alpha) \cdot V_{\text{total}} \cdot \frac{A_I}{k_C} = \alpha \cdot V_{\text{total}} \cdot \frac{A_V}{k_C}$$

$$\alpha = \frac{A_I}{A_V + A_I}$$

Next, solve for the compound concentration difference between the vessels and the interstitial space ( $\Delta C$ ):

$$\Delta C(t) = C_V(t) - C_I(t)$$

$$\Delta C(t) = k_{0,V} \cdot e^{-k_V t} + k_{\infty,I} \cdot e^{-k_I t} + k_{r,V} - k_{\infty,I} - k_{r,I}$$

$k_{r,V} = k_{\infty,I} + k_{r,I}$ because the compound concentration inside the vessels and initial space are equal as $t \to \infty$:

$$\Delta C(t) = k_{0,V} \cdot e^{-k_V t} + k_{\infty,I} \cdot e^{-k_I t}$$

$$\Delta C(t) = \frac{A_V}{k_C} \cdot e^{-\lambda_V t} + \frac{A_I}{k_C} \cdot e^{-\lambda_I t}$$

Next, start with the definition of solute flux ( $J$ ). Note that using $J_V(t)$ yields the same result.

$$J(t) = \frac{d}{dt} N$$

$$J_I(t) = V_I \cdot \frac{d}{dt} C_I$$

$$J_I(t) = V_I \cdot k_I \cdot k_{\infty,I} \cdot e^{-k_I t}$$

$$J_I(t) = (1-\alpha) \cdot V_{\text{total}} \cdot \lambda_I \cdot \frac{A_I}{k_C} \cdot e^{-\lambda_I t}$$

Substituting in:

$$J = \Delta C \cdot PS_V$$

$$(1-\alpha) \cdot V_{\text{total}} \cdot \lambda_I \cdot \frac{A_I}{k_C} \cdot e^{-\lambda_I t} = \left( \frac{A_V}{k_C} \cdot e^{-\lambda_V t} + \frac{A_I}{k_C} \cdot e^{-\lambda_I t} \right) \cdot PS_V$$

Let $\lambda_I = \lambda_V = \lambda$ and substitute in vessel volume fraction ( $\alpha$ ):

$$\frac{A_V}{A_I + A_V} \cdot V_{\text{total}} \cdot \lambda \cdot A_I = (A_V + A_I) \cdot PS_V$$

$$P = \frac{V_{\text{total}}}{S_V} \cdot \lambda \cdot \frac{A_V A_I}{(A_V + A_I)^2}$$

To approximate the ratio $V_{\text{total}}/S_V$, we assumed that the vasculature was one single long cylinder with diameter equal to the average vessel diameter ( $\overline{d_V}$ ).

$$V_V = \alpha \cdot V_{\text{total}} = \frac{\pi \cdot \overline{d_V}^2 \cdot l}{4} \qquad S_V = \pi \cdot \overline{d_V} \cdot l$$

Substituting in:

$$\frac{V_{\text{total}}}{S_V} = \left(\frac{\pi \cdot \overline{d_V}^2 \cdot l}{4}\right)\left(\frac{A_I + A_V}{A_I}\right)\left(\frac{1}{\pi \cdot \overline{d_V} \cdot l}\right)$$

$$\frac{V_{\text{total}}}{S_V} = \frac{\overline{d_V}}{4}\left(\frac{A_I + A_V}{A_I}\right)$$

Solving for the vessel permeability coefficient ( $P$ ):

$$P = \lambda \cdot \frac{A_V A_I}{(A_V + A_I)^2} \cdot \frac{\overline{d_V}}{4} \cdot \left(\frac{A_I + A_V}{A_I}\right)$$

$$P = \frac{\overline{d_V} \cdot \lambda}{4} \frac{A_V}{A_V + A_I}$$

$$P = \frac{\overline{d_V} \lambda (1 - \alpha)}{4}$$

Derivation of Formulas for Tracking of Fluorescent Beads

The input data to the following analysis is a fast time-lapse sequence (i.e. 500 FPS) of fluorescent beads flowing through vessels. Frames ( $F_n$ ) from each time-lapse sequence were first split into multiple groups using the following pattern, followed by calculating the maximum intensity projection ( $M_n$ ) of all frames per group. We used 25 groups in our analyses, which we found to be the most robust. For illustrative purposes, we show 10 groups.

$$M_1 = \mathrm{MaxIP}(F_1, F_{11}, F_{21}, F_{31}, F_{41}, \cdots)$$

$$M_2 = \mathrm{MaxIP}(F_2, F_{12}, F_{22}, F_{32}, F_{42}, \cdots)$$

$$M_3 = \mathrm{MaxIP}(F_3, F_{13}, F_{23}, F_{33}, F_{43}, \cdots)$$

...

$$M_{10} = \mathrm{MaxIP}(F_{10}, F_{20}, F_{30}, F_{40}, F_{50}, \cdots)$$

After adding random noise to non-vessel regions, pairs of adjacent maximum intensity projections were processed using particle image velocimetry (PIV) using OpenPIV to obtain velocity vector fields ( $\vec{u}$ ). The random noise ensured OpenPIV did not miscalculate non-vessel regions where no fluorescent signal was present.

$$\vec{u}_{1,2} = \mathrm{PIV}(M_1 \Longleftrightarrow M_2)$$

$$\vec{u}_{3,4} = \mathrm{PIV}(M_3 \Longleftrightarrow M_4)$$

$$\vec{u}_{5,6} = \mathrm{PIV}(M_5 \Longleftrightarrow M_6)$$

...

To reduce noise, the median of all calculated velocity vector fields was utilized for the final velocity vector field result.

$$\vec{u} = \mathrm{Median}(\vec{u}_{1,2}, \vec{u}_{3,4}, \vec{u}_{5,6}, \cdots)$$

The 2D velocity gradient tensor ( $\mathbf{L}$ ), which was utilized because of the 2D implementation of particle image velocimetry (PIV), can be further decomposed into symmetric ( $\mathbf{E}$ ) and skew symmetric components ( $\mathbf{W}$ ):

$$\mathbf{L} = \nabla \vec{u} = \begin{bmatrix} \frac{\partial u_x}{\partial x} & \frac{\partial u_x}{\partial y} \\ \frac{\partial u_y}{\partial x} & \frac{\partial u_y}{\partial y} \end{bmatrix} = \frac{1}{2}(\mathbf{L} + \mathbf{L}^T) + \frac{1}{2}(\mathbf{L} - \mathbf{L}^T) = \mathbf{E} + \mathbf{W}$$

The general form of a stress tensor for a Newtonian fluid, where $\mathbf{I}$ is the identity matrix,

$$\sigma = -P\mathbf{I} + 2\mu\mathbf{E} + \lambda\mathbf{I}\nabla \cdot \vec{u}$$

was simplified to,

$$\sigma = 2\mu\mathbf{E}$$

by ignoring the hydrostatic pressure term and assuming zero divergence. Time-lapse sequences were taken only of vessels parallel to the imaging plane.

Given the traction vector ( $t$ ),

$$t = \sigma \cdot \hat{n}$$

we construct a normal unit vector ( $\hat{n}$ ) by rotating the velocity field unit vector ( $\hat{u}$ ) by 90 degrees. Although this rotated vector won't always point towards the fluid at the vessel wall, we later take the magnitude of the wall shear stress.

$$\hat{n} = \begin{bmatrix} 0 & -1 \\ 1 & 0 \end{bmatrix} \cdot \hat{u} = \begin{bmatrix} -\hat{u}_y \\ \hat{u}_x \end{bmatrix}$$

Wall shear stress ( $\tau_{\mathrm{WSS}}$ ) is the magnitude of the traction vector ( $t$ ) tangential to the vessel wall surface and flow direction:

$$\tau_{\mathrm{WSS}} = \left|(\sigma \cdot \hat{n}) \cdot \hat{u}\right|$$

$$\tau_{\text{WSS}} = \mu \left| \left( \left( \nabla \vec{u} + (\nabla \vec{u})^T \right) \cdot \hat{n} \right) \cdot \hat{u} \right|$$

$$\tau_{\text{WSS}} = \mu \left| \left( \begin{bmatrix} 2\dfrac{\partial u_x}{\partial x} & \dfrac{\partial u_x}{\partial y} + \dfrac{\partial u_y}{\partial x} \\ \dfrac{\partial u_x}{\partial y} + \dfrac{\partial u_y}{\partial x} & 2\dfrac{\partial u_y}{\partial y} \end{bmatrix} \cdot \begin{bmatrix} -\hat{u}_y \\ \hat{u}_x \end{bmatrix} \right) \cdot \begin{bmatrix} \hat{u}_x \\ \hat{u}_y \end{bmatrix} \right|$$

$$\tau_{\text{WSS}} = \mu \left| \left( 2\frac{\partial u_x}{\partial x} \right) (-\hat{u}_x \hat{u}_y) + \left( \frac{\partial u_x}{\partial y} + \frac{\partial u_y}{\partial x} \right) (\hat{u}_x^2 - \hat{u}_y^2) + \left( 2\frac{\partial u_y}{\partial y} \right) (\hat{u}_x \hat{u}_y) \right|$$

Note that this general formula collapses down to the expected formula for the simple case of uniform flow past a wall:

$$\hat{u} = \begin{bmatrix} \hat{u}_x \\ \hat{u}_y \end{bmatrix} = \begin{bmatrix} 1 \\ 0 \end{bmatrix} \quad \Longrightarrow \quad \tau_{\text{WSS}} = \mu \left| \left( \frac{\partial u_x}{\partial y} + \frac{\partial u_y}{\partial x} \right) (1^2) \right| = \mu \frac{\partial u_x}{\partial y}$$
